# Temporal Transcriptome and Promoter Architecture of the African Swine Fever Virus

**DOI:** 10.1101/847343

**Authors:** Gwenny Cackett, Dorota Matelska, Michal Sýkora, Raquel Portugal, Michal Malecki, Jürg Bähler, Linda Dixon, Finn Werner

**Affiliations:** Institute for Structural and Molecular Biology, Darwin Building, University College London, Gower Street, London WC1E 6BT, United Kingdom; Pirbright Institute, UK; Charles University, Prague, CZ; Department of Genetics, Evolution & Environment and Institute of Healthy Ageing, Gower Street, University College London, London WC1E 6BT, UK; Institute of Genetics and Biotechnology, Faculty of Biology, University of Warsaw, Warsaw, Poland

## Abstract

The African Swine Fever Virus causes haemorrhagic fever in domestic pigs and presents the biggest global threat to animal farming in recorded history. Despite its importance, very little is known the mechanisms and temporal regulation of transcription in ASFV. Here we report the first detailed viral transcriptome analysis of ASFV during early and late infection of *Vero* cells. In addition to total RNA sequencing, we have characterised the transcription start sites and transcription termination sites at nucleotide-resolution, revealing the distinct DNA consensus motifs of early and late promoters, as well as the sequence determinants for transcription termination. ASFV can utilise alternative promoters to generate distinct proteins from the same transcription unit that differ with respect to the polypeptide N-terminus. Finally, our results reveal that the ASFV-RNAP undergoes transcript slippage at the 5’ end of transcription units that in a promoter sequence-specific manner results in the addition of 5’-AT and 5’-ATAT tails to mRNAs.

## Introduction

African swine fever virus (ASFV) causes incurable and lethal haemorrhagic fever in domestic pigs. In 2019, ASF presents an acute and global animal health emergency that has the potential to devastate entire national economies since effective vaccines or antiviral drugs are not available (FAO of UN). Urgent action is needed to advance our knowledge about the fundamental biology of ASFV, including the mechanisms and temporal regulation of transcription. A thorough understand of RNAP and factor function, as well as accurate knowledge of which genes are expressed and their amino acid sequence, is direly needed for the development of antiviral drugs and vaccines, respectively. ASFV is the sole member of *Asfarviridae* ^1^, a family resembling others in the Nucleocytoplasmic Large DNA Viruses (NCLDV) ^2, 3^. ASFV originated in East Sub-Saharan Africa where it remains endemic, though transcontinental spread to in Georgia 2007 ^4^ and subsequent spread in Europe and to Asia 2018 ^5^ has resulted in the current emergency situation. ASFV has a linear double-stranded DNA (dsDNA) genome of ∼170–194 kbp encoding ∼150–170 open reading frames (ORFs). The genomic variation between strains predominantly originates from loss or gain of genes at the genome termini that belong to multigene families (MGFs) ^6^. Despite its global economic importance, little is known about ASFV transcription, even though it is believed to be similar to the vaccinia virus (VACV) system^7–9^, a distantly-related NCLDV member of the *Poxviridae* family ^10^. ASFV encodes both an RNA polymerase (RNAP), a poly-A polymerase, and an mRNA capping enzyme, and extracts obtained from mature virus particles are fully transcription competent ^9, 11, 12^. The basal ASFV transcription machinery resembles the eukaryotic RNAPII system by encompassing an (8-subunit) ASFV-RNAP and distant relatives of the TATA-binding protein (TBP), the transcription initiation factor II B (TFIIB) and the elongation factor TFIIS. ASFV also encodes a histone-like DNA binding protein, pA104R^13^. Of particular interest is the possibility that the ASFV-RNAP can gain promoter-specificity in terms of temporal (early or late) gene expression dependent on the association with either host (eukaryote)-like TBP/TFIIB or virus-specific factors such as D1133L/G1340L, which are homologous to the VACV D6-A7 ETF heterodimer ^14, 15^. The promoter consensus motifs for early and late ASFV genes have not been characterised on a genome-wide scale, or in fact in any detail, with the exception of an AT-rich sequence motif upstream of the p72 transcription start site (TSS) and a consistently AT-rich region overlapping the TSS in late genes ^16^. Importantly, information about the temporal ASFV gene expression, TSS and transcription termination sites (TTS) is sparse ^9, 10^.

We have focused our analysis on the BA71V strain (170,101 bp genome, with 153 annotated ORFs ^17, 18^), because this is the best studied ASFV strain with regards to viral transcription and mRNA modification and protein expression ^9, 19^. Here we have applied a combination of transcriptome sequencing (RNA-seq), RNA 5’-end (<u>c</u>ap <u>a</u>nalysis <u>g</u>ene <u>e</u>xpression sequencing or ‘CAGE-seq’)- and RNA 3’-end (3’ RNA-seq) determination. We report (i) a map of the ASFV transcriptome at one early and one late time point during infection (5 h and 16 h post-infection), in (Figure 1)), (ii) a genome-wide TSS map that has allowed us to define early and late ASFV promoter consensus motifs as well as shed light on 5’ UTRs and the phenomenon of RNA-5’ leaders in ASFV, and (iii) a genome-wide TTS map that provides novel insights into the mechanism of transcription termination in ASFV and by inference in NCLDVs.

**Fig. 1.**
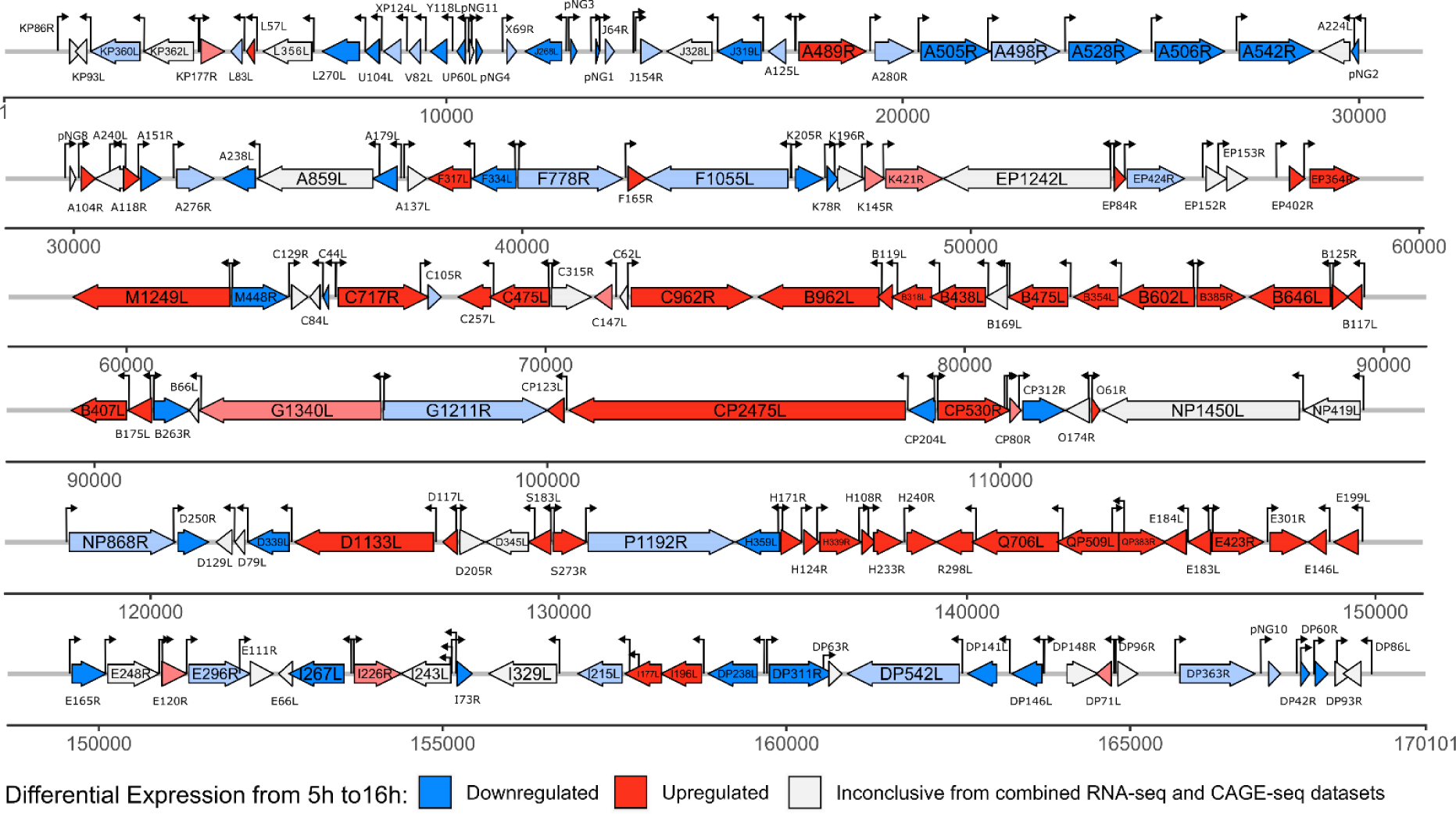
Annotated genome of ASFV-BA71V indicating transcription start sites (TSS) and early and late genes. The map includes 153 previously annotated as well as novel genes identified in this study and their differential expression pattern. Downregulated-(blue) and upregulated (red) genes were differentially expressed according to both RNA-seq and CAGE-seq data, while colour coding in light-blue and light red showed the same type of differential expression in the both techniques but only statistically significant (adjusted *p*-value < 0.05) according to either CAGE-seq or RNA-seq, respectively. The map was visualised with the R package gggenes.

## Results

### ASFV transcriptome and genome structure

A transcriptome is defined by the overall expression levels of mRNAs, and their 5’ and 3’ termini. We carried out RNA-seq, CAGE-seq and 3’ RNA-seq in order to characterise these parameters during early and late ASFV infection, which combined inform about the ASFV transcriptome and DNA sequence signatures associated with transcription initiation and termination.

*Vero* cells were infected with BA71V, and at 5h and 16h post-infection RNA was extracted from cells. Bowtie 2 ^20^ mapping of the RNA-seq, CAGE-seq and 3’ RNA-seq reads (summarised in Supplementary Table 1) showed a strong correlation between replicates (Pearson correlation coefficient *r* ≥ 0.98), with one exception of RNA-seq from 16h (*r* of 0.74 and 0.84 for two strands, Supplementary Figure 1). For both time points, reads mapped well to the ASFV genome and were consistent between replicates for all three techniques. A genome-wide map of reads mapped from all three NGS techniques is shown in Figure 2a, while examples of signals for transcription start sites (TSSs) and transcription termination sites (TTSs) revealed from downstream analysis are shown in Figure 2. b-e. The RNA-seq sequencing depth was sufficient to analyse significant changes in ASFV gene transcription (i.e. reads) at early and late infection due the small genome size (170 kb), and unsurprisingly most CAGE-seq reads were aligned upstream of ORFs start codons. However, a subset of late infection sample reads did map to intergenic regions or within ORFs, perhaps indicative of misannotated ASFV ORFs. Alternatively, these RNAs may be generated by pervasive transcription, a phenomenon observed during late VACV infection ^21, 22^. Another origin of intra-ORF CAGE RNA-5’ signals could be due to mRNA de-capping and degradation followed by re-capping as observed with eukaryotic mRNAs ^23, 24^.

**Fig. 2.**
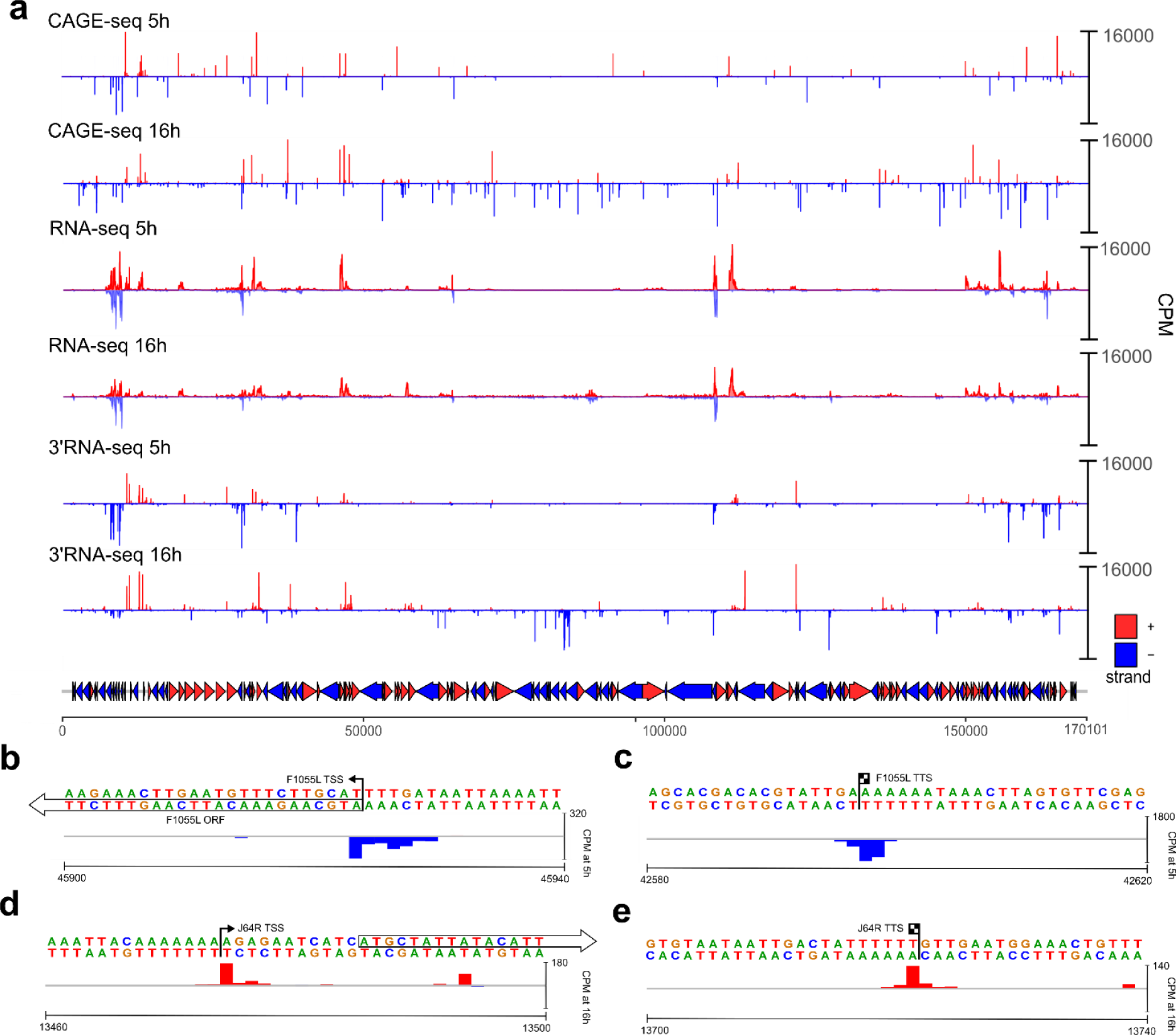
The ASFV transcriptome including transcription start sites and termination sites. (a) Whole genome view of normalized coverage counts per million (CPM) of RNA-seq, 5’ CAGE-seq and 3’ RNA-seq reads. The coverage was capped at 16000 CPM. 153 BA71V annotated ORFs are represented as arrows and coloured according to strand. Peak cluster shape example from F1055L 5’ CAGE-seq ends (b) and 3’ RNA-seq ends (c) showing a wide multi-peaked distribution, and J64R 5’ CAGE-seq (d) and 3’ RNA-seq (e) showing a narrow peak distribution.

### Mapping of ASFV Primary Transcription Start Sites

Following mapping of CAGE-seq reads to the ASFV-BA71V genome, we located regions including an enrichment of reads corresponding to the 5’ ends of transcripts and thereby the transcription start sites (TSS). Using the CAGEfightR ^25^ package we detected a total of 779 TSS organised in clusters with a predominant sharp peak within each cluster (Supplementary Figure 2a), somewhat broader than the 657 termination clusters from 3’ RNA-seq (Supplementary Figure 2b). Six additional TSS clusters were annotated manually since they were clearly detectable when viewing alignments manually but missed by CAGEfightR. Not all ∼780 clusters were located within 500 bp upstream of the ASFV gene translation initiation codons indicative of genuine gene promoters, but within and antisense relative to ORF coding sequences (CDS) (Figure 3a). 28 clusters in total were not found associated with any annotated ASFV-BA71V ORFs. The clusters associated with annotated ORFs and in the sense direction were manually investigated for their feasibility as ‘primary’ TSSs (pTSSs), based on peak height, proximity to the ORF initiation codon, and coverage from our complementing RNA-seq data. We identified pTSSs fulfilling these criteria upstream of 151 of the 153 annotated ORFs in the ASFV-BA71V genome, the missing ORFs being E66L and C62L. Overall, our data showed good agreement with previously individually mapped TSSs of 44 ORFs, because in 86 % the identified pTSSs matched (Supplementary Table 2). Our sequencing data resulted in the re-annotation of eleven ORFs where the pTSS / RNA-seq coverage was not compatible with the ORF annotations in the published ASFV-BA71V genome. Based on the edited annotations (Table 1), we provide a novel gene feature file (Separate ‘GFF’).

**Fig. 3.**
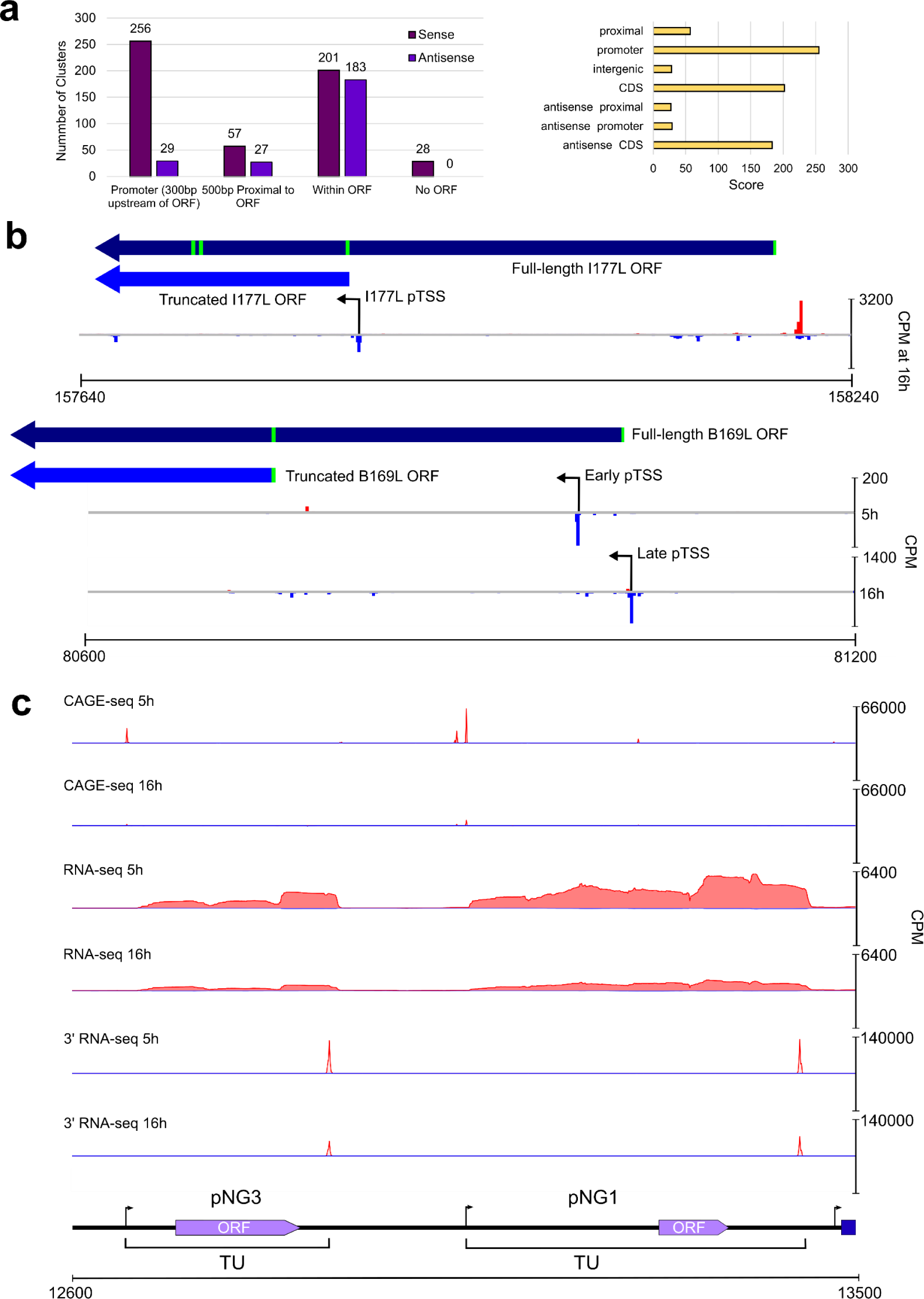
Transcriptome mapping aids the reannotation of the ASFV BA71C genome. (a-left) Summary bar graph of CAGEfightR TSS clusters and their locations relative to the 153 annotated BA71V ORFs. (a-right) Types of CAGEfightR clusters detected and the distribution of their respective CAGEfightR scores. (b) Two examples of ORFs requiring re-annotation following pTSS identification downstream of annotated start codon, encoding shorter ORFs from the pTSS (I177L, above) or during one expression stage (B169L, below). (c) Examples of two putative novel genes (pNG3, left and pNG1 right) annotated with the normalized RNA-seq and CAGE-seq read coverage (CPM) and their genome neighbourhood.

**Table 1.** Summary of ASFV genes where pTSS locations required re-annotation of ORFs. E, I and L refer to alternative pTSSs from early, intermediate and late infection, respectively. TSSs for I243L were compared against co-ordinates reported by Rodríguez *et al*. ^26^.

| ORF | Strand | pTSS<br>Coordinate | Corrected<br>Start Codon<br>Coordinate | ORF<br>Length | Comment |
| --- | --- | --- | --- | --- | --- |
| K93L | - | 2131 | 2122 | 83 | Alternative ATG codon 30 nt downstream. Another strong TSS was detected at 2037- whose transcripts would encode a 36 AA protein. |
| F165R | + | 42354 | 42359 | 136 | Alternative ATG codon 63 nt downstream. |
| C84L | - | 64618 | 64492 \| 64616 | 38 \| 76 | 38 AA ORF was in-frame with original C84L start codon. 76 AA ORF encoded from first ATG after pTSS. |
| G1211R | + | 96370 | 96377 | 1207 | Alternative ATG codon 12 nt downstream. |
| CP204L | - | 108573 | 108567 | 196 | Alternative ATG codon 24 nt downstream. |
| CP312R | + | 110491 | 110501 | 307 | Alternative ATG codon 15 nt downstream. |
| I177L | - | L: 157857 | 157849 | 66 | Strongest pTSS only detected in late time-point. |
| DP93R | + | 167971 | 167980 | 83 | Alternative ATG codon 30 nt downstream. |
| EP402R | + | 56862 | 57104 \| 56991 | 115 \| 148 | Encodes 115 AA in-frame with original EP402R start codon. 148 AA alternative ORF encoded from first ATG after pTSS. |
| **B169L** | - | **E: 80983 \|**<br>**L: 81025** | **81018 \| 80745** | **169 \| 78** | **Late pTSS can produce full-length B169L and early pTSS: 78 AA.** |
| **I243L** | - | **E: 155122**<br>**I: 155124**<br>**L: 155115** | **E/I: 155119**<br>**L: 154969** | **243 \| 191** | **Late pTSS produces shorter transcript with closest downstream ATG encoding a shorter 191 AA protein.** |

Several genes (including B169L and I177L) have a pTSS upstream of the annotated start codon and an alternative TSS apparently residing within the ORF (Figure 3b). This was previously observed for gene I243L that is associated with distinct TSSs for early, intermediate and late stages of gene expression described by Rodríguez *et al*. ^26^, in agreement with our CAGE-seq results (Supplementary Figure 3a). I243L encodes a homologue of the important RNA polymerase II transcript cleavage factor TFIIS that is highly conserved in eukaryotes and among NCLDV members albeit with varying domain conservation^27^. Since the late I243L TSS is downstream of the start codon at the I243L ORF 5’ end, the protein encoded by the shorter mRNA is 52 amino acid (AA) residues smaller. Multiple sequence alignments of ASFV and eukaryotic TFIIS homologues (Supplementary Figure 3b) illustrate ASFV-TFIIS domain organisation. The domains which are important for RNA cleavage (domain III) and RNA pol II-binding (domain II and linker between domains II and III) remain intact, while the N-terminal domain I is largely absent during late infection. The role for this domain in transcription is little understood but appears to be important during preinitiation complex assembly and stability, independent of the transcript cleavage activity ^28, 29^. We identified 7 further genes with alternative pTSSs during early and late infection, these are summarised in Table 2. In most cases, the re-annotated (single pTSS downstream of start codon) or alternative pTSSs (multiple pTSSs, some downstream of start codon) did not substantially alter the protein products of the ORFs, except for re-annotated I177L and alternative pTSSs of B169L (Figure 3 b.), two putative transmembrane proteins ^11, 18^. The longer B169L mRNA synthesised during late infection encodes the complete 169 AA protein, while the early transcript encodes an N-terminally truncated protein of 78 AA, and the predominant late pTSS for I177L would truncate it to less than half its original length.

**Table 2.** Alternative pTSS usage during early and late ASFV infection. List of ASFV genes (besides I243L and B169L) with alternative pTSSs used in early and late infection.

| Gene | Early<br>pTSS | Late<br>pTSS | Function |
| --- | --- | --- | --- |
| X69R | 11315 | 11280 | Uncharacterised |
| J154R | 14174 | 14150 | MGF 300-2R |
| EP1242L | 53125 | 53135 | ASFV-RPB2 |
| C315R | 70137 | 70131 | ASFV-TFIIB |
| CP80R | 110208 | 110191 | ASFV-RPB10 |
| D345L | 129357 | 129257 | Lambda-like exonuclease <sup>6</sup> |
| E120R | 150949 | 150911 | Structural protein <sup>71</sup> |

### Novel Genes Supported by Sequencing Data

28 pTSSs in our CAGE-seq data set were not associated with annotated ORFs (Supplementary Table 3). Our manual analysis of both CAGE-seq and RNA-seq maps revealed that seven of these pTSSs were associated with transcripts that encode short ORFs, which we call putative novel genes (pNGs). These encode polypeptides of 25–56 AA length (Table 3) without a clear similarity to characterised ASFV genes that were likely missed in initial BA71V ORF prediction as only those ≥ 60 AA were annotated ^18^. Figure 3c. illustrates the features of pNG1 and pNG3, with distinct TSS and TTS, and robust RNA-seq read coverage across the entire gene. In support of the notion that the pNGs are *bona fide* ASFV genes, five of seven pNG transcription units include a 5–8 nucleotide poly-T sequence at their 3’ termini; this sequence signature has been proposed to serve as transcription termination motif ^9, 30^.

**Table 3.** Details of the 7 putative novel genes from the 28 non-ORF-associated TSSs. NCBI ORFfinder and BLAST were used to predict the putative encoded ORFs and subsequently analysed for putative homologous sequences ^66, 72^. *: defined as pTSS from CAGE-seq. **: defined from 3’ RNA-seq, underlined transcription ends defined from only RNA-seq. pNG5 is antisense to pNG2, and the pNG6 transcription unit (TU) does not have such a clear ending according to RNA-seq and may overlap with DP42R. pNG7 also is overlapping the pNG4 TU, on the same strand.

| Putative Gene | Strand | Transcription Start* | Transcription End** | Putative protein length (AA) | Similarity according to NCBI Blast | Gene-End oligoT (nt) |
| --- | --- | --- | --- | --- | --- | --- |
| pNG1 | + | 13053 | 13435 | 25 | 13 residues had 92% identity to ASFV-G-ACD-00350 (AZP54308.1), E-value: 0.11 | 6 |
| pNG2 | - | 30091 | 29827 | 50 | 50 residues had 100% identity with ASFV26544 00600 (AKM05534.1) | 8 |
| pNG3 | + | 12664 | 12896 | 44 | 38 residues had 59% identity to ASFV-G-ACD-00290 (AZP54130.1), E-value: 0.13 | 6 |
| pNG4 | + | 10583 | 10835 | 44 | 42 residues had 65% identity with ASFV-G-ACD-00290 (AZP54130.1), E-value: 1e-09. | 6 |
| pNG5 | + | 29817 | <u>30080</u> | 31 | No significant similarity. | None |
| pNG6 | + | 167005 | 167336 | 56 | 56 residues aligned with 40% identity to pKP93L (AIY22188.1), E-value: 6e-07 | 5 |
| pNG7 | + | 10484 | <u>10616</u> | 32 | 32 residues aligned with a 31 AA hypothetical protein with from ASFV Belgium 2018/1 (BioProject: PRJEB31287) 87% identity (VFV47940.1), E-value: 8e-10. | 3 |

### Highly expressed ASFV genes during Early and Late Infection

In order to gain insights into the expression of the individual TUs, we quantified the steady-state mRNA levels using CAGE-seq and compared the most highly abundant mRNAs at early and late time points (Figure 4a). Supplementary table 4 summarises the expression of all detected ASFV-BA71V TUs including the newly annotated pNGs. We also re-defined the ASFV gene transcription unit TUs as pTSS to stop codon (or entire RNA-seq peak for pNGs) and quantified TU expression from RNA-seq which reflected CAGE-seq analysis (Supplementary Figure 4a, Supplementary Table 5). Our highly expressed genes also matched those identified in the viral proteome of infected tissue culture cells determined by mass spectrometry (highlighted in Figure 4a, Supplementary Figure 4a) ^31^. Surprisingly, six genes were found in the top-20 expressed genes during both early and late infection from CAGE-seq (CP312R, A151R, K205R, Y118L, pNG1, I73R). However, a simple comparison of gene steady-state transcript levels would not be sufficient to draw conclusions about differential gene expression between the two time points; a thorough analysis of this kind requires both CAGE- and RNA-seq data sets as described below ^32^.

**Fig. 4.**
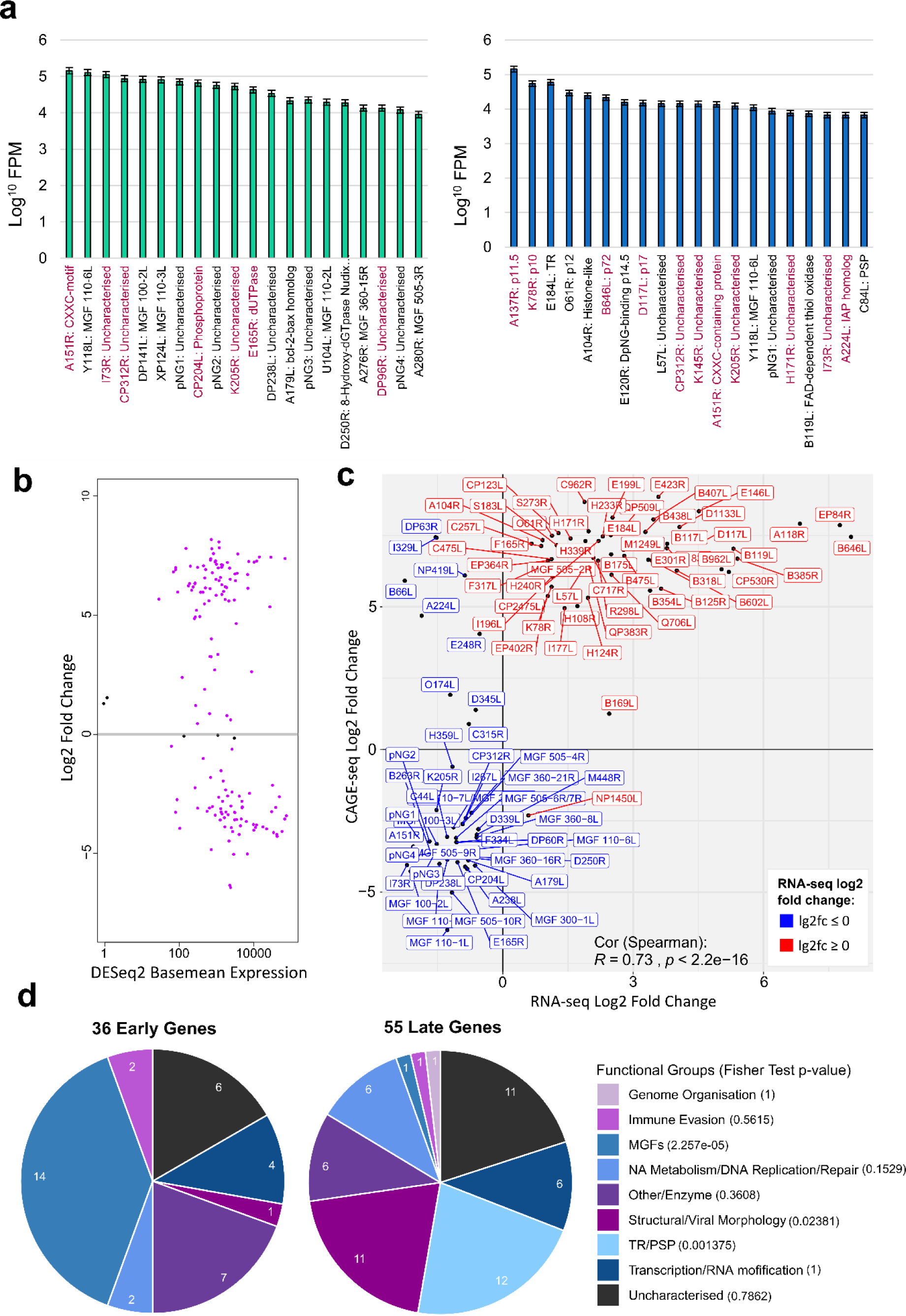
Early and late gene expression of ASFV BA71V. (a) FPKM values for 20 most highly expressed ASFV TUs according to CAGE-seq at 5h (left) and 16h (right) post-infection, RNA-seq equivalent results found in SUPPLEMENTARY Figure 3. Genes highlighted in maroon indicate those encoding proteins which were also found in the 20 most-abundantly expressed AFSV proteins during infection of either WSL-HP, HEK293 or *Vero* cells according to proteome analysis done by Keßler *et al*. ^31^. Gene functions are shown after their name with TR and PSP referring to predicted transmembrane region and putative signal peptide, respectively. (b) MAplot from DESeq2 analysis of CAGE-seq representing the DESeq2 base mean of transcript levels versus their log2 fold change, with significantly differentially expressed genes in purple (adjusted *p*-value < 0.05). (c) Scatter plot comparing log2fold changes of the 101 significantly differentially expressed genes in common between RNA-seq and CAGE-seq. Labels were coloured according to their significant upregulation or downregulation from RNA-seq. (d) Pie chart of gene functional categories downregulated from 5 h to 16 h (36 early genes) and upregulated from 5 h to 16 h (55 late genes). Fisher test carried out on gene counts for functional groups between early and late infection, for this all MGF members were pooled into the ‘MGFs’ functional group.

### Differential Expression of ASFV Genes

We characterised the differential expression of ASFV genes between early and late infection by comparing separate DESeq2 analyses of RNA-seq and CAGE-seq datasets. From RNA-seq data we concluded 103 ASFV TUs showed significant differential expression (adjusted *p*-value < 0.05), with 47 genes down- and 56 genes upregulated during the progression from early (5 hrs) to late (16 hrs) infection (Supplementary Figure 4b). CAGE-seq appeared to be more sensitive, indicating that 149 genes were significantly differentially expressed: 65 genes were down- and 84 genes were upregulated (Figure 4b). 101 genes were congruent in both datasets, and the changes in expression were significantly correlated between CAGE-seq and RNA-seq data sets (Spearman’s rank correlation coefficient ρ = 0.73, Figure 4c). The directions of changes were only reversed for 10 (out of 101) genes between the common RNA-seq and CAGE-seq datasets (DP63R, I329L, NP419L, B66L, A224L, E248R, O174L, D345L, C315R and NP1450L), giving us a total of 91 genes we confidently classified as early (36) and late (55) from our analysis. Supplementary Table 6 provides all details of differentially expressed genes from the RNA-seq and CAGE-seq analyses, their functions, and whether previously detected in viral particles. The 91 genes with correlated differential expression (from both CAGE-seq and RNA-seq) were assigned with functional categories based on their annotation in the VOCS database ^33^ and ASFVdb ^34^ (Figure 4d). Around one fifth of the early and late genes were classified as ‘uncharacterised’ without any functional predictions. The transition between 5 h and 16 h post infection is accompanied by a significant transcript-level increase in genes important for viral morphology and structure, but also the overall diversity of genes differentially expressed changed. A significant difference was seen in the multigene family (MGF) members; they constitute nearly a half of the early genes, but only one (MGF 505-2R) belongs to the late genes. ORFs annotated with a ‘transmembrane region’ or ‘putative signal peptide’ were also overrepresented in the late infection (Fisher Test: *p* < 0.05); they remain poorly characterised beyond a domain prediction and nine proteins (out of 12) of these ORFs could be detected in BA71V virions by mass spectrometry ^11^.

### Architecture of ASFV Gene Promoters and Consensus Elements

The genome-wide TSS map combined with information about their differential temporal expression allowed us to analyse the sequence context of TSSs and thereby characterise the consensus motifs and promoter architecture of early and late ASFV genes. For the sequence analyses we compared our clearly defined 36 early genes and 54 late genes (quartiles in Figure 4c). Eukaryotic RNA pol II core promoters are characterised by a plethora of motifs, including TATA- and BRE boxes, and the Initiator (Inr). The former two interact with the TBP and TFIIB initiation factors, while the latter is interacting with RNA pol II, respectively. Multiple sequence alignments of the regions proximal to the TSS revealed several interesting ASFV promoter signatures that are related to RNA pol II motifs ^35^. The Initiator (Inr) element overlapping the TSS is a feature that distinguishes between early and late gene promoters (Figure 5a and b). The Inr of early genes appears as a TA(+1)NA tetranucleotide motif (where N has no nucleobase preference, Figure 5c), while the late gene Inr shows a strong preference for the sequence TA(+1)TA (Figure 5d), that is not to be confused with the TBP-binding TATA box. Our late Inr consensus motif is in good agreement with a small number (20) of previously characterised late gene promoters ^9, 16^. To search for additional gene promoter elements that likely interact with transcription initiation factors, we extended our search to include sequences up to 40 bp upstream of the TSS. A MEME ^36^ analysis identified and located a significant 19-nt motif (Figure 5a) located approximately 10 bp upstream of the TSS for 36 (out of 36) early gene promoter sequences, which we have called the Early Promoter Motif (EPM). The ASFV EPM is related to the VACV early gene promoter motif (‘Upstream Control Element’ or UCE) ^37, 38^ as well as the yeast Virus-Like Elements (VLEs) promoters ^39^ (Supplementary Figure 5.c). Importantly, the distance distribution between the EPM and the TSS is limited to 9–10 bp (Figure 5b), indicative of constraints defined by distinct protein-DNA interactions, e.g. by transcription initiation factors binding upstream of the TSS and the ASFV-RNAP engaging with the promoter DNA on the TSS. However, the EPM is not limited to the 36 early genes, since a FIMO ^40^ motif search identified the EPM within 60 bp upstream of a much larger subset of 81 TSS/TUs including the pNGs and alternative pTSSs, four of which were the early alternative pTSS, i.e. gene internal promoters, for I243L, B169L, J154L and CP80R. Supplementary Figure 6a illustrates expression profiles of all genes with an EPM upstream according to FIMO, the majority showing a negative log2 fold change between 5 and 16 hours. Using the same approach, we searched for promoter sequence motifs associated with late genes. MEME identified a conserved motif upstream of only 17 (out 54) late genes (E-value: 1.6e-003), which we called Late Promoter Motif (LPM), shown in Figure 5f. The LPM could be identified within 60 bp upstream of 54 (out of 158, including pNGs, Supplementary Figure 5c) ASFV genes, and Supplementary Figure 6b shows the vast majority of genes with an LPM upstream had a positive log2 fold change. However, the spacing between the LPM and the TSS of 4–12 bp shows a much greater diversity compared to the EPM. A Tomtom ^41^ search identified the LPM motif as a match for 28 motifs including the TATA-box motif (*p*-value: 2.85e-03, E-value: 5.16e+00, Supplementary Figure 5d). This was not a strong hit and these motifs only bear a limited resemblance to each other except for their AT-rich bias.

**Fig. 5.**
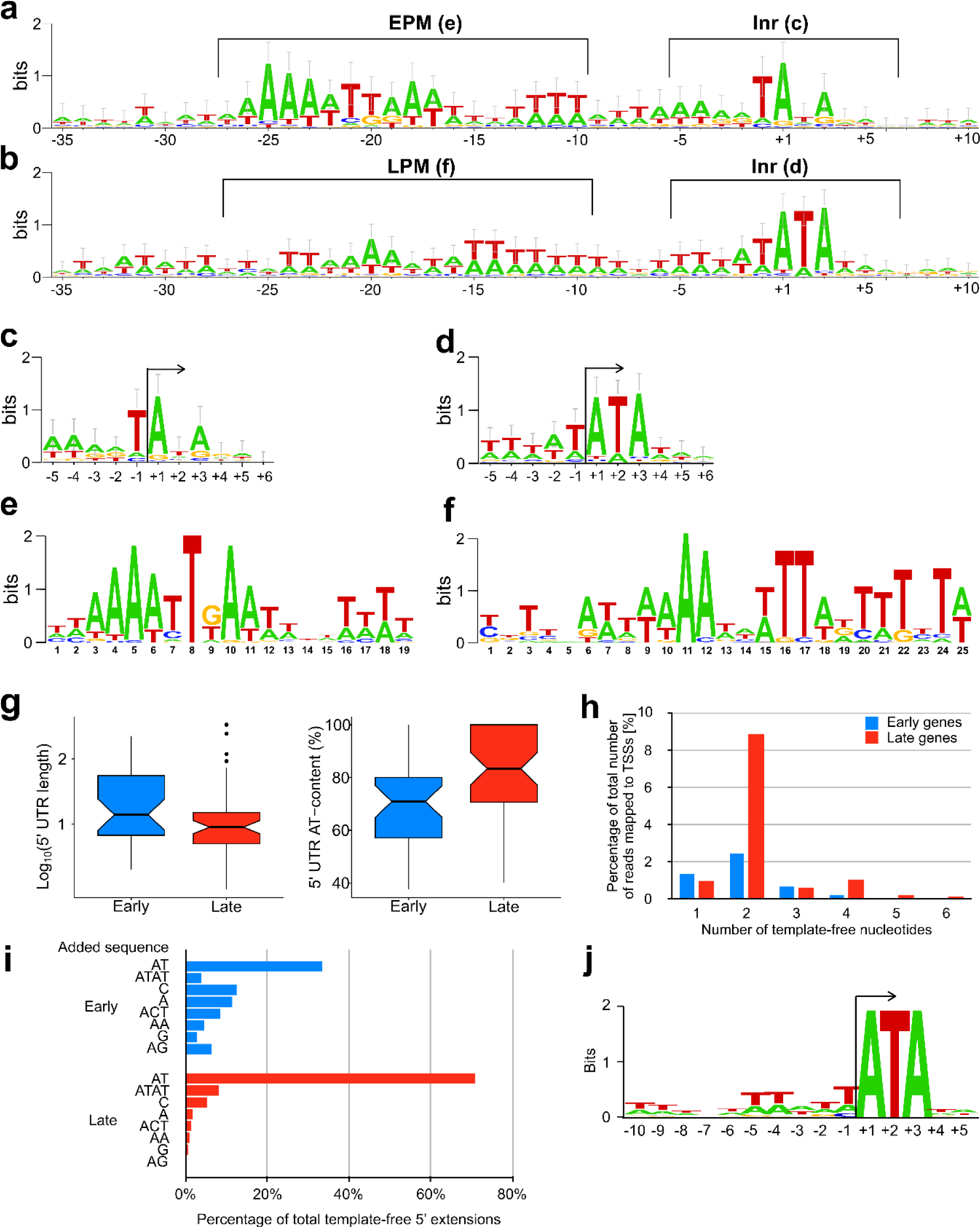
Promoter sequence signatures of early and late ASFV genes. WebLogo 3 ^73, 74^ of aligned early (a) and late (b) sequences surrounding the Inr (+1) from −35 to +10, with annotations of features illustrated in figures c-f. WebLogo 3 consensus motif with error-bars, of the 36 early (c) and 55 late (d) gene sequences surrounding their respective pTSSs (5 nt up- and downstream), i.e. initiator (INR) motif. (e) EPM located upstream of all 36 of our classified early genes according to MEME motif search, FIMO with a threshold of *p*-value < 1.0E-4 then identified at least one iteration of this motif upstream of 81 ASFV genes. (f) The LPM detected upstream of 17 of our classified late genes from MEME motif search. (g) (left) 5’ UTR lengths in nt of the 91 early (mean: 39, median: 14) or late (mean: 25, median: 9) classified ASFV genes, starting from the most upstream pTSS (in the case of alternating pTSSs) until the first ATG start codon nt, represented as a notched boxplot. (g) (right) Percentage AT content of early (mean: 69.0, median: 70.9) and late (mean: 81.7, median: 83.3) 5’ UTRs, omitting those of 0 length. (h) Frequency of different lengths of template-free extensions in early and late stage samples. (i) Frequency of most common template-free extensions in the early and late stage samples. (j) Sequence logo of region surrounding TSS of “AT”-extended transcripts.

### ASFV mRNAs have 5’ leader regions

Early and late genes in ASFV differ with regard to the length of 5’ untranslated ‘leader’ regions (UTR). 5’ UTRs of the late genes are significantly shorter and have a higher AT-content compared to early genes (*p*-value < 0.05, Figure 5c). Surprisingly, a subset of late gene CAGE-seq reads extended upstream of the assigned TSSs and were not complementary to the DNA template strand sequence. In order to rule out any mapping artefacts, we trimmed the CAGE-seq reads by removing the upstream 25 nt, and aligned them to the genome at the 5’ boundary of the reads. This did not significantly impair the mapping statistics but highlighted that nearly half of the annotated TSS (74/158) among both early and late genes are associated with mRNAs that have short 5’ extensions, including 7 genes with multiple TSSs (Supplementary Table 7). The majority of the 5’ leaders are 2 or 4 nt long (Figure 5h and Supplementary Figure 5e) and not correlated with expression phase (Supplementary Figure 5f). The most common sequence motif is AT (33% and 71% of early and late genes, respectively) and ATAT (7% in late genes, Figure 5i). In order to investigate any potential sequence-dependency of the mRNAs associated with AT- and ATAT-5’ leaders, we scrutinised the template DNA sequence downstream of the TSS and found that all TUs, contained the motif ATA at positions +1 to +3 (Figure 5i-j). This suggests that the formation of AT-leaders is generated by RNA polymerase slippage on the first two nucleotides of the initial A(+1)TANNN template sequence, generating AUA(+1)UANNN or AUAUA(+1)UANNN mRNAs. A different but related slippage has been observed in the VACV transcription system, where all post-replicative mRNAs contain short polyA leaders which are associated with consensus Inr TAAAT motif ^21^. The structural determinants underlying RNAP slippage are interactions between the template DNA sequence and the RNAP and/or transcription initiation factors; the differential use of distinct initiation factors for the transcription of early and late ASFV genes thus likely accounts for difference in leader sequences.

### Transcription termination of ASFV-RNAP

Mapping of the 3’ mRNA ends of a several ASFV genes revealed a conserved sequence motif consisting of ≥7 thymidylate residues in the template, which is consistent with native 3’ end formation generated by transcription termination ^30^. In order to investigate the sequence context of transcription termination, mRNA 3’ end sequencing was used to obtain the sequences immediately preceding the poly(A) tails, generating a genome-wide map of mRNA 3’ end peaks (Figure 2). Using a similar approach as for pTSS mapping, CAGEfightR detected a total of 657 termination site clusters, 212 TTSs within 1000 bp downstream of 1–3 ORFs. Because multiple ORFs had more than one cluster within that region (Supplementary Table 8), we defined 114 primary transcription termination site (pTTS) as the the TTS with the highest CAGEfightR-score in closest proximity to a a stop codon; we classified the 98 remaining peaks as non-primary TTSs (npTTS). By carrying out sequence analyses similar to the TSS mapping, we identified a highly conserved poly-T signal within 10 bp upstream of 126 TTSs (83 pTTSs, 43 npTTSs) that was characterised by ≥4 consecutive T residues (Figure 6a), with the ultimate residue located <3 bp of the last T residue in the motif (Figure 6b). The remaining 86 TTSs were not associated with any recognisable sequence motif besides a a single T residue 1 bp upstream of the TTS. Our results are in good agreement with a previous S1 nuclease mapping of 6 ORFs (Supplementary Table 2), but less so with 17 proposed TTSs which were predicted based on transcript length estimates relative to upstream transcription start sites (Supplementary Table 2). This may be because only ≥7 consecutive Ts in the template were included to serve as terminators. Our results demonstrate that the total number of consecutive Ts of the polyT motif can vary, with polyTs of early genes being longer than those of late genes (Figure 6c). Finally, we observed differences between early and late gene termination, in as much as poly-T terminators were overrepresented in early- and underrepresented in late genes (Figure 6d), and the 3’ UTRs of late genes were significantly longer compared to 3’ UTRs of early genes (Figure 6e), in good agreement with previous studies on a small number of mRNAs.

**Fig 6.**
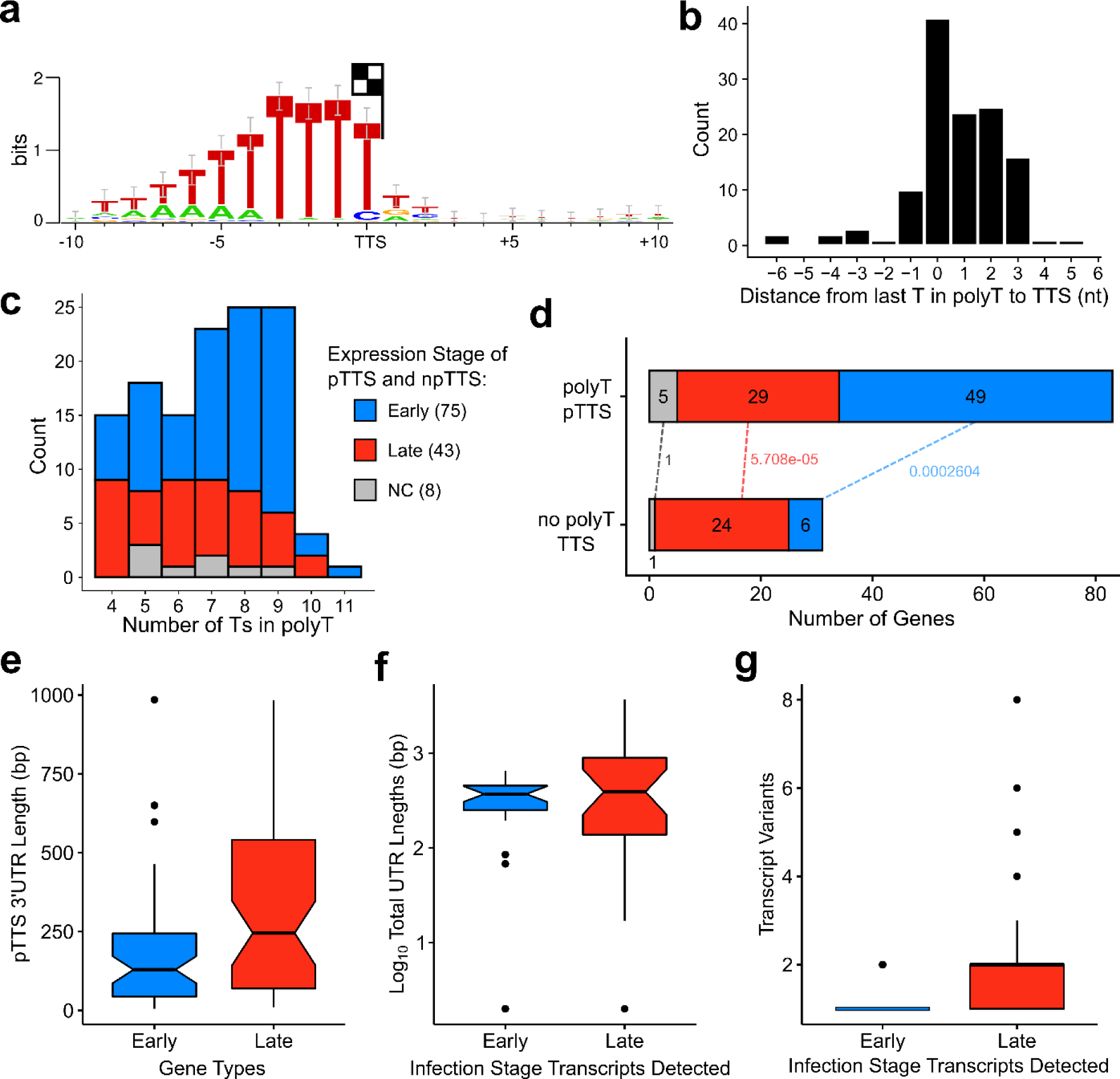
Transcription termination in ASFV. (a) WebLogo 3 motif of 10 nt upstream and 10 downstream of all pTTS and npTTSs with a polyT upstream with ≥4 consecutive Ts (126 TTSs). (b) Distance from 3’ terminal T in polyT to the TTS (median). (c) The distribution of polyT lengths among 126 polyT TTSs (median: 7), split into expression stages according to CAGE-seq differential expression analysis (NC: not-classified), showing late gene polyTs are shorter in length (Wilcoxon rank sum test, *p*-value: 0.0216). (d) Distribution of gene expression types among the 83 polyT pTTSs and 31 non-polyT pTTSs. Dotted lines labels indicate Fisher test *p*-values of gene types between the two pTTS-types, classified from CAGE-seq. (e) 55 Early and 53 late gene 3’ UTR lengths from stop codon to pTTS (Wilcoxon rank sum test, *p*-value: 0.003). (f) Approximate total UTR length (including poly(A) tails) of the shortest full-length transcript variants of 40 ORFs detected during early and/or late infection stage using *Northern blots*. F-test indicated that variance of late UTRs length was greater than that of early UTRs (*p*-value = 3.559e-11). Immediate early and intermediate stage was treated as early and late, respectively, for this and the next figure. For details, see Supplementary Table 2, and references therein. (g) Number of transcription variants of 40 ORFs detected using *Northern blot* during early and/or late infection stage (Wilcoxon rank sum test, *p*-value: 0.001). For details, see Supplementary Table 2, and references therein.

## Discussion

Here we report the first comprehensive ASFV transcriptome study at single-nucleotide resolution. The unequivocal mapping of 158 TSS and 114 TTS for the 160 viral genes allowed us to reannotate the ASFV BA71V genome. Our results provide detailed information about the differential gene expression during early- and late infection, the sequence motifs for early and late gene promoters (EPM and LPM, and Inr elements) and terminators (poly-T motif), and evidence quasi-templated ‘AU’ RNA-5’ tailing by the ASFV-RNAP.

Eukaryotic RNA pol II and the archaeal RNAP critically rely on the two initiation factors TBP and TFIIB for transcription initiation on all mRNA genes. In contrast, the bacterial RNAP obtains specificity for subsets of gene promoters by associating with distinct sigma (σ for selectivity) factors. The ASFV RNAP is related to the eukaryotic RNA pol II, however, transcription initiation of early- and late genes is directed by two distinct sets of general initiation factors and their cognate DNA recognition motifs. Our TSS mapping demonstrates that early and late gene promoters are associated with a combination of conserved and distinct sequence signatures. The first feature of all ASFV promoters is the Initiator (Inr) element, a tetranucleotide motif overlapping the TSS with an A-residue serving as initiating nucleotides similar to most RNAP systems. The Inr sequence is similar for early-(TA[+1]NA) and late (TA[+1]TA) promoters, likely because the Inr makes sequence-specific contacts with amino acid sidechains of the two largest RNAP subunits (RPB1 and 2). The early- and late promoter motifs (EPM and LPM) are located upstream of the TSS, both are AT-rich and different from each other (Figure 5). The distance distribution of the EPM is narrow (located 9–10 bp upstream of the TSS) while the distance between the LPM and TSS shows greater variation and is located closer (4–6 bp) to the TSS. We cannot strictly rule out a limited overlap between early and late genes, or that some genes have hybrid promoters that would enable the expression of genes during both early and late stages, both of which might impede the concise analysis of specific promoter consensus motifs. To unequivocally attribute factors to their cognate binding motifs, a chromatin immunoprecipitation approach is required. Considering the close relationship between ASFV and VACV, we posit that the ASFV EPM is recognised by heterodimeric ASFV-initiation factor (VACV D6/A7)^10^ consistent with the late expression of D1133L/G1340L (Figure 4, also ref ^43^). The presence of D1133L/G1340L along with RNAP in viral particles ^11^ provides a system that is primed to initiate ASFV transcription of early genes. ASFV-TBP (B263R) is an early gene and ASFV-TFIIB (C315R) is expressed throughout infection. We posit that the LPM is utilised by the ASFV TBP and TFIIB homologues, neither of which were detected in virions ^11^. A functional comparison of the ASFV LPM to the classical RNA pol II core promoter elements BRE/TATA-box is compelling. However, the tight spacing between the LPM and the TSS is incompatible with the overall topology of a classical eukaryotic and archaeal TATA-TBP-TFIIB-RNA pol II preinitiation complex (PIC), where the BRE/TATA promoter elements are located ∼ 24 bp upstream of the TSS ^44^. Considering low sequence conservation between cellular and ASFV-TBP ^7^ and unusual spacing of LPM and Inr, the structure of ASFV LPM-TBP-TFIIB-RNAP PIC is likely very different from canonical RNA pol II PICs. Additional factors including ASFV B175L and B385R may contribute to the PIC, as was proposed for VACV-A1 and A2^45, 46^.

We found several examples of alternative, gene-internal, TSS utilisation with the potential to increase the complexity of the viral proteome; protein variants provide the means to generate distinct functionalities. Our TSS mapping uncovered a form of transcript slippage by the ASFV RNAP occurring on promoters that start with an A(+1)TA motif, where mRNAs are extended by one or two copies of the dinucleotide AU. This is reminiscent of VACV, where late gene transcripts containing a poly-A 5’ UTR^21^ are associated with improved translation efficiency and reduced reliance on cap-dependent translation initiation ^47, 48^; likewise, distinct functional attributes of polyA leaders in translation have been documented in eukaryotes ^49^. Whether the 5’ AU- and AUAU-tailing is a peculiarity of the ASFV-RNAP initiation, or whether mRNA 5’ leaders have any functional implications, remains to be investigated.

The mechanisms underlying transcription termination of multisubunit RNAP are diverse ^50, 51^. Our analyses of genome-wide ASFV RNA-3’ ends allowed the mapping of the ASFV ‘terminome’. Over half of ASFV gene mRNA 3’ are characterised by a stretch of seven U residues, with the TTS mostly coinciding with the last T residue in the template DNA motif, which is in good agreement with a few ASFV terminators that have been characterised individually ^30, 52^. In contrast, VACV appears to utilise *a* motif ∼ 40 nt upstream of the mRNA 3’ ^53, 54^. In essence, the ASFV-RNAP is akin to archaeal RNAPs and RNA pol III, where a poly-U stretch is the sole *cis*-acting motif without any RNA secondary structures characteristic of bacterial intrinsic terminators ^51^. The mapped mRNA 3’ ends of genes without any association with poly-U motifs are still likely to represent *bona fide* nascent termination sites, since RNA-seq reads were decreasing towards these termination sites. ASFV encodes several (VACV-related) RNA helicases that have been speculated to facilitate transcription termination and/or mRNA release ^9, 55^. Future functional studies will address the molecular mechanisms of termination including the role of putative termination factors.

Understanding the molecular mechanisms of the ASFV transcription system is not only of academic interest. More than 6 million pigs have died in Asia since 2018 and the Chinese pig population, which comprised half of the world’s population, has decreased by 40% (FAO OIE WAHIS). Unless effective vaccines in conjunction with antiviral treatments against the ASFV are developed, the larger part of the global pig population may succumb to this terrible disease or has to be culled to prevent its propagation (OIE, https://www.oie.int). Structure-based drug design is crucially dependent on our knowledge about the fundamental biology of ASFV transcription machinery and the temporal gene expression pattern, and our results directly contribute to these burning issues for animal husbandry.

## Methods

### RNA Sample Extraction from Vero Cells infected with BA71V

*Vero* cells (Sigma-Aldrich, cat #84113001) were grown in 6-well plates, plates and were infected in 2 replicate wells for 5h or 16h with a multiplicity of infection of 5 of the ASFV BA71V strain, collected in Trizol Lysis Reagent (Thermo Fisher Scientific) separately, after growth medium was removed. Infected cells were collected at 5h post-infection (samples for RNA-seq: S3-5h and S4-5h, CAGE-seq: S1-5h and S2-5h and 3’ RNA-seq: E-5h_1 and E-5h_1), and at 16h post-infection (RNA-seq: S5-16h and S6-16h, CAGE-seq: S3-16h and S4-16h, and 3’ RNA-seq: L-16h_1, L-16h_1). RNA was extracted according to manufacturer’s instructions for Trizol extraction and the subsequent RNA-pellets were resuspended in 50µl RNase-free water and DNase-treated (Turbo DNAfree kit, Invitrogen). RNA quality was assessed *via* Bioanalyzer (Agilent 2100), before ethanol precipitation. For CAGE-seq and 3’ RNA-seq, samples were sent to *CAGE-seq* (Kabushiki Kaisha DNAFORM, Japan) and Cambridge Genomic Services (Department of Pathology, University of Cambridge, Cambridge, UK), respectively.

### RNA-seq, CAGE-seq and 3’ RNA-seq Library Preparations and Sequencing

For RNA-seq, samples were resuspended in 100µl RNase-free water, and polyA-enriched using the BIOO SCIENTIFIC NEXTflex™ Poly(A) Beads kit according to manufacturer’s instructions and quality was assessed *via* Bioanalyzer. NEXTflex™ Rapid Directional qRNA-Seq™ Kit was utilised to produce paired-end indexed cDNA libraries from the polyA-enriched RNA samples, according to the manufacturer’s instructions. Per-sample cDNA library concentrations were calculated *via* Bioanalyzer, and Qubit Fluorometric Quantitation (Thermo Fisher Scientific). Sample S3-5h, S4-5h, S5-16h and S6-16h cDNA libraries were twice separately sequenced on Illumina MiSeq generating 75 bp reads (Supplementary Table 1) and 12 FASTQ files.

Library preparation and CAGE-sequencing of RNA samples S1-5h, S2-5h, S3-16h and S4-16h was carried out by *CAGE-seq* (Kabushiki Kaisha DNAFORM, Japan). Library preparation produce single-end indexed cDNA libraries for sequencing: in brief, this included reverse transcription with random primers, oxidation and biotinylation of 5’ mRNA cap, followed by RNase ONE treatment removing RNA not protected in a cDNA-RNA hybrid. Two rounds of cap-trapping using Streptavidin beads, washing away uncapped RNA-cDNA hybrids. Next, RNase ONE and RNase H treatment degraded any remaining RNA, and cDNA strands were subsequently released from the Streptavidin beads and quality-assessed *via* Bioanalyzer. Single strand index linker and 3’ linker was ligated to released cDNA strands, and primer containing Illumina Sequencer Priming site was used for second strand synthesis. Samples were sequenced using the Illumina HiSeq platform producing 76 bp reads (Supplementary Table 1). 3’ RNA-seq was carried out with samples E-5h_1, E-5h_2, L-16h_1 and L-16h_2 using the Lexogen QuantSeq 3’ mRNA-Seq Library Prep Kit FWD for Illumina according to manufacturer’s instructions. Library preparation and sequencing were carried out Cambridge Genomic Services (Department of Pathology, University of Cambridge, Cambridge, UK) on a single NextSeq flowcell producing 150 bp (Supplementary Table 1).

### Sequencing Quality Checks and Mapping to ASFV and Vero Genomes

FastQC ^56^ analysis was carried out on all FASTQ files: for RNA-seq FASTQ files were uploaded to the web-platform Galaxy (www.usegalaxy.org/) ^57, 58^ and all reads were trimmed by the first 10 and last 1 nt using FASTQ Trimmer ^59^. After read-trimming, FASTQ files originating from the same RNA samples were concatenated. RNA-seq reads were mapped to the ASFV-BA71V (NC_001659.2) and *Vero* (GCF_000409795.2) genomes using Bowtie 2 directly after trimming ^20^, with alignments output in SAM file format. FASTQ analysed CAGE-seq reads showed consistent read quality across the 76 bp reads, except for the nucleotide 1. This was an indicator of the 5’ mRNA methylguanosine due to the reverse transcriptase used in library preparation ^60^, therefore, the reads were mapped in their entirety to the ASFV-BA71V (U18466.2) and *Vero* (GCF_000409795.2) genomes.

FASTQC analysed 3’ RNA-seq reads showed relatively varying and poorer quality after nucleotide 65. Cutadapt ^61^ was utilised to extract only fastq reads with 18 consecutive A’s at the 3’ end followed by the sample i7 Illumina adapter, selecting only for reads containing the 3’ mRNA end and the polyA tail. The 18A-adapter sequences were then trimmed and FASTQC-analysed reads were mapped *via* Bowtie2 to ASFV-BA71V (U18466.2) and *Vero* (GCF_000409795.2) genomes.

### CAGE Analysis, TSS-Mapping

CAGE-seq mapped sample BAM files were converted to BigWig (BW) format with BEDtools ^62^ genomecov, to produce per-strand BW files of 5’ read ends. Stranded BW files were input for TSS-prediction in RStudio ^63^ with Bioconductor ^64^ package CAGEfightR ^25^. Genomic feature locations were imported as a TxDb object from U18466.2 genome gene feature file (GFF3), modified to include C44L^17^. CAGEfightR was used to quantify the CAGE tag transcripts mapping at base pair resolution to the ASFV-BA71V genome - at CAGE TSSs (CTSSs). CTSS values were normalized by tags-per-million for each sample, pooled and only CTSSs supported by presence in ≥2 samples were kept. CTSSs were assigned to clusters, merging CTSSs within 50 bp of one another, filtering out pooled, TPM-normalized CTSS counts below 25, and assigned a ‘thick’ value as the highest CTSS peak within that cluster. CTSS clusters were assigned to annotated U18466.2 ORFs (if clusters were between 300 bp upstream and 200 bp downstream of an ORF). Clusters were classified ‘tssUpstream’ if located within 300 bp upstream of an ORF, ‘proximal’ if located within 500 bp of an ORF, ‘CDS’ if within the ORF, ‘NA’ if no annotated ORF was within these regions (excepting pNG), and antisense if within these regions but antisense relative to the ORF.

Cluster classification was not successful in all cases, therefore, manual adjustment was necessary. Integrative Genomics Viewer (IGV) ^65^ was used to visualise BW files relative to the BA71V ORFs, and incorrectly classified clusters were corrected. Clusters with the ‘tssUpstream’ classification were split into subsets for each ORF. ‘Primary’ cluster subset contained either the highest scoring CAGEfightR cluster or the highest scoring manually-annotated peak, and the highest peak coordinate was defined as the primary TSS (pTSS) for an ORF. Further clusters associated these ORFs were classified as ‘non-primary’, highest peak as a non-primary TSS (npTSS).

If the strongest CTSS location was intra-ORF and corroborated with RNA-seq coverage, then the ORF was re-defined as starting from the next ATG downstream. For the 28 intergenic CTSSs, IGV was used to visualise if CAGE BW peaks were followed by RNA-seq coverage downstream, and whether the transcribed region encode a putative ORF using NCBI Open Reading Frame Finder ^66^.

### TTS-Mapping

TTSs were mapped in a similar manner to TSSs and CAGEfightR was utilised as above to locate clusters of 3’ RNA-seq peaks, though differed in some respects: input BigWig files contained the 3’ read-end coverage extracted from BAM files using BEDtools genomecov. Clusters were detected for the 3’ RNA-seq peaks in the same manner as before, except merging clusters < 25 nt apart, which detected a total of 567 clusters. BEDtools was used to check whether the highest point of each cluster (TTS) was within 500 bp or 1000 bp downstream of annotated ORFs and pNGs. TTSs were then filtered out if 10 nt downstream of the 3’ end had ≥ 50% As, to exclude clusters potentialy originated from miss-priming. TTS clusters for pNG3 and pNG4 were initially filtered out but included in final 212 TTSs due to their strong RNA-seq agreement. In cases of multiple TTS clusters per gene we defined the highest CAGEfightR-scored one within 1000 bp downstream of ORFs as primary (pTTS) unless no clear RNA-seq coverage was shown, or manually annotated from the literature for O61R ^52^.

### DESeq2 Differential Expression Analysis of ASFV Genes

A new GFF was produced for investigating differential expression of ASFV genes across the genome with changes from the original U18466.2.gff: for all 151 ASFV ORFs which had identified pTSSs, we defined their transcription unit (TU) as beginning from the pTSS coordinate to ORF end. Since no pTSS was identified for ORFs E66L and C62L these entries were left as ORFs within the GFF, while the 7 putative pNGs were defined as their pTSS down to the genome coordinate at which the RNA-seq coverage ends. In 8 cases where genes had alternative pTSSs for the different time-points the TUs were defined as the most upstream pTSS down to the ORF end. For analysing differential expression with the CAGE-seq dataset a GFF was created with BEDtools extending from the pTSS coordinate, 25 bp upstream and 75 bp downstream, however, in cases of alternating pTSSs this TU was defined as 25 bp upstream of the most upstream pTSS and 75 bp downstream of the most downstream pTSS. HTSeq-count ^67^ was used to count reads mapping to genomic regions described above for both the RNA-and CAGE-seq sample datasets. The raw read counts were then used to analyse differential expression across these regions between the time-points using DESeq2 (default normalisation described by Love et al. ^68^) and those regions showing changes with an adjusted *p*-value (padj) of <0.05 were considered significant. Further analysis of ASFV genes used their characterised or predicted functions as found in the VOCS tool database (https://4virology.net/) ^33, 69^ or ASFVdb ^34^ entries for the ASFV-BA71V genome.

### Early and Late Promoter Analysis

DESeq2 results were used to categorise ASFV genes into two simple sub-classes: early; genes downregulated from early to late infection and late; those upregulated from early to late infection. For those with newly annotated pTSSs (151 including 7 pNGs but excluding 15 alternative pTSSs), sequences 30 bp upstream and 5 bp downstream were extracted from the ASFV-BA71V genome in FASTA format using BEDtools. The 36 Early, 55 Late and all 166 pTSSs (including alternative ones) at once were analysed using MEME software (http://meme-suite.org) ^70^, searching for 5 motifs with a width of 10–25 nt, other settings at default. Significant motifs (E-value < 0.05) detected *via* MEME were submitted to a following FIMO ^40^ search (*p*-value cut-off < 0.0001) of 60 nt upstream of the total 166 pTSS sequences (including pNGs and alternative pTSSs), and Tomtom ^41^ search (UP00029_1, Database: uniprobe_mouse) to find similar known motifs.

## Supporting information

Summary Figures and Tables

