## Supplementary material for "Temporal Transcriptome and Promoter Architecture of the African Swine Fever Virus": Summary Figures and Tables

### Supplementary Figures and Tables

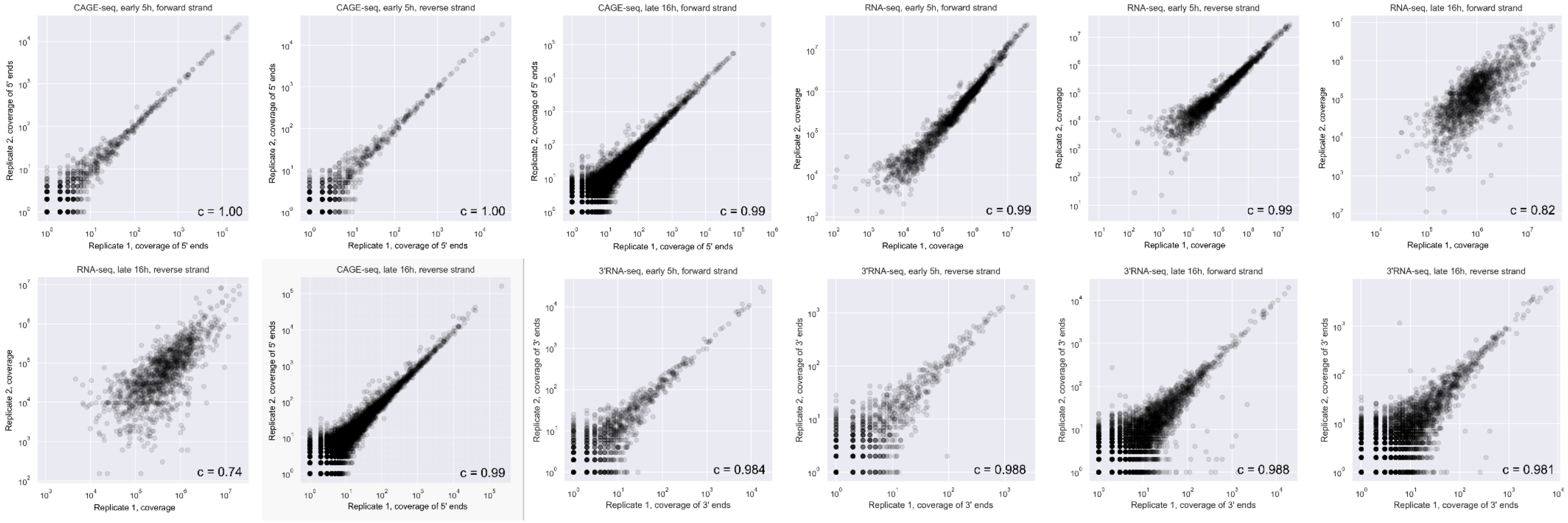


**Supplementary Fig. 1.** Scatter graph comparisons of average mapping coverage (reads per kilobase million, RPKM) in 100 nt bins across the BA71V genome (RPKM) between strands and replicates from RNA-seq, CAGE-seq and 3' RNA-seq, c represents Pearson's Correlation coefficient. Early Replicate 1: S3-5h(RNA-seq), S1-5h (CAGE-seq) and E-5h_1 (3' RNA-seq). Early Replicate 2: S4-5h(RNA-seq), S2-5h (CAGE-seq) and E-5h_2 (3' RNA-seq). Late Replicate 1: S5-16h(RNA-seq), S3-16h (CAGE-seq), and L-16h_1 (3' RNA-seq). Late Replicate 2: S6-16h(RNA-seq), S4-16h (CAGE-seq), and L-16h_2 (3' RNA-seq).

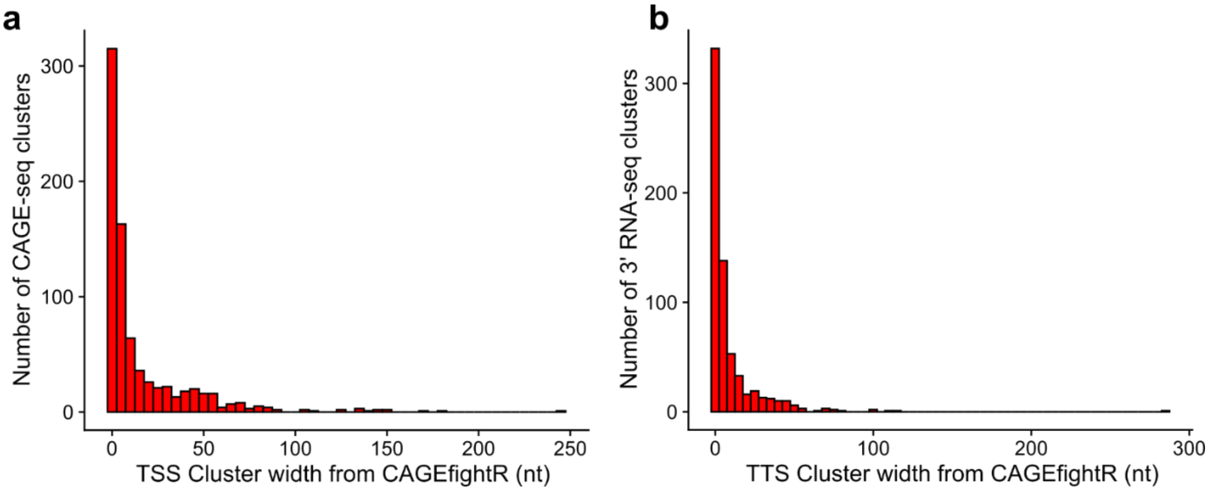


**Supplementary Fig. 2.** Histogram representing the distribution of the cluster widths detected with CAGEfightR analysis from CAGE-seq (a) and 3' RNA-seq (b).

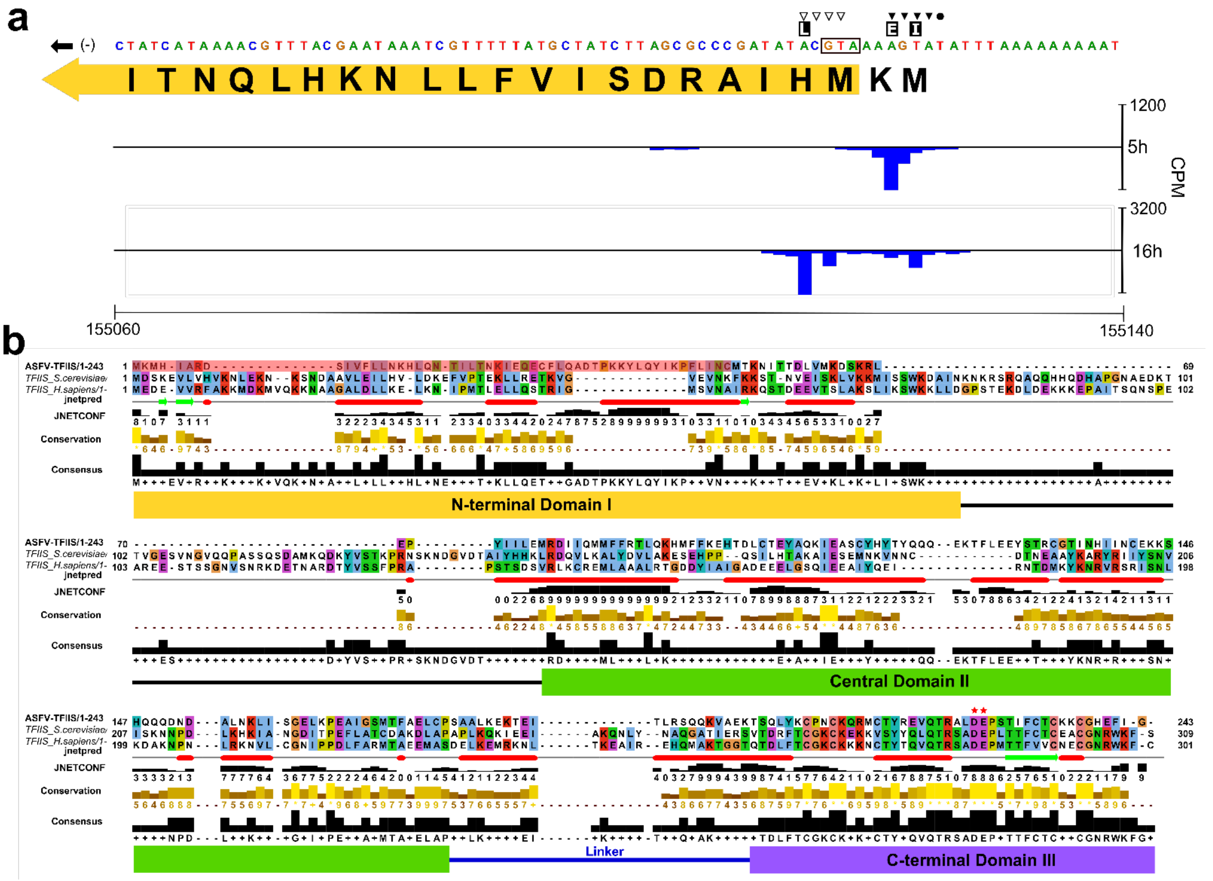


**Supplementary Fig. 3. Analysis of alternative pTSS usage in I243L.** (a) Close up of CAGE-seq alignments 5' end peaks on the minus strand at the start of the I243L ORF. Symbols indicate the TSS sites for early (▼), intermediate (●)and late (▽) gene expression according to Rodríguez *et al.* [1], while E, I and L indicate their respective pTSS positions concluded from our data. The first 21 AA residues of the annotated I243L ORF are shown, in yellow is the re-annotated ORF which could be encoded in transcripts initiating from both our annotated Early pTSS. (b) ClustalW multiple sequence alignment with ClustalX [2,3] colouring, illustrated by Jalview [4], of TFIIS homologues from ASFV (I243L, UniProt: P27948), yeast (UniProt: P07273) and human (P23193). *S. cerevisiae* TFIIS domain locations according to Kettenberger *et al.* [5] are shown below the alignment and acidic (DE) catalytic residues are indicated with★. The ASFV-TFIIS residues highlighted in red are those not encoded in transcripts initiating from the Late pTSS.

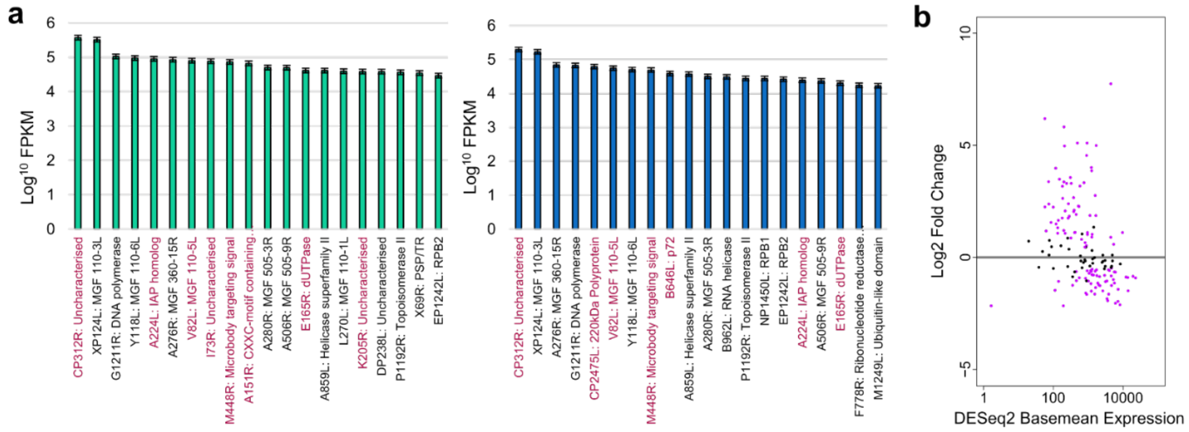


**Supplementary Fig. 4.** (a) 20 most-expressed genes during early (green) and late (blue) infection according to RNA-seq data over gene TU, defined from TSS to ORF stop codon. (b) MAplot representing expression of ASFV TUs including pNGs from DESeq2 analysis of RNA-seq data.

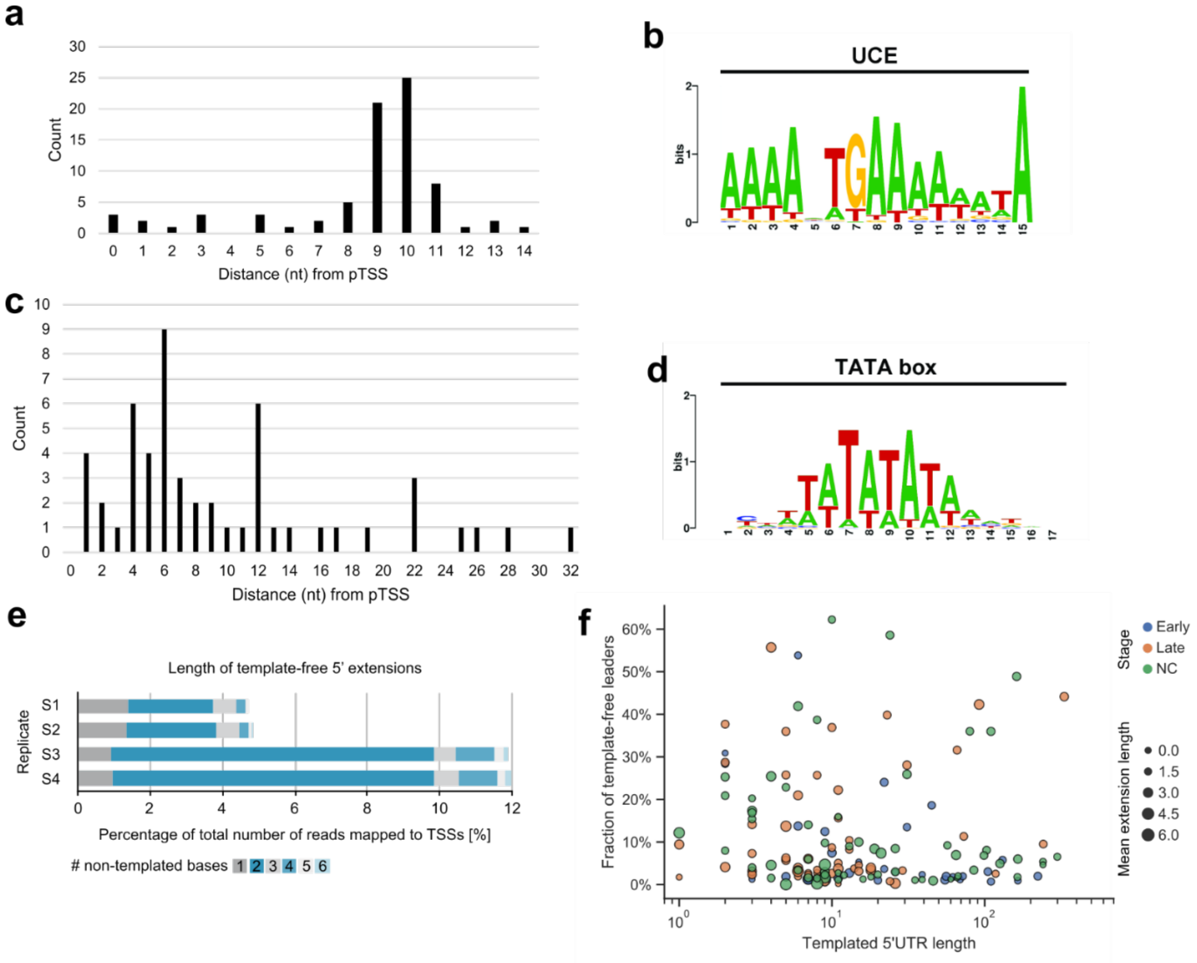


**Supplementary Fig. 5.** (a) Distances of the EPM motif end (nt 19) relative to the 78 pTSSs (alternative pTSSs excluded). (b) VACV Upstream Control Element (UCE) motif located upstream of early genes [6]. (c) Distances from a FIMO search (threshold *p*-value < 1.0E-4) identified the LPM upstream of 53 ASFV genes (excluding those with alternative pTSSs), motif distances from pTSSs are represented as a bar chart. (d) The eukaryotic TATA-box motif which was one of 28 hits in a Tomtom search of the LPM. (e) Frequency of template-free extensions of different lengths in four samples (S1 and S2, early stage after 5 h; S3 and S4, late stage after 16 h). Total number of reads mapped to TSSs corresponds to all CAGE-seq reads that include manually assigned TSSs. (f) Relationship between the length of templated 5' UTRs and fraction of template-free extensions. Gene 5' UTRs split into 36 early (blue), 55 late (orange) and not-classified ('NC', green).

**a**

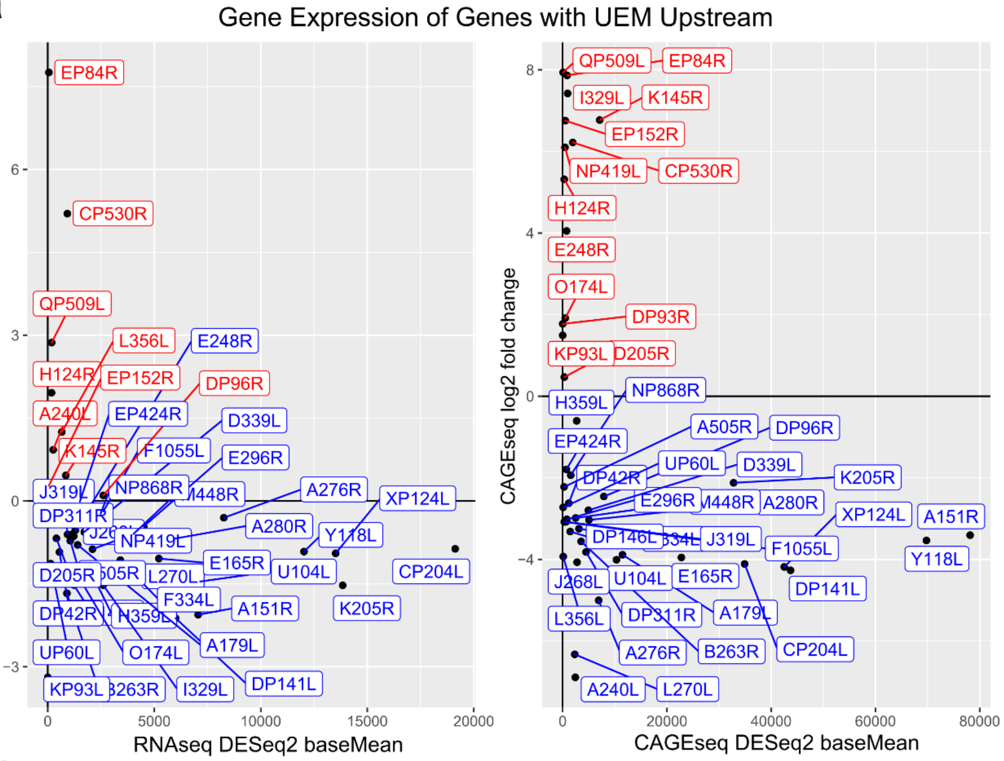


**b**

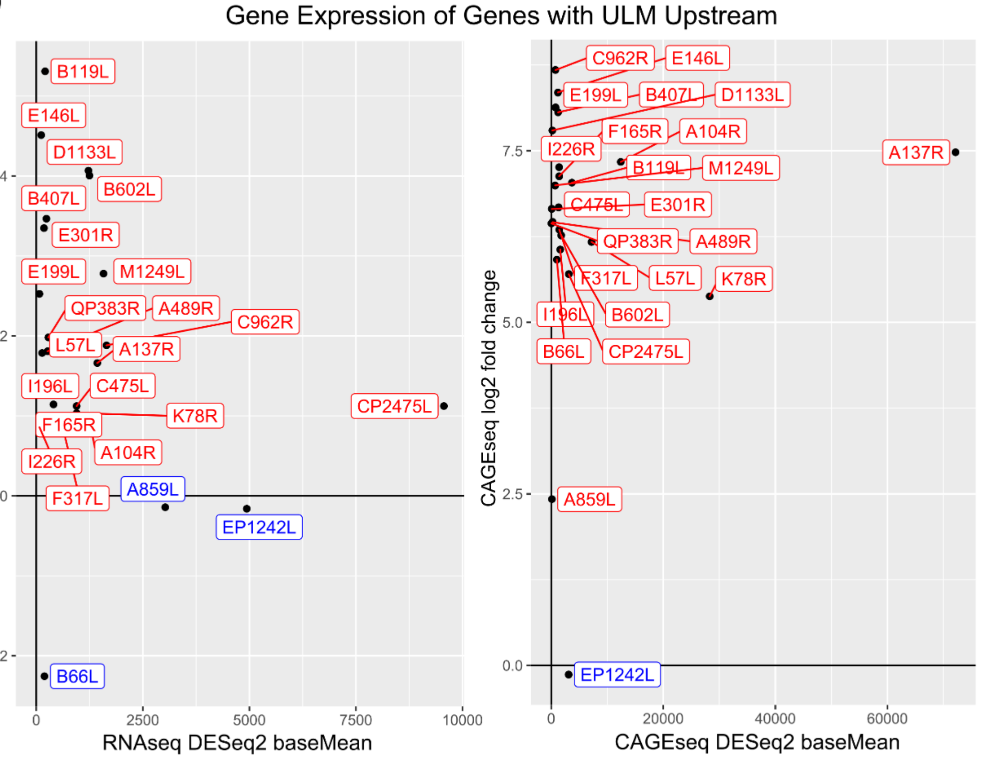


**Supplementary Fig. 6.** Expression profiles from DESeq2 analysis (log2fold change vs. base mean expression) of genes with only an EPM (a) or a LPMs (b) from the FIMO search of 60 bp upstream of pTSSs. Genes for which FIMO detected both EPM and LPM upstream of pTSSs were excluded.

Genes in blue showed a negative log2 fold change (early genes) and in red a positive log2fold change (regardless of significance).

| | **Total Reads in Sample** | **ASFV Genome Alignment Rate** | **Vero Genome Alignment Rate** | **Total Alignment** | **Unmapped** |
|---|---|---|---|---|---|
| **RNA-seq: S3-5h** | 5742140 | 2.8% | 87.8% | 90.6% | 9.4% |
| **RNA-seq: S4-5h** | 10376075 | 2.4% | 78.7% | 81.1% | 18.9% |
| **RNA-seq: S5-16h** | 4530889 | 5.4% | 64.4% | 69.8% | 30.2% |
| **RNA-seq: S6-16h** | 2862427 | 5.5% | 80.2% | 85.7% | 14.3% |
| **CAGE-seq: S1-5h** | 20784893 | 1.8% | 78.1% | 79.9% | 20.1% |
| **CAGE-seq: S2-5h** | 21527000 | 1.7% | 78.5% | 80.2% | 19.8% |
| **CAGE-seq: S3-16h** | 25622459 | 12.5% | 67.1% | 79.6% | 20.4% |
| **CAGE-seq: S4-16h** | 23215375 | 12.2% | 69.7% | 81.9% | 18.1% |
| **3’ RNA-seq: E-5h_1 *** | 5844816 | 2.9% | 95.0% | 97.9% | 2.1% |
| **3’ RNA-seq: E-5h_2 *** | 7865113 | 3.0% | 95.0% | 98.0% | 2.0% |
| **3’ RNA-seq: L-16h_1 *** | 5895169 | 6.6% | 90.7% | 97.3% | 2.7% |
| **3’ RNA-seq: L-16h_2 *** | 6325881 | 6.6% | 90.0% | 96.6% | 3.4% |

**Supplementary Table 1. Summary of Bowtie 2 mapping statistics of samples from RNA-seq.** CAGE-seq, and 3’ RNA-seq to *Vero* host and ASFV-BA71V genomes. * 3’ RNA-seq mapping of reads post-filtering described in Methods.

| ORF | Size | Class [a] | Initiation site [b] | Transcript length [c] | Termination site [d] | Proposed TER [e] | Source [f] | INR cluster [g] | TER cluster [h] | INR agreement [i] | TER agreement [j] |
|---|---|---|---|---|---|---|---|---|---|---|---|
| A137R | 414 | late | 37441+, 37442+ | not mapped | not mapped | - | 10933729 | 37441+ (127) | 38077+ (92) | YES | N/A |
| A151R | 456 | early | 31498+, 31499+ | not available / not mapped | not mapped | - | 23041356 | 31498+ (111) | 32167+ (68) | YES | N/A |
| A224L | 675 | late | 29816-, 29819- | 0.75kb, 1.3 kb | not mapped | 28707-, 29075- | 8553574 | 29816- (498) | 29074- (385) | YES | YES, also YES for cluster 384 |
| A224L | 675 | late | not mapped | 0.7 kb, 1.0 kb, 1.5 kb | not mapped | 28271-, 28707-, 29075- | 19477086 | 29816- (498) | 29074- (385) | N/A | YES, also YES for cluster 384 and 383 |
| A240L | 723 | early | 31206-, **31207-** | ~ 1.0 kb | not mapped | 30175- | 8393914 | 31207- (503) | 29827- (391) | YES | NO, YES for cluster 395 |
| A280R | 843 | early+late | 19352+, 19353+ | 1.1 kb | not mapped | 20389+ | 8139051 | 19353+ (68) | 20387+ (45) | YES | YES |
| A489R | 1470 | late | **17714+, 17715+** | **2.8 kb,** 4.5 kb | not mapped | **20389+** | 8139051 | 17716+ (66) | - | YES | N/A, but YES for cluster 45 |
| A498R | 1497 | early | 21939+, 21940+ | 1.9 kb | not mapped | 23622+ | 8139051 | 21941+ (79) | 23619+ (50) | YES | YES |
| A505R | 1518 | early | 20394+, 20395+, 20396+ | 1.9 kb | not mapped | 21929+ | 8139051 | 20397+ (74) | 21929+ (48) | YES | YES |
| A506R | 1521 | early+late | 25517+, 25518+, 25522+, 25524+, 25525+, 25526+ | 2.0 kb | not mapped | 27360+ | 8139051 | 25526+ (93) | 27357+ (52) | YES | YES |
| A528R | 1587 | early | 23590+ | 2.2 kb | not mapped | 25516+ | 8139051 | 23632+ (87) | 27357+ (53) | NO | NO |
| A542R | 1629 | early | 27369+, 27370+ | 2.1 kb | not mapped | 29117+ | 8139051 | 27370+ (98) | - | YES | N/A |
| B318L | 957 | late | 79335-, **79333-, 79332-**, **79331-** | 1.1 kb, 1.7 kb, 2.4 kb, 3.0 kb, **4.4 kb**, 6.6 kb, **8.0 kb**, 9.4 kb | not mapped | **78219-** | 9083080 | 79331- (597) | 78229- (485) | YES | NO |
| B438L | 1317 | late | 80537-, **80504-, 80502-** | 0.5 kb, 1.0 kb, 2.3 kb, 2.5 kb, 2.8 kb, **4.2 kb**, 5.5 kb | not mapped | **78219-** | 10640542 | 80503- (600) | 79127- (489) | YES | NO |
| B646L | 1941 | late | not mapped | **2.0 kb**, 5.0 kb | not mapped | **86786- ***, 83398- * | 17035321 | 88735- (627) | 86645- (520) | N/A | NO |
| B646L | 1941 | late | 88735-, 88736-, 88737-, 88738- | not mapped | not mapped | - | 10933729 | 88735- (627) | 86645- (520) | YES | N/A |
| C129R | 390 | late | 63933+, 63935+, 63937+ | not available / not mapped | not mapped | - | 23041356 | 63934+ (192) | 64452+ (167) | YES | N/A |
| CP204L | 615 | early | 108575-, 108576-, 108577- | not available / not mapped | not mapped | - | 23041356 | 108573- (666) | 107914- (551) | YES | N/A |

| ORF | Size | Class [a] | Initiation site [b] | Transcript length [c] | Termination site [d] | Proposed TER [e] | Source [f] | INR cluster [g] | TER cluster [h] | INR agreement [i] | TER agreement [j] |
|---|---|---|---|---|---|---|---|---|---|---|---|
| CP2475L | 7428 | late | 107908-, 107909- | not available / not mapped | not mapped | - | 23041356 | 107907- (663) | - | YES | N/A |
| CP530R | 1593 | late | 108610+, 108611+ | not available / not mapped | not mapped | - | 23041356 | 108611+ (251) | - | YES | N/A |
| D117L | 354 | late | 127538-, **127540-**, 127542- | 0.5 kb | not mapped | 126941- | 7856088 | 127539- (694) | - | YES | N/A |
| DP238L | 717 | early | 159623-, 159624-, 159626- | not available / not mapped | not mapped | - | 23041356 | 159631- (753) | 158788- (634) | NO | N/A |
| DP311R | 936 | early | not mapped | 1.3 kb | not mapped | - | 2325203 | 159744+ (363) | - | N/A | N/A |
| DP363L | 1092 | early+late | not mapped | 1.5 kb (E+L), 1.7 kb (L), 2.0 kb (L), 0.6 kb (L) | not mapped | - | 2325203 | 165705+ (384) | 166842+ (322) | N/A | N/A |
| E165R | 498 | early+late | **149603+**, 149604+, **149605+** | 0.5 kb (E+L) | not mapped | 150119+ | 10515998 | 149604+ (328) | 150117+ (269) | YES | YES |
| E183L | 552 | late | **145978-, 145979-** | 1.45 kb | not mapped | - | 7933107 | 145979- (721) | 145178- (609) | YES | N/A |
| EP153R | 462 | early+late | 55542+ (E), 55642+ (L) | **0.6 kb** (L), **0.7 kb** (E), **1.0 kb** (L), **2.2 kb** (L), **2.3 kb** (E), **2.6 kb** (L), 2.7 kb (L) | not mapped | **56256+**, 57850+, 58360+ | 10639320 | 55541+ (173) | 56502+ (144) | YES, also YES for cluster 174 (late) | NO for pTTN, YES for cluster 153 |
| EP402R | 1209 | late | **56220+, 56221+**, 56237+ | **1.6 kb,** 2.2 kb | not mapped | **57850+**, 58359+ | 8102411 | 56221+ (177) | 57850+ (153) | YES | YES |
| G1211R | 3636 | early+late | **96369+** | **4.1 kb** (E+L) | not mapped | - | 8293992 | 96370+ (240) | 100168+ (199) | YES | N/A |
| H339R | 1020 | late | not mapped | 3.0 kb, 5.0 kb, 6.0 kb, 6.5 kb, 7.0 kb, 10. kb | not mapped | **139757-** | 12208975 | 136394+ (299) | 138527+ (248) | N/A | NO |
| I177L | 534 | late | 158196-, 158197-, 158198-, 158199-, 158200- | 1.6 kb, 2.3 kb | not mapped | - | 23041356 + 1309282 | 158127- (747) | - | NO | N/A |
| I196L | 591 | late | 158772-, 158773- | 2.0 kb | not mapped | - | 23041356 + 1309282 | 158773- (749) | 158130- (I196L-man) | YES | N/A |
| I215L | 648 | early + late | 157668-, 157669- | 1.1 kb (E+L), 1.7 kb (L), 2.3 kb (L), 3.3 kb (E+L) | not mapped | - | 23041356 + 1309282 | 157668- (745) | 156714- (627) | YES | N/A |
| I226R | 681 | interm+late | 153706+..153709+ (L), 153716+..1537017+ (I) | 1.0 kb | not mapped | 154698- | 8970983 + 1309282 | 153709- (342) | 155328- (289) | YES | NO |
| I243L | 732 | E+I+L | 155115-..18- (L), 155122-..25- (E), 155126- (I) | 0.8 kb | not mapped | 154234- | 8970983 + 1309282 | 155124- (732) | 154331- (618) | YES | NO |
| I267L | 804 | early | 153677-, 153678- | not available / not mapped | not mapped | - | 23041356 | 153675- (730) | - | YES | N/A |

| ORF | Size | Class [a] | Initiation site [b] | Transcript length [c] | Termination site [d] | Proposed TER [e] | Source [f] | INR cluster [g] | TER cluster [h] | INR agreement [i] | TER agreement [j] |
|---|---|---|---|---|---|---|---|---|---|---|---|
| I329L | 990 | late | 156665-, 156666-, 156667- | 1.8 kb | not mapped | - | 23041356 + 1309282 | 156665- (739) | 155276- (623) | YES | N/A |
| I73R | 222 | early | 155164+, 155165+ | 0.6 kb, 1.6 kb | not mapped | - | 23041356 + 1309282 | 155165+ (347) | 155700+ (293) | YES | N/A |
| J154L | 465 | late | 14147+, 14249+ | 0.6 kb | not mapped | - | 9049318 | 13863+ (50) | 14821+ (38) | NO for pTSS, YES for cluster 52 | N/A |
| J268L | 807 | early+late | **12560-** (E), **12561-** (E), 12870- (L) | **1.1 kb** (E), **1.4 kb** (L), 1.5 kb (E), 1.9 kb (L) | not mapped | **11566-** | 9049318 | 12560- (447) | - | YES, also YES for cluster 449 (late) | N/A |
| J319L | 960 | early | not mapped | 1.3 kb | not mapped | - | 2325203 | 16899- (463) | 15916- (368) | N/A | N/A |
| K78R | 237 | late | 46789+, 46791+, 46793+ | not available / not mapped | not mapped | - | 23041356 | 46793+ (146) | 47428+ (116) | YES | N/A |
| KP360L | 1083 | early+late | not mapped | 1.5 kb (E+L), 1.1 kb (L) | not mapped | - | 2325203 | 3305- (405) | 2174- (332) | N/A | N/A |
| KP362L | 1089 | early+late | not mapped | 1.4 kb | not mapped | - | 2325203 | 4524- (412) | - | N/A | N/A |
| L270L | 813 | imm. early | 8160- | 1.3 kb | **7268- to 7279-**, 7147- to 7157-, 7064- to 7074- | N/A | 2325202 + 1404609 | 8161- (429) | 7150- (342) | YES | YES |
| L83L | 252 | late | 5589-, 5590- | not available / not mapped | not mapped | - | 23041356 | 5528- (416) | - | NO | N/A |
| O61L | 186 | late | 112007+, **112008+**, 112009+ | 1.25 kb | 113180+ to 113186+ | N/A | 8416381 | 112008+ (262) | 113189+ (O61R-man) | YES | YES |
| S273R | 822 | late | **129851+**, 129852+ | **4.5 kb**, longer minor variants | not mapped | **134359+** | 11031264 | 129851+ (287) | - | YES | N/A |
| U104L | 315 | early+late | 8543-, 8544-, **8545-**, **8546-**, 8547-, 8548- | **0.4 kb** (E+L), 0.7 kb (L) | 8188- to 8195- | N/A | 2325202 + 1404609 | 8545- (431) | 8190- (345) | YES | YES |
| V82L | 249 | early | **9499-**, 9500-, **9501-** | 0.9 kb | 9173- to 9178- | N/A | 2325202 + 1404609 | 9496- (434) | 9172- (351) | NO | YES |
| XP124L | 375 | early+late | **9072-**, 9073-, 9074- | **0.6 kb**, 0.8 kb | 8599- to 8604-, 8574- to 8579- | N/A | 2325202 + 1404609 | 9072- (433) | 8601- (348) | YES | YES |
| Y118L | 357 | early | 10064-, **10066-**, **10067-** | 0.55 kb | 9598- to 9607- | N/A | 2325202 + 1404609 | 10066- (437) | 9600- (354) | YES | YES |

**Supplementary Table 2. Summary of ASFV transcript TSSs, TTSs and lengths available in the literature.**
a. Classes are assigned based on relative abundance of transcripts determined *via* Northern blot or primer extension during different time points. Classes are: immediate early (IE) early (E), intermediate (I) and late (L).

b. Initiation sites determined *via* primer extension or S1 nuclease mapping. Major initiation sites are in bold. Initiation sites specific to certain time points are indicated by the time points in brackets.

c. Transcript length determined *via* Northern blot. Major transcription variants are in bold. Transcription variants specific to certain time points are indicated by the time points in brackets.

d. Termination sites determined *via* S1 nuclease mapping. Major termination sites are in bold.

e. Proposed termination sites based on length of transcripts and occurrence of 7T motifs. Putative primary terminations sites of the major transcription variants are in bold. * These proposed 7T motifs are on the opposite strand, therefore, they are not the conventional termination sites.

f. PMID numbers used as standard article identifier.

g. Initiation sites determined *via* cluster identification in CAGE-seq data. Unique cluster number is indicated in brackets.

h. Termination sites determined *via* cluster identification in 3' mRNA-seq data. Unique cluster number or name (in case of manually annotated clusters) is indicated in brackets.

i. Agreement of previously determined initiation sites with initiation site clusters identified in CAGE-seq data within 2 nt tolerance.

j. Agreement of previously determined or proposed termination sites with termination site clusters identified in 3' mRNA-seq data within 3 nt tolerance.

| Start Coordinate | Stop Coordinate | Cluster Width | Strand | CAGEfightR Score | TSS Location | RNA-seq Coverage | ORF | Name | Comments |
|---|---|---|---|---|---|---|---|---|---|
| 418 | 418 | 1 | + | 26 | 418 | Yes | - | - | - |
| 5829 | 5831 | 3 | + | 6477 | 5829 | Yes | - | - | Antisense L356L |
| *10477* | *10485* | *9* | + | *5369* | *10484* | *Yes* | *Yes-short* | *pNG7* | *Overlaps pNG4* |
| *10581* | *10585* | *5* | + | *20016* | *10583* | *Yes* | *Yes* | *pNG4* | * |
| 10695 | 10695 | 1 | + | 63 | 10695 | Yes | - | - | - |
| 10799 | 10816 | 18 | - | 685 | 10808 | Yes | - | - | - |
| 11035 | 11035 | 1 | - | 92 | 11035 | No | - | - | - |
| *12659* | *12667* | *9* | + | *37667* | *12664* | *Yes* | *Yes* | *pNG3* | * |
| 12767 | 12770 | 4 | + | 256 | 12767 | Yes | - | - | - |
| 12873 | 12911 | 39 | + | 3332 | 12910 | No | - | - | - |
| *12995* | *13055* | *61* | + | *128698* | *13053* | *Yes* | *Yes-short* | *pNG1* | * |
| 13106 | 13106 | 1 | - | 205 | 13106 | No | - | - | - |
| 13248 | 13252 | 5 | + | 9807 | 13250 | Yes | - | - | - |
| 13776 | 13776 | 1 | + | 137 | 13776 | Yes | - | - | - |
| 14027 | 14074 | 48 | - | 193 | 14074 | Yes | - | - | Antisense to J64R |
| *29815* | *29822* | *8* | + | *7110* | *29817* | *Yes* | *Yes* | *pNG5* | * |
| *30089* | *30132* | *44* | - | *98243* | *30091* | *Yes* | *Yes* | *pNG2* | * |
| 32133 | 32136 | 4 | - | 157 | 32135 | No | - | - | - |
| 55089 | 55112 | 24 | - | 344 | 55100 | No | - | - | - |
| 64946 | 64959 | 14 | - | 15800 | 64956 | Yes | - | - | Antisense to C44L |
| 156727 | 156729 | 3 | + | 429 | 156727 | Yes | - | - | Antisense I215L |
| 165222 | 165242 | 21 | + | 439 | 165227 | Yes | - | - | - |
| 165313 | 165370 | 58 | - | 1364 | 165334 | No | - | - | - |
| 165468 | 165468 | 1 | - | 77 | 165468 | Yes | - | - | - |
| 165579 | 165581 | 3 | - | 2477 | 165580 | Yes | - | - | - |
| 165650 | 165667 | 18 | - | 173 | 165650 | Yes | - | - | - |
| *167002* | *167006* | *5* | + | *5780* | *167005* | *Yes* | *Yes* | *pNG6* | * |
| 168446 | 168446 | 1 | + | 43 | 168446 | Yes | - | - | - |

**Supplementary Table 3. Details of 28 intergenic TSS clusters.** Columns 1-6 are details of CAGE-seq peaks as detected CAGEfightR analysis and TSS location is the genome coordinate of the cluster's highest peak. RNA-seq coverage was confirmed from viewing alignments manually and whether if RNA-seq reads covered a transcript downstream of TSS and if covered regions could encode ORFs with NCBI ORF Finder [7]. Comments include noteworthy features of potential transcripts arising from 28 intergenic TSSs.

| Gene Name | DESeq2 Basemean | log2 Fold Change | lfcSE | Adjusted *p*-value | S1-5h FPM | S2-5h FPM | S3-16h FPM | S4-16h FPM | Function (VOCS) | Gene Type |
|---|---|---|---|---|---|---|---|---|---|---|
| A104R | 12412.91 | 7.34 | 0.31 | 5.53E-121 | 105.30 | 198.70 | 27948.49 | 21399.14 | Histone-like structural protein, IHF-like DNA-binding protein | Late |
| A118R | 80.86 | 7.91 | 1.46 | 6.87E-08 | 0.00 | 0.00 | 191.18 | 132.28 | Uncharacterised | Late |
| A125L | 3181.56 | -3.26 | 0.20 | 1.13E-57 | 6066.26 | 5457.26 | 517.16 | 685.55 | MGF 360-9L | Early |
| A137R | 72167.18 | 7.48 | 0.60 | 1.91E-35 | 492.67 | 1117.71 | 164442.30 | 122616.06 | p11.5 | Late |
| A151R | 78164.82 | -3.41 | 0.16 | 2.37E-94 | 145808.39 | 139901.71 | 15031.84 | 11917.33 | CXXC-motif containing protein | Early |
| A179L | 11468.43 | -3.89 | 0.08 | 0.00E+00 | 22027.33 | 20938.42 | 1434.09 | 1473.87 | bcl-2-bax homolog | Early |
| A224L | 3487.03 | 4.68 | 0.25 | 1.21E-80 | 259.50 | 262.57 | 7717.58 | 5708.46 | IAP homolog | Late |
| A238L | 4120.89 | -4.18 | 0.19 | 3.13E-112 | 8089.60 | 7533.01 | 375.82 | 485.12 | IkB-like protein | Early |
| A240L | 2427.65 | -6.89 | 1.15 | 2.24E-09 | 5073.39 | 4556.00 | 10.89 | 70.30 | Thymidylate kinase | Early |
| A276R | 6897.81 | -5.00 | 0.14 | 3.12E-263 | 14479.30 | 12277.06 | 429.74 | 405.16 | MGF 360-15R | Early |
| A280R | 5044.40 | -3.03 | 0.12 | 2.44E-141 | 9300.60 | 8679.10 | 1153.59 | 1044.29 | MGF 505-3R | Early |
| A489R | 243.66 | 6.46 | 0.84 | 1.74E-14 | 3.76 | 7.10 | 490.47 | 473.31 | MGF 505-2R | Late |
| A498R | 734.79 | -4.82 | 0.14 | 8.70E-248 | 1447.93 | 1390.93 | 50.38 | 49.91 | MGF 505-5R | Early |
| A505R | 268.97 | -2.22 | 0.15 | 8.20E-51 | 417.46 | 468.37 | 96.95 | 93.11 | MGF 505-4R | Early |
| A506R | 1651.43 | -3.48 | 0.15 | 8.38E-117 | 3294.51 | 2767.66 | 268.79 | 274.76 | MGF 505-9R | Early |
| A528R | 1318.63 | -3.10 | 0.16 | 4.19E-81 | 2598.75 | 2125.42 | 274.78 | 275.56 | MGF 505-6R/7R | Early |
| A542R | 2945.95 | -5.01 | 0.19 | 4.26E-147 | 6231.74 | 5198.23 | 158.22 | 195.60 | MGF 505-10R | Early |
| A859L | 121.97 | 2.42 | 0.33 | 2.95E-13 | 37.61 | 39.03 | 211.06 | 200.16 | Helicase superfamily II | Late |
| B117L | 1496.91 | 7.21 | 0.46 | 1.38E-54 | 7.52 | 31.93 | 3122.27 | 2825.92 | TR containing protein | Late |
| B119L | 3694.37 | 7.04 | 0.79 | 6.52E-19 | 26.33 | 85.16 | 6757.07 | 7908.93 | FAD-dependent thiol oxidase; ALR/ERV-like region | Late |
| B125R | 2220.97 | 5.64 | 0.24 | 2.85E-119 | 71.46 | 102.90 | 4275.59 | 4433.95 | E2 early regulatory protein | Late |
| B169L | 284.58 | 1.25 | 0.20 | 2.19E-10 | 188.04 | 149.03 | 421.02 | 380.21 | PSP, TR and Bacteriocin AS-48 | Late |
| B175L | 517.84 | 6.84 | 1.34 | 3.64E-07 | 0.00 | 17.74 | 1141.07 | 912.55 | Late TF VLTF-2 | Late |
| B263R | 3529.89 | -3.56 | 0.12 | 1.87E-206 | 6525.09 | 6489.81 | 581.70 | 522.95 | TBP | Early |
| B318L | 273.49 | 6.63 | 0.84 | 3.81E-15 | 0.00 | 10.64 | 555.28 | 528.05 | Prenyltransferase | Late |
| B354L | 486.65 | 5.57 | 0.46 | 1.28E-33 | 22.57 | 17.74 | 1011.98 | 894.30 | P-loop-containing nucleoside triphosphate hydrolases | Late |

| Gene Name | DESeq2 Basemean | log2 Fold Change | lfcSE | Adjusted *p*-value | S1-5h FPM | S2-5h FPM | S3-16h FPM | S4-16h FPM | Function (VOCS) | Gene Type |
|---|---|---|---|---|---|---|---|---|---|---|
| B385R | 647.95 | 6.68 | 1.74 | 1.29E-04 | 0.00 | 24.84 | 1162.85 | 1404.11 | A2L-like TF | Late |
| B407L | 1223.70 | 8.06 | 0.67 | 7.02E-33 | 7.52 | 10.64 | 2674.83 | 2201.81 | Uncharacterised | Late |
| B438L | 723.05 | 7.62 | 0.74 | 9.65E-25 | 0.00 | 14.19 | 1531.32 | 1346.69 | p49 | Late |
| B475L | 3255.55 | 6.79 | 0.29 | 2.72E-117 | 52.65 | 63.87 | 6836.86 | 6068.81 | Uncharacterised | Late |
| B602L | 1779.52 | 6.27 | 0.38 | 8.70E-62 | 30.09 | 60.32 | 3033.22 | 3994.44 | Chaperone | Late |
| B646L | 10645.63 | 7.45 | 0.69 | 4.89E-27 | 63.93 | 177.41 | 20599.64 | 21741.51 | p72 | Late |
| B66L | 1006.17 | 5.91 | 0.39 | 1.94E-51 | 22.57 | 42.58 | 2165.30 | 1794.24 | PSP | Late |
| B962L | 220.71 | 6.32 | 0.84 | 6.37E-14 | 0.00 | 10.64 | 431.10 | 441.11 | RNA helicase | Late |
| C105R | 2549.03 | -3.90 | 0.22 | 5.28E-67 | 5001.94 | 4552.45 | 267.43 | 374.30 | Uncharacterised | Early |
| C129R | 1440.70 | 5.96 | 0.44 | 2.14E-41 | 26.33 | 63.87 | 2246.73 | 3425.88 | Mn-dependent superoxide dismutase | Late |
| C147L | 1120.71 | 3.39 | 0.21 | 5.57E-56 | 221.89 | 170.32 | 2174.83 | 1915.79 | RPB6 | Late |
| C257L | 2402.34 | 7.21 | 1.30 | 3.02E-08 | 3.76 | 60.32 | 5034.85 | 4510.42 | Putative transmembrane domain | Late |
| C315R | 667.43 | 0.89 | 0.18 | 1.09E-06 | 458.82 | 475.47 | 944.17 | 791.27 | TFIIB | Late |
| C44L | 5000.81 | -3.06 | 0.17 | 2.43E-72 | 9545.05 | 8320.72 | 1167.75 | 969.70 | Uncharacterised | Early |
| C475L | 1315.92 | 6.67 | 0.41 | 1.75E-60 | 22.57 | 28.39 | 2665.85 | 2546.87 | E1L-like polyA polymerase large subunit | Late |
| C717R | 193.82 | 6.72 | 1.03 | 7.60E-11 | 0.00 | 7.10 | 362.74 | 405.43 | Uncharacterised | Late |
| C84L | 3371.73 | 6.72 | 0.79 | 2.83E-17 | 41.37 | 85.16 | 3936.00 | 9424.39 | PSP | Late |
| C962R | 748.25 | 8.68 | 1.04 | 8.78E-17 | 0.00 | 7.10 | 1643.52 | 1342.39 | Putative DNA primase | Late |
| CP123L | 703.85 | 7.58 | 0.75 | 1.12E-23 | 11.28 | 3.55 | 1575.71 | 1224.87 | PSP | Late |
| CP204L | 34902.57 | -4.11 | 0.13 | 6.76E-211 | 64224.14 | 67743.82 | 4158.76 | 3483.57 | Phosphoprotein | Early |
| CP2475L | 3133.75 | 5.70 | 0.33 | 5.39E-66 | 90.26 | 145.48 | 5070.25 | 7229.01 | 220kDa Polyprotein | Late |
| CP312R | 50190.58 | -2.61 | 0.17 | 1.04E-55 | 89339.14 | 83139.82 | 15858.64 | 12424.72 | Uncharacterised | Early |
| CP530R | 1934.22 | 6.22 | 0.32 | 1.63E-85 | 63.93 | 39.03 | 4060.45 | 3573.45 | 60 kDa polyprotein | Late |
| CP80R | 1958.20 | 0.81 | 0.23 | 5.94E-04 | 1462.97 | 1387.38 | 2916.93 | 2065.51 | RPB10 | Late |
| D1133L | 203.95 | 7.80 | 1.44 | 6.31E-08 | 0.00 | 3.55 | 404.41 | 407.84 | Helicase Superfamily II | Late |
| D117L | 7593.63 | 7.32 | 0.30 | 6.64E-134 | 60.17 | 127.74 | 16138.32 | 14048.31 | p17 | Late |

| Gene Name | DESeq2 Basemean | log2 Fold Change | lfcSE | Adjusted *p*-value | S1-5h FPM | S2-5h FPM | S3-16h FPM | S4-16h FPM | Function (VOCS) | Gene Type |
|---|---|---|---|---|---|---|---|---|---|---|
| D129L | 1206.85 | 6.55 | 0.43 | 9.40E-53 | 15.04 | 35.48 | 2585.51 | 2191.35 | Uncharacterised | Late |
| D205R | 315.06 | 0.47 | 0.16 | 3.91E-03 | 240.69 | 287.41 | 374.45 | 357.67 | RPB5 | Late |
| D250R | 10071.89 | -3.76 | 0.13 | 2.00E-197 | 19466.19 | 18053.67 | 1294.12 | 1473.60 | 8-Hydroxy-dGTpase Nudix hydrolase | Early |
| D339L | 4933.46 | -2.79 | 0.16 | 2.45E-66 | 8849.29 | 8398.79 | 1370.64 | 1115.13 | RPB7-fusion | Early |
| D345L | 889.03 | 1.38 | 0.20 | 1.50E-12 | 462.59 | 521.60 | 1410.13 | 1161.82 | Lambda exonuclease | Late |
| D79L | 1149.45 | 7.12 | 0.51 | 2.47E-43 | 11.28 | 21.29 | 2480.39 | 2084.83 | Uncharacterised | Late |
| DP141L | 43712.89 | -4.27 | 0.11 | 0.00E+00 | 86390.63 | 79829.28 | 4562.36 | 4069.31 | MGF 100-2L | Early |
| DP146L | 1438.92 | -3.32 | 0.12 | 8.86E-165 | 2523.53 | 2707.34 | 270.15 | 254.63 | MGF 100-3L | Early |
| DP148R | 71.52 | 7.73 | 1.45 | 1.01E-07 | 0.00 | 0.00 | 149.78 | 136.31 | MGF 360-18R | Late |
| DP238L | 18113.22 | -3.86 | 0.10 | 0.00E+00 | 35551.37 | 32236.15 | 2302.56 | 2362.81 | Uncharacterised | Early |
| DP311R | 4483.79 | -3.82 | 0.09 | 0.00E+00 | 8413.04 | 8334.92 | 595.32 | 591.91 | MGF 360-16R | Early |
| DP363R | 1282.31 | -3.57 | 0.12 | 5.03E-209 | 2410.71 | 2320.58 | 203.16 | 194.80 | MGF 360-19R | Early |
| DP42R | 1167.96 | -2.62 | 0.15 | 3.39E-64 | 2053.43 | 1965.75 | 353.21 | 299.44 | MGF 360-21R | Early |
| DP542L | 566.30 | -3.21 | 0.14 | 2.26E-121 | 1071.84 | 972.23 | 110.57 | 110.55 | MGF 505-11L | Early |
| DP60R | 80.20 | -3.26 | 0.25 | 8.17E-40 | 112.83 | 177.41 | 15.80 | 14.76 | Uncharacterised | Early |
| DP63R | 318.18 | 7.44 | 1.03 | 6.43E-13 | 3.76 | 3.55 | 675.38 | 590.03 | Uncharacterised | Late |
| DP71L | 916.95 | 2.73 | 0.19 | 1.08E-46 | 274.54 | 205.80 | 1628.81 | 1558.66 | Virulence associated protein | Late |
| DP93R | 1.15 | 1.77 | 1.49 | 2.39E-01 | 0.00 | 0.00 | 3.00 | 1.61 | Uncharacterised | NC |
| DP96R | 7901.93 | -2.46 | 0.12 | 2.60E-93 | 13569.17 | 13171.23 | 2587.41 | 2279.89 | Uncharacterised | Early |
| E111R | 1763.46 | 5.73 | 0.38 | 9.09E-50 | 37.61 | 92.26 | 3953.70 | 2970.28 | Uncharacterised | Late |
| E120R | 9090.78 | 2.65 | 0.15 | 4.84E-69 | 2391.90 | 2593.80 | 17031.57 | 14345.87 | DNA-binding p14.5 | Late |
| E146L | 1195.83 | 8.35 | 0.74 | 2.74E-29 | 3.76 | 10.64 | 2557.19 | 2211.74 | PSP | Late |
| E165R | 22775.78 | -3.95 | 0.14 | 1.02E-183 | 43735.00 | 41841.36 | 3019.60 | 2507.16 | dUTPase | Early |
| E183L | 2320.48 | 7.06 | 0.38 | 4.10E-75 | 18.80 | 49.68 | 5003.81 | 4209.64 | p54 | Late |
| E184L | 29823.34 | 7.51 | 0.52 | 3.19E-46 | 218.13 | 432.89 | 67883.89 | 50758.45 | TR containing protein | Late |
| E199L | 768.76 | 8.13 | 0.84 | 6.84E-22 | 0.00 | 10.64 | 1562.09 | 1502.31 | Vaccinia J5-like virion membrane protein | Late |

| Gene Name | DESeq2 Basemean | log2 Fold Change | lfcSE | Adjusted *p*-value | S1-5h FPM | S2-5h FPM | S3-16h FPM | S4-16h FPM | Function (VOCS) | Gene Type |
|---|---|---|---|---|---|---|---|---|---|---|
| E248R | 751.47 | 4.05 | 0.29 | 4.25E-44 | 56.41 | 113.55 | 1496.73 | 1339.17 | Putative transmembrane domain containing protein | Late |
| E296R | 787.77 | -3.02 | 0.16 | 1.07E-83 | 1470.49 | 1334.15 | 184.91 | 161.53 | AP endonuclease class II | Early |
| E301R | 92.73 | 6.65 | 1.44 | 4.45E-06 | 3.76 | 0.00 | 208.33 | 158.84 | Proliferating cell nuclear antigen-like protein | Late |
| E423R | 155.35 | 8.85 | 1.45 | 1.07E-09 | 0.00 | 0.00 | 308.28 | 313.13 | Uncharacterised | Late |
| EP1242L | 3103.38 | -0.13 | 0.13 | 3.15E-01 | 3264.42 | 3225.39 | 2772.87 | 3150.85 | RPB2 | NC |
| EP152R | 497.12 | 6.75 | 0.66 | 1.51E-24 | 7.52 | 10.64 | 1023.96 | 946.36 | PSP, TR and tRNA-guanine transglycosylase | Late |
| EP153R | 4717.39 | -3.54 | 0.12 | 2.25E-184 | 8838.01 | 8533.62 | 793.84 | 704.07 | Lectin-like protein | Early |
| EP364R | 995.14 | 6.62 | 0.49 | 1.24E-41 | 11.28 | 28.39 | 2210.78 | 1730.11 | ERCC4 domain; predicted nuclease domain; potential DEATH domain | Late |
| EP402R | 367.37 | 5.02 | 0.44 | 6.40E-30 | 30.09 | 14.19 | 757.35 | 667.84 | CD2 homolog | Late |
| EP424R | 707.89 | -1.79 | 0.17 | 1.79E-25 | 1086.89 | 1110.61 | 347.77 | 286.29 | FTS J-like Methyltransferase domain containing protein | Early |
| EP84R | 853.30 | 7.86 | 0.75 | 2.81E-25 | 3.76 | 10.64 | 1912.85 | 1485.94 | PSP | Late |
| F1055L | 315.94 | -3.08 | 0.15 | 3.35E-94 | 530.28 | 599.66 | 66.72 | 67.08 | Helicase superfamily II | Early |
| F165R | 1417.55 | 7.13 | 0.47 | 1.56E-52 | 22.57 | 17.74 | 3024.78 | 2605.10 | PSP, TR | Late |
| F317L | 1431.82 | 6.35 | 0.37 | 1.39E-64 | 26.33 | 42.58 | 3047.11 | 2611.27 | Uncharacterised | Late |
| F334L | 3114.64 | -3.25 | 0.10 | 2.34E-243 | 5573.59 | 5698.54 | 578.70 | 607.74 | Ribonucleotide reductase small subunit | Early |
| F778R | 2029.90 | -2.56 | 0.12 | 2.86E-99 | 3599.14 | 3346.03 | 563.72 | 610.69 | Ribonucleotide reductase large subunit | Early |
| G1211R | 1481.86 | -2.79 | 0.11 | 2.63E-151 | 2643.88 | 2533.47 | 373.37 | 376.72 | DNA polymerase | Early |
| G1340L | 738.32 | 3.29 | 0.21 | 6.02E-53 | 131.63 | 141.93 | 1277.50 | 1402.23 | Vaccinia A7L-like TF | Late |
| H108R | 276.04 | 5.02 | 0.50 | 2.83E-23 | 26.33 | 7.10 | 575.43 | 495.31 | PSP | Late |
| H124R | 298.16 | 5.32 | 0.52 | 2.13E-24 | 18.80 | 10.64 | 584.69 | 578.49 | Uncharacterised | Late |
| H171R | 3854.74 | 7.65 | 0.35 | 1.02E-107 | 18.80 | 56.77 | 7914.20 | 7429.18 | Uncharacterised | Late |
| H233R | 971.47 | 7.46 | 0.61 | 3.22E-34 | 3.76 | 17.74 | 2044.11 | 1820.27 | Putative transmembrane domain | Late |
| H240R | 1019.12 | 7.31 | 0.59 | 1.75E-35 | 11.28 | 14.19 | 2273.42 | 1777.60 | Uncharacterised | Late |

| Gene Name | DESeq2 Basemean | log2 Fold Change | lfcSE | Adjusted *p*-value | S1-5h FPM | S2-5h FPM | S3-16h FPM | S4-16h FPM | Function (VOCS) | Gene Type |
|---|---|---|---|---|---|---|---|---|---|---|
| H339R | 2033.61 | 7.31 | 0.40 | 4.98E-73 | 18.80 | 31.93 | 4031.31 | 4052.40 | Alpha-NAC-binding protein | Late |
| H359L | 2679.65 | -0.60 | 0.12 | 6.64E-07 | 3264.42 | 3200.55 | 2239.65 | 2013.99 | RPB3 | Early |
| I177L | 262.46 | 4.95 | 0.50 | 1.19E-22 | 7.52 | 24.84 | 547.66 | 469.82 | PSP, TR and periplasmic chaperone C-domain-containing protein | Late |
| I196L | 1605.16 | 6.06 | 0.34 | 5.51E-71 | 22.57 | 70.97 | 3288.39 | 3038.70 | PSP | Late |
| I215L | 2206.18 | -4.47 | 0.15 | 3.82E-196 | 4298.66 | 4144.39 | 207.24 | 174.41 | Ubiquitin-conjugation enzyme | Early |
| I226R | 1404.92 | 7.26 | 0.53 | 4.95E-43 | 7.52 | 28.39 | 3227.94 | 2355.83 | Uncharacterised | Late |
| I243L | 1107.43 | -0.02 | 0.21 | 9.25E-01 | 1158.34 | 1071.58 | 1246.46 | 953.33 | TFIIS | NC |
| I267L | 2660.01 | -2.40 | 0.14 | 7.10E-69 | 4776.29 | 4172.78 | 877.18 | 813.81 | RING finger containing protein | Early |
| I329L | 944.22 | 7.42 | 0.60 | 1.22E-34 | 11.28 | 10.64 | 1920.48 | 1834.49 | PSP, TR and L-domain-like-region-containing protein | Late |
| I73R | 59244.89 | -4.05 | 0.08 | 0.00E+00 | 114487.98 | 109031.65 | 6884.52 | 6575.39 | Uncharacterised | Early |
| J154R | 1417.99 | 0.31 | 0.23 | 1.68E-01 | 1289.97 | 1238.35 | 1821.89 | 1321.73 | MGF 300-2R | NC |
| J268L | 2763.65 | -4.07 | 0.15 | 8.39E-155 | 5408.11 | 5024.37 | 337.96 | 284.15 | MGF 300-1L | Early |
| J319L | 656.63 | -3.08 | 0.21 | 1.74E-47 | 1308.78 | 1039.65 | 153.59 | 124.50 | MGF 360-8L | Early |
| J328L | 938.44 | -3.37 | 0.14 | 1.93E-129 | 1812.73 | 1610.92 | 169.12 | 160.99 | MGF 300-4L | Early |
| J64R | 1343.33 | -1.52 | 0.23 | 2.74E-11 | 2136.17 | 1845.11 | 801.74 | 590.30 | Uncharacterised | Early |
| K145R | 7102.55 | 6.77 | 0.30 | 3.38E-111 | 90.26 | 166.77 | 15601.01 | 12552.17 | Uncharacterised | Late |
| K196R | 4168.53 | -2.58 | 0.17 | 2.41E-54 | 7581.89 | 6702.71 | 1303.65 | 1085.88 | Thymidine kinase | Early |
| K205R | 32788.74 | -2.12 | 0.14 | 1.58E-50 | 54603.87 | 52049.77 | 13466.76 | 11034.56 | Uncharacterised | Early |
| K421R | 268.72 | 6.60 | 0.84 | 4.48E-15 | 3.76 | 7.10 | 536.76 | 527.24 | Uncharacterised | Late |
| K78R | 28287.64 | 5.38 | 0.16 | 7.06E-239 | 1233.56 | 1419.31 | 60500.46 | 49997.23 | p10; DNA-binding activity | Late |
| KP177R | 792.48 | 6.94 | 0.59 | 3.86E-32 | 11.28 | 14.19 | 1767.97 | 1376.47 | p22 | Late |
| KP360L | 449.58 | -0.99 | 0.14 | 1.25E-12 | 567.89 | 628.05 | 292.21 | 310.18 | MGF 360-1L | Early |
| KP362L | 136.52 | -0.04 | 0.24 | 8.62E-01 | 131.63 | 145.48 | 116.56 | 152.40 | MGF 360-2L | NC |
| KP93L | 0.95 | 1.49 | 1.49 | 3.22E-01 | 0.00 | 0.00 | 1.63 | 2.15 | Uncharacterised | NC |
| L270L | 2311.46 | -6.33 | 0.16 | 0.00E+00 | 4874.07 | 4257.94 | 58.82 | 55.01 | MGF 110-1L | Early |

| Gene Name | DESeq2 Basemean | log2 Fold Change | lfcSE | Adjusted *p*-value | S1-5h FPM | S2-5h FPM | S3-16h FPM | S4-16h FPM | Function (VOCS) | Gene Type |
|---|---|---|---|---|---|---|---|---|---|---|
| L356L | 104.47 | -3.93 | 0.24 | 1.79E-59 | 221.89 | 170.32 | 11.17 | 14.49 | MGF 360-3L | Early |
| L57L | 7188.17 | 6.17 | 0.25 | 5.17E-136 | 157.96 | 234.19 | 15579.50 | 12781.04 | Uncharacterised | Late |
| L83L | 3125.37 | -3.57 | 0.11 | 1.19E-221 | 5998.57 | 5531.77 | 497.82 | 473.31 | Uncharacterised | Early |
| M1249L | 690.38 | 7.00 | 1.50 | 3.19E-06 | 0.00 | 21.29 | 1340.96 | 1399.28 | Ubiquitin-like domain containing protein | Late |
| M448R | 2486.37 | -2.98 | 0.14 | 6.09E-96 | 4614.57 | 4215.36 | 519.61 | 595.93 | Microbody targeting signal-containing protein | Early |
| NP1450L | 1790.41 | -2.31 | 0.16 | 2.46E-46 | 3106.47 | 2856.37 | 652.78 | 546.03 | RPB1 | Early |
| NP419L | 444.10 | 6.10 | 0.56 | 2.56E-27 | 3.76 | 21.29 | 912.85 | 838.49 | DNA ligase | Late |
| NP868R | 1539.58 | -1.94 | 0.15 | 7.00E-38 | 2579.95 | 2302.84 | 596.95 | 678.58 | mRNA guanylyltransferase | Early |
| O174L | 492.07 | 1.92 | 0.19 | 9.87E-23 | 203.09 | 209.35 | 827.89 | 727.95 | DNA polymerase beta-like protein | Late |
| O61R | 14672.24 | 7.49 | 0.26 | 3.06E-183 | 124.11 | 198.70 | 32360.52 | 26005.61 | p12 | Late |
| P1192R | 1351.28 | -3.24 | 0.12 | 4.60E-155 | 2335.49 | 2551.22 | 255.17 | 263.22 | Topoisomerase II | Early |
| pNG1 | 39968.54 | -3.05 | 0.11 | 2.93E-183 | 75758.69 | 66853.20 | 8724.93 | 8537.33 | Uncharacterised | Early |
| pNG2 | 31296.12 | -3.22 | 0.07 | 0.00E+00 | 57702.82 | 55374.51 | 6108.11 | 5999.05 | Uncharacterised | Early |
| pNG3 | 12216.03 | -3.78 | 0.18 | 3.69E-97 | 24761.48 | 20789.39 | 1479.85 | 1833.41 | Uncharacterised | Early |
| pNG4 | 6456.81 | -3.64 | 0.12 | 1.02E-186 | 12738.02 | 11170.00 | 942.81 | 976.41 | Uncharacterised | Early |
| pNG5 | 1432.82 | 7.15 | 0.47 | 5.45E-53 | 15.04 | 24.84 | 2647.87 | 3043.53 | Uncharacterised | Late |
| pNG6 | 1816.36 | -3.24 | 0.12 | 4.13E-162 | 3253.14 | 3317.64 | 363.56 | 331.10 | Uncharacterised | Early |
| pNG7 | 1704.79 | -3.86 | 0.17 | 1.36E-117 | 3151.60 | 3228.94 | 196.35 | 242.29 | Uncharacterised | Early |
| Q706L | 386.23 | 6.12 | 0.62 | 4.04E-23 | 3.76 | 17.74 | 687.09 | 836.35 | Helicase superfamily II | Late |
| QP383R | 29.19 | 6.44 | 1.45 | 9.05E-06 | 0.00 | 0.00 | 56.64 | 60.10 | NifS-like PLP-dependent transferase | Late |
| QP509L | 82.30 | 7.94 | 1.45 | 4.61E-08 | 0.00 | 0.00 | 162.04 | 167.16 | Helicase superfamily II | Late |
| R298L | 180.89 | 6.62 | 1.03 | 1.38E-10 | 7.52 | 0.00 | 374.18 | 341.84 | Serine protein kinase | Late |
| S183L | 523.55 | 7.15 | 0.73 | 1.63E-22 | 0.00 | 14.19 | 1039.21 | 1040.80 | Uncharacterised | Late |
| S273R | 462.74 | 7.39 | 0.85 | 5.05E-18 | 3.76 | 7.10 | 835.78 | 1004.31 | Ulp1 protease Family | Late |
| U104L | 10316.43 | -4.01 | 0.13 | 1.49E-200 | 19421.06 | 19430.40 | 1307.46 | 1106.81 | MGF 110-2L | Early |
| UP60L | 88.57 | -2.72 | 0.24 | 5.64E-30 | 173.00 | 134.83 | 19.88 | 26.56 | MGF 110-7L/MGF 360-6L | Early |

| Gene Name | DESeq2 Basemean | log2 Fold Change | lfcSE | Adjusted *p*-value | S1-5h FPM | S2-5h FPM | S3-16h FPM | S4-16h FPM | Function (VOCS) | Gene Type |
|---|---|---|---|---|---|---|---|---|---|---|
| V82L | 2086.94 | -3.36 | 0.17 | 1.28E-83 | 4035.40 | 3569.57 | 406.86 | 335.93 | MGF 110-5L | Early |
| X69R | 62.64 | -0.47 | 0.30 | 1.14E-01 | 63.93 | 81.61 | 44.12 | 60.91 | PSP, TR | Early |
| XP124L | 42504.67 | -4.18 | 0.11 | 0.00E+00 | 85025.44 | 76124.87 | 4585.23 | 4283.15 | MGF 110-3L | Early |
| Y118L | 69780.77 | -3.54 | 0.10 | 5.53E-260 | 135138.84 | 121826.76 | 11256.79 | 10900.67 | MGF 110-6L | Early |

**Supplementary Table 4. DESEq2 analysis of reads mapping to ASFV TSSs (25 nt upstream and 75 nt downstream of pTSS).** Columns 1-10 are results output from DESEq2 as described by Love *et al*. [8], lfcSE is the logarithmic fold change standard error, adjusted *p*-value (padj) is a Wald test *p*-value with Benjamini–Hochberg correction [9]. Per sample coverage is reported in fragments per million mapped reads (FPM), gene functions were from the VOCS tool database and gene type was defined as genes with a significant (padj < 0.05) downregulation (negative log2 fold change, early) or upregulation (positive log2 fold change, late). PSP and TR refer to putative signal peptide and transmembrane region, respectively.

| Gene Name | DESeq2 Basemean | log2 Fold Change | lfcSE | Adjusted *p*-value | S3-5h FPKM | S4-5h FPKM | S5-16h FPKM | S6-16h FPKM | Product Function |
|---|---|---|---|---|---|---|---|---|---|
| A104R | 1263.92 | 0.92 | 0.24 | 3.41E-04 | 5607.02 | 5080.09 | 11839.88 | 8243.62 | Histone-like structural protein, IHF-like DNA-binding protein |
| A118R | 200.73 | 6.84 | 0.91 | 3.91E-13 | 37.75 | 43.94 | 7909.57 | 1425.56 | Uncharacterised |
| A125L | 728.50 | -0.23 | 0.26 | 4.10E-01 | 2210.88 | 2138.71 | 1548.71 | 2172.16 | MGF 360-9L |
| A137R | 1432.07 | 1.66 | 1.05 | 1.49E-01 | 852.24 | 2785.18 | 8638.62 | 2860.06 | p11.5 |
| A151R | 7059.23 | -2.06 | 0.47 | 3.46E-05 | 62873.93 | 70420.66 | 23041.82 | 8926.53 | CXXC-motif containing protein |
| A179L | 1401.14 | -0.79 | 0.17 | 8.71E-06 | 21822.11 | 21661.19 | 11966.54 | 13184.19 | bcl-2-bax homolog |
| A224L | 3447.03 | -1.87 | 0.23 | 1.58E-15 | 86427.87 | 94946.15 | 27330.50 | 22304.70 | IAP homolog |
| A238L | 1759.09 | -0.82 | 0.17 | 2.47E-06 | 6605.13 | 6181.17 | 3704.98 | 3523.38 | IkB-like protein |
| A240L | 256.90 | 0.93 | 0.62 | 1.70E-01 | 452.32 | 778.06 | 1685.21 | 641.08 | Thymidylate kinase |
| A276R | 8265.85 | -0.30 | 0.31 | 3.77E-01 | 91719.46 | 79604.39 | 56885.30 | 82003.73 | MGF 360-15R |
| A280R | 5981.39 | -0.68 | 0.36 | 8.45E-02 | 52183.14 | 49601.18 | 22464.78 | 41182.89 | MGF 505-3R |
| A489R | 264.68 | 1.81 | 0.40 | 1.86E-05 | 1266.88 | 1022.00 | 3598.78 | 4426.55 | MGF 505-2R |
| A498R | 1327.01 | -0.07 | 0.22 | 7.70E-01 | 11529.90 | 8638.64 | 9035.01 | 10177.63 | MGF 505-5R |
| A505R | 1059.59 | -0.72 | 0.20 | 7.97E-04 | 22291.01 | 18538.24 | 12589.78 | 12184.64 | MGF 505-4R |
| A506R | 3109.56 | -1.10 | 0.23 | 5.77E-06 | 56728.91 | 44200.54 | 23197.92 | 24027.18 | MGF 505-9R |
| A528R | 1006.51 | -1.08 | 0.27 | 2.01E-04 | 7283.55 | 4848.44 | 3078.67 | 2659.03 | MGF 505-6R/7R |
| A542R | 5598.07 | -1.17 | 0.27 | 3.34E-05 | 30761.39 | 25668.36 | 11198.37 | 13949.04 | MGF 505-10R |
| A859L | 3026.02 | -0.14 | 0.33 | 6.89E-01 | 45091.54 | 37260.63 | 27088.56 | 47616.93 | Helicase superfamily II |
| B117L | 148.83 | 3.78 | 0.63 | 1.25E-08 | 97.13 | 221.48 | 2753.97 | 1654.67 | TR containing protein |
| B119L | 207.76 | 5.31 | 0.57 | 2.08E-19 | 143.64 | 161.00 | 7880.81 | 4175.69 | FAD-dependent thiol oxidase; ALR/ERV-like region |
| B125R | 72.77 | 3.64 | 0.61 | 1.47E-08 | 51.30 | 64.13 | 954.19 | 476.61 | E2 early regulatory protein |
| B169L | 99.03 | 2.45 | 0.67 | 5.65E-04 | 86.15 | 90.08 | 704.38 | 247.20 | PSP, TR and Bacteriocin AS-48 |
| B175L | 57.04 | 2.49 | 0.72 | 1.16E-03 | 22.45 | 85.77 | 358.08 | 256.86 | Late TF VLTF-2 |

| Gene Name | DESeq2 Basemean | log2 Fold Change | lfcSE | Adjusted *p*-value | S3-5h FPKM | S4-5h FPKM | S5-16h FPKM | S6-16h FPKM | Product Function |
|---|---|---|---|---|---|---|---|---|---|
| B263R | 899.64 | -1.67 | 0.20 | 7.87E-16 | 7576.41 | 7836.90 | 2292.22 | 2577.33 | TBP |
| B318L | 325.01 | 3.43 | 0.43 | 1.84E-14 | 118.13 | 110.72 | 1532.53 | 918.89 | Prenyltransferase |
| B354L | 750.77 | 3.39 | 0.29 | 1.19E-30 | 414.45 | 492.77 | 5630.39 | 3856.51 | P-loop-containing nucleoside triphosphate hydrolases |
| B385R | 486.13 | 5.40 | 0.53 | 3.18E-23 | 28.43 | 61.00 | 2605.64 | 1208.41 | A2L-like TF |
| B407L | 239.05 | 3.47 | 0.49 | 1.75E-11 | 60.20 | 100.70 | 1035.71 | 747.18 | Uncharacterised |
| B438L | 577.17 | 3.28 | 0.39 | 2.54E-16 | 141.71 | 269.34 | 2257.55 | 1750.58 | p49 |
| B475L | 378.50 | 2.80 | 0.49 | 5.51E-08 | 145.27 | 235.26 | 1752.81 | 890.00 | Uncharacterised |
| B602L | 1249.21 | 4.01 | 0.41 | 7.15E-21 | 198.11 | 347.82 | 6015.06 | 2758.12 | Chaperone |
| B646L | 4428.35 | 8.01 | 0.41 | 8.68E-85 | 98.86 | 197.26 | 46996.66 | 30423.70 | p72 |
| B66L | 192.78 | -2.26 | 0.45 | 1.79E-06 | 3746.95 | 3634.41 | 720.38 | 838.87 | PSP |
| B962L | 1680.97 | 5.04 | 0.21 | 1.63E-121 | 896.17 | 985.55 | 34275.87 | 27516.16 | RNA helicase |
| C105R | 451.62 | -0.65 | 0.32 | 5.79E-02 | 2181.16 | 2418.63 | 1261.64 | 1694.77 | Uncharacterised |
| C129R | 792.49 | -0.11 | 0.27 | 6.91E-01 | 3449.59 | 3874.59 | 3980.43 | 2744.96 | Mn-dependent superoxide dismutase |
| C147L | 595.75 | 0.06 | 0.30 | 8.41E-01 | 3736.26 | 4045.31 | 3300.86 | 4887.64 | RPB6 |
| C257L | 811.28 | 0.66 | 0.20 | 1.72E-03 | 2282.50 | 2231.91 | 3408.14 | 3746.42 | Putative transmembrane domain |
| C315R | 1273.01 | -0.78 | 0.18 | 3.69E-05 | 6812.14 | 6465.63 | 3636.08 | 4125.46 | TFIIB |
| C44L | 2005.29 | -1.28 | 0.42 | 4.10E-03 | 2726.24 | 3834.14 | 764.91 | 1942.88 | Uncharacterised |
| C475L | 946.30 | 1.12 | 0.29 | 2.01E-04 | 3389.10 | 3210.27 | 8842.65 | 5467.60 | E1L-like polyA polymerase large subunit |
| C62L | 74.87 | 0.11 | 0.52 | 8.41E-01 | 428.55 | 607.88 | 630.72 | 476.46 | Uncharacterised |
| C717R | 383.28 | 2.08 | 0.30 | 4.96E-11 | 465.63 | 466.90 | 1932.92 | 2019.49 | Uncharacterised |
| C84L | 278.63 | 0.39 | 0.57 | 5.39E-01 | 760.98 | 736.19 | 1415.54 | 526.99 | PSP |
| C962R | 1648.22 | 1.88 | 0.22 | 5.71E-17 | 775.87 | 1045.51 | 3271.96 | 3450.45 | Putative DNA primase |
| CP123L | 198.18 | 1.28 | 0.56 | 3.54E-02 | 983.89 | 724.93 | 2797.27 | 1318.42 | PSP |
| CP204L | 19125.48 | -0.87 | 0.30 | 7.04E-03 | 22176.76 | 25380.58 | 10044.78 | 16055.92 | Phosphoprotein |

| Gene Name | DESeq2 Basemean | log2 Fold Change | lfcSE | Adjusted *p*-value | S3-5h FPKM | S4-5h FPKM | S5-16h FPKM | S6-16h FPKM | Product Function |
|---|---|---|---|---|---|---|---|---|---|
| CP2475L | 9563.70 | 1.12 | 0.28 | 1.41E-04 | 32544.40 | 24550.74 | 60595.70 | 63723.22 | 220kDa Polyprotein |
| CP312R | 22673.02 | -0.93 | 0.20 | 7.18E-06 | 354268.37 | 397366.32 | 184055.78 | 211797.71 | Uncharacterised |
| CP530R | 920.44 | 5.20 | 0.31 | 2.82E-63 | 202.51 | 289.36 | 10758.25 | 7429.95 | 60 kDa polyprotein |
| CP80R | 88.36 | 0.50 | 0.53 | 3.85E-01 | 541.84 | 476.74 | 880.66 | 539.08 | RPB10 |
| D1133L | 1224.92 | 4.07 | 0.26 | 1.03E-52 | 350.87 | 313.85 | 6645.29 | 4454.14 | Helicase Superfamily II |
| D117L | 892.77 | 4.65 | 0.33 | 1.41E-44 | 66.59 | 45.77 | 1709.47 | 1086.61 | p17 |
| D129L | 614.32 | -0.16 | 0.28 | 6.12E-01 | 7059.95 | 7026.98 | 5203.96 | 7469.65 | Uncharacterised |
| D205R | 406.76 | -0.68 | 0.44 | 1.57E-01 | 1056.08 | 1324.06 | 509.83 | 995.13 | RPB5 |
| D250R | 2924.38 | -0.81 | 0.19 | 3.93E-05 | 23773.72 | 24523.52 | 13192.48 | 14401.49 | 8-Hydroxy-dGTpase Nudix hydrolase |
| D339L | 1715.94 | -0.56 | 0.16 | 8.16E-04 | 9975.15 | 9950.31 | 6713.32 | 6794.89 | RPB7-fusion |
| D345L | 1577.47 | -0.62 | 0.21 | 5.19E-03 | 4492.66 | 4973.32 | 2696.95 | 3479.74 | Lambda exonuclease |
| D79L | 99.79 | -0.57 | 0.84 | 5.39E-01 | 257.37 | 370.92 | 354.21 | 60.50 | Uncharacterised |
| DP141L | 5999.33 | -2.12 | 0.41 | 9.63E-07 | 7693.12 | 8167.13 | 2485.29 | 1157.21 | MGF 100-2L |
| DP146L | 1222.60 | -1.52 | 0.19 | 2.52E-15 | 4314.35 | 3851.26 | 1471.94 | 1367.70 | MGF 100-3L |
| DP238L | 2160.35 | -1.27 | 0.19 | 4.96E-11 | 35651.15 | 41360.07 | 15572.34 | 16300.55 | Uncharacterised |
| DP311R | 1165.81 | -0.60 | 0.21 | 8.12E-03 | 13399.66 | 11813.93 | 7438.12 | 9293.48 | MGF 360-16R |
| DP363R | 1342.45 | -0.27 | 0.21 | 2.34E-01 | 2302.35 | 1848.79 | 1829.30 | 1607.13 | MGF 360-19R |
| DP42R | 550.10 | -0.92 | 0.26 | 9.18E-04 | 5350.16 | 5694.67 | 2655.11 | 3200.97 | MGF 360-21R |
| DP542L | 4634.68 | -0.49 | 0.26 | 7.91E-02 | 22352.16 | 19709.54 | 13124.14 | 16918.15 | MGF 505-11L |
| DP60R | 455.17 | -1.07 | 0.34 | 2.97E-03 | 1181.90 | 1409.94 | 517.30 | 732.86 | Uncharacterised |
| DP63R | 111.33 | -1.55 | 0.50 | 3.73E-03 | 512.28 | 547.24 | 201.36 | 156.60 | Uncharacterised |
| DP71L | 132.77 | 1.14 | 0.54 | 5.14E-02 | 626.60 | 493.02 | 899.62 | 1595.57 | Virulence associated protein |
| DP96R | 2607.42 | 0.10 | 0.44 | 8.31E-01 | 2585.09 | 2705.29 | 1517.24 | 4170.28 | Uncharacterised |
| E111R | 883.40 | -1.19 | 0.81 | 1.75E-01 | 10133.96 | 9186.72 | 6516.53 | 1933.29 | Uncharacterised |
| E120R | 1278.98 | 0.31 | 0.18 | 1.06E-01 | 3412.48 | 3082.47 | 4159.94 | 3870.14 | DNA-binding p14.5 |

| Gene Name | DESeq2 Basemean | log2 Fold Change | lfcSE | Adjusted *p*-value | S3-5h FPKM | S4-5h FPKM | S5-16h FPKM | S6-16h FPKM | Product Function |
|---|---|---|---|---|---|---|---|---|---|
| E146L | 116.36 | 4.51 | 0.80 | 7.91E-08 | 8.90 | 35.50 | 708.21 | 320.18 | PSP |
| E165R | 5216.83 | -1.04 | 0.49 | 4.82E-02 | 36361.97 | 46600.86 | 10661.22 | 29689.99 | dUTPase |
| E183L | 306.57 | 3.78 | 0.53 | 1.28E-11 | 65.89 | 111.85 | 1616.36 | 820.78 | p54 |
| E184L | 518.07 | 2.49 | 0.64 | 2.50E-04 | 187.13 | 707.06 | 3460.80 | 1550.54 | TR containing protein |
| E199L | 73.84 | 2.53 | 0.54 | 1.09E-05 | 30.17 | 43.20 | 251.45 | 169.52 | Vaccinia J5-like virion membrane protein |
| E248R | 1290.76 | -0.53 | 0.18 | 6.39E-03 | 4226.54 | 3771.06 | 2633.14 | 2906.22 | Putative transmembrane domain containing protein |
| E296R | 4500.14 | -0.46 | 0.31 | 1.72E-01 | 8661.86 | 7704.58 | 4599.44 | 7298.04 | AP endonuclease class II |
| E301R | 181.49 | 3.35 | 0.51 | 3.91E-10 | 191.64 | 328.23 | 2899.37 | 2421.61 | Proliferating cell nuclear antigen-like protein |
| E423R | 199.37 | 3.58 | 0.55 | 4.60E-10 | 220.57 | 306.65 | 4122.03 | 2181.44 | Uncharacterised |
| E66L | 38.41 | -0.46 | 0.48 | 3.77E-01 | 100.55 | 121.78 | 86.82 | 71.75 | Uncharacterised |
| EP1242L | 4939.44 | -0.16 | 0.29 | 6.12E-01 | 29874.62 | 29007.90 | 21357.16 | 31328.41 | RPB2 |
| EP152R | 48.72 | 0.27 | 0.49 | 6.12E-01 | 174.28 | 153.39 | 165.65 | 239.39 | PSP, TR and tRNA-guanine transglycosylase |
| EP153R | 19.58 | 0.76 | 0.52 | 1.79E-01 | 126.33 | 295.35 | 324.01 | 436.88 | Lectin-like protein |
| EP364R | 1291.15 | 1.04 | 0.24 | 4.67E-05 | 757.42 | 523.52 | 1355.63 | 1276.05 | ERCC4 domain; predicted nuclease domain; potential DEATH domain |
| EP402R | 4410.73 | 0.92 | 0.39 | 2.73E-02 | 2982.96 | 2133.48 | 3322.29 | 6381.65 | CD2 homolog |
| EP424R | 1506.60 | -0.01 | 0.22 | 9.56E-01 | 11979.21 | 12284.09 | 13869.22 | 10098.84 | FTS J-like Methyltransferase domain containing protein |
| EP84R | 56.78 | 7.76 | 1.21 | 7.53E-10 | 0.53 | 0.00 | 65.37 | 27.32 | PSP |
| F1055L | 2616.04 | -0.18 | 0.18 | 3.53E-01 | 14864.46 | 13611.86 | 12622.87 | 12437.78 | Helicase superfamily II |
| F165R | 318.63 | 0.88 | 0.35 | 1.72E-02 | 420.04 | 433.16 | 844.52 | 726.69 | PSP, TR |
| F317L | 642.03 | 0.81 | 0.31 | 1.48E-02 | 4365.58 | 6242.74 | 10642.59 | 7968.76 | Uncharacterised |
| F334L | 3409.92 | -1.07 | 0.23 | 1.48E-05 | 16188.28 | 13986.41 | 6438.97 | 7964.17 | Ribonucleotide reductase small subunit |
| F778R | 3781.69 | -0.57 | 0.31 | 8.45E-02 | 25456.73 | 26425.29 | 13311.73 | 21671.70 | Ribonucleotide reductase large subunit |
| G1211R | 5570.60 | -0.67 | 0.33 | 6.19E-02 | 104321.77 | 108312.60 | 50457.87 | 83566.44 | DNA polymerase |

| Gene Name | DESeq2 Basemean | log2 Fold Change | lfcSE | Adjusted *p*-value | S3-5h FPKM | S4-5h FPKM | S5-16h FPKM | S6-16h FPKM | Product Function |
|---|---|---|---|---|---|---|---|---|---|
| G1340L | 886.08 | 0.53 | 0.27 | 6.91E-02 | 477.59 | 540.53 | 871.25 | 588.87 | Vaccinia A7L-like TF |
| H108R | 85.60 | 1.72 | 0.71 | 2.44E-02 | 71.27 | 122.23 | 470.35 | 160.20 | PSP |
| H124R | 178.46 | 1.96 | 0.75 | 1.48E-02 | 57.91 | 113.32 | 517.37 | 145.39 | Uncharacterised |
| H171R | 448.97 | 1.97 | 0.39 | 1.50E-06 | 692.40 | 717.34 | 3527.45 | 1985.88 | Uncharacterised |
| H233R | 540.57 | 2.30 | 0.25 | 6.30E-19 | 1469.77 | 1389.82 | 7348.70 | 6697.88 | Putative transmembrane domain |
| H240R | 393.07 | 2.20 | 0.47 | 1.07E-05 | 1646.17 | 1734.17 | 10700.28 | 4794.63 | Uncharacterised |
| H339R | 1117.24 | 1.90 | 0.30 | 1.25E-09 | 1158.67 | 1648.35 | 6355.06 | 4132.64 | Alpha-NAC-binding protein |
| H359L | 1872.34 | -1.16 | 0.19 | 4.35E-09 | 2258.13 | 2366.22 | 940.57 | 1140.03 | RPB3 |
| I177L | 132.99 | 1.42 | 0.61 | 3.16E-02 | 694.37 | 486.15 | 2171.26 | 962.29 | PSP, TR and periplasmic chaperone C-domain-containing protein |
| I196L | 404.43 | 1.14 | 0.41 | 8.57E-03 | 1205.91 | 1194.37 | 3391.13 | 1880.27 | PSP |
| I215L | 4665.43 | -0.40 | 0.28 | 1.82E-01 | 13896.29 | 14503.33 | 8844.43 | 12672.25 | Ubiquitin-conjugation enzyme |
| I226R | 76.82 | 0.86 | 0.80 | 3.32E-01 | 199.10 | 399.79 | 864.24 | 204.13 | Uncharacterised |
| I243L | 1111.38 | -0.39 | 0.25 | 1.54E-01 | 4388.67 | 4798.17 | 2826.63 | 4201.33 | TFIIS |
| I267L | 4538.64 | -0.85 | 0.35 | 2.32E-02 | 4701.55 | 5236.32 | 1971.42 | 3545.24 | RING finger containing protein |
| I329L | 2601.08 | -1.52 | 0.27 | 9.65E-08 | 10894.17 | 10755.98 | 4581.33 | 2928.12 | PSP, TR and L-domain-like-region-containing protein |
| I73R | 8083.83 | -2.21 | 0.44 | 2.29E-06 | 66941.30 | 87158.31 | 22999.62 | 10351.65 | Uncharacterised |
| J154R | 834.17 | -0.96 | 0.36 | 1.21E-02 | 8432.28 | 9361.33 | 6071.40 | 2980.44 | MGF 300-2R |
| J268L | 1219.57 | -0.64 | 0.17 | 4.97E-04 | 4606.71 | 4260.72 | 2840.78 | 2860.96 | MGF 300-1L |
| J319L | 930.92 | -0.60 | 0.19 | 3.61E-03 | 10932.74 | 9703.95 | 6785.65 | 6778.16 | MGF 360-8L |
| J328L | 817.46 | -0.17 | 0.21 | 4.66E-01 | 3434.69 | 3424.21 | 2838.40 | 3302.15 | MGF 300-4L |
| J64R | 353.93 | -1.35 | 0.64 | 5.02E-02 | 2033.64 | 2273.66 | 1334.05 | 329.56 | Uncharacterised |
| K145R | 849.64 | 0.46 | 0.33 | 1.96E-01 | 2364.67 | 2415.17 | 4270.08 | 2288.30 | Uncharacterised |
| K196R | 4882.28 | 0.10 | 0.22 | 6.64E-01 | 7035.62 | 6447.07 | 7385.90 | 7088.93 | Thymidine kinase |
| K205R | 13843.27 | -1.53 | 0.31 | 4.03E-06 | 36649.95 | 40650.22 | 10470.11 | 16392.22 | Uncharacterised |

| Gene Name | DESeq2 Basemean | log2 Fold Change | lfcSE | Adjusted *p*-value | S3-5h FPKM | S4-5h FPKM | S5-16h FPKM | S6-16h FPKM | Product Function |
|---|---|---|---|---|---|---|---|---|---|
| K421R | 796.19 | 0.46 | 0.25 | 8.66E-02 | 1828.52 | 1620.57 | 2034.18 | 2732.58 | Uncharacterised |
| K78R | 947.35 | 1.03 | 0.26 | 2.12E-04 | 3254.43 | 3447.72 | 8213.82 | 5463.04 | p10; DNA-binding activity |
| KP177R | 164.43 | 0.55 | 0.57 | 3.77E-01 | 735.27 | 710.95 | 1442.39 | 654.85 | p22 |
| KP360L | 354.62 | -0.93 | 0.59 | 1.49E-01 | 1328.17 | 901.59 | 873.99 | 283.34 | MGF 360-1L |
| KP362L | 216.68 | 1.16 | 0.62 | 8.45E-02 | 444.18 | 462.36 | 1494.23 | 516.25 | MGF 360-2L |
| KP93L | 1.65 | -3.19 | 1.50 | 5.00E-02 | 22.06 | 10.19 | 0.00 | 0.00 | Uncharacterised |
| L270L | 4621.35 | -1.27 | 0.26 | 3.89E-06 | 39256.97 | 40155.19 | 13957.22 | 18915.53 | MGF 110-1L |
| L356L | 661.85 | 1.25 | 0.97 | 2.34E-01 | 1569.91 | 1625.29 | 6204.14 | 1378.52 | MGF 360-3L |
| L57L | 138.36 | 1.79 | 0.63 | 8.12E-03 | 506.76 | 531.21 | 2564.81 | 983.21 | Uncharacterised |
| L83L | 257.89 | -0.48 | 0.90 | 6.27E-01 | 969.51 | 1075.95 | 1325.77 | 127.61 | Uncharacterised |
| M1249L | 1578.68 | 2.78 | 0.18 | 1.76E-51 | 2294.83 | 2630.73 | 17694.82 | 16122.76 | Ubiquitin-like domain containing protein |
| M448R | 4026.76 | -0.60 | 0.25 | 2.49E-02 | 75734.24 | 72012.47 | 42318.18 | 55508.65 | Microbody targeting signal-containing protein |
| NP1450L | 5337.12 | 0.59 | 0.23 | 1.56E-02 | 18057.77 | 18876.76 | 26075.03 | 29458.13 | RPB1 |
| NP419L | 2101.55 | -0.87 | 0.26 | 1.55E-03 | 16595.53 | 18037.02 | 7509.16 | 11483.45 | DNA ligase |
| NP868R | 2486.35 | -0.23 | 0.24 | 3.87E-01 | 14703.13 | 12621.72 | 9884.80 | 13502.99 | mRNA guanylyltransferase |
| O174L | 1192.72 | -1.21 | 0.38 | 3.24E-03 | 6054.41 | 7124.70 | 3984.61 | 1680.39 | DNA polymerase beta-like protein |
| O61R | 420.87 | 1.13 | 0.30 | 3.28E-04 | 870.15 | 837.29 | 2023.72 | 1712.57 | p12 |
| P1192R | 3070.69 | -0.41 | 0.34 | 2.68E-01 | 36824.00 | 36755.39 | 19266.58 | 36303.83 | Topoisomerase II |
| pNG1 | 3595.43 | -1.97 | 0.31 | 7.53E-10 | 16777.25 | 21526.79 | 5900.89 | 3827.67 | Uncharacterised |
| pNG6 | 662.05 | -0.48 | 0.29 | 1.28E-01 | 3429.62 | 3374.58 | 1982.71 | 2950.19 | Uncharacterised |
| pNG2 | 3836.43 | -1.69 | 0.35 | 6.68E-06 | 20765.66 | 28247.40 | 5517.20 | 9746.19 | Uncharacterised |
| pNG3 | 844.55 | -1.61 | 0.25 | 3.69E-10 | 13032.84 | 16201.42 | 5244.75 | 4246.57 | Uncharacterised |
| pNG4 | 1529.09 | -1.66 | 0.25 | 1.04E-10 | 5956.01 | 8249.76 | 2474.80 | 2007.27 | Uncharacterised |
| Q706L | 269.21 | 2.50 | 0.47 | 4.08E-07 | 129.59 | 172.25 | 1063.31 | 641.91 | Helicase superfamily II |
| QP383R | 284.72 | 1.98 | 0.62 | 2.75E-03 | 356.01 | 834.78 | 3199.49 | 1495.75 | NifS-like PLP-dependent transferase |

| Gene Name | DESeq2 Basemean | log2 Fold Change | lfcSE | Adjusted *p*-value | S3-5h FPKM | S4-5h FPKM | S5-16h FPKM | S6-16h FPKM | Product Function |
|---|---|---|---|---|---|---|---|---|---|
| QP509L | 184.71 | 2.87 | 0.65 | 3.34E-05 | 149.65 | 455.39 | 1992.68 | 2453.56 | Helicase superfamily II |
| R298L | 217.24 | 2.19 | 0.53 | 9.40E-05 | 268.77 | 371.45 | 1909.61 | 1006.87 | Serine protein kinase |
| S183L | 61.81 | 1.24 | 0.53 | 2.81E-02 | 139.07 | 89.93 | 236.96 | 308.06 | Uncharacterised |
| S273R | 119.41 | 1.56 | 0.47 | 1.85E-03 | 564.62 | 670.78 | 1923.96 | 1711.85 | Ulp1 protease Family |
| U104L | 7156.39 | -1.46 | 0.32 | 1.30E-05 | 26597.39 | 30972.65 | 8445.88 | 12550.27 | MGF 110-2L |
| UP60L | 140.41 | -1.13 | 0.52 | 4.51E-02 | 1579.87 | 1970.79 | 982.72 | 609.53 | MGF 110-7L/MGF 360-6L |
| V82L | 2897.02 | -0.53 | 0.38 | 2.00E-01 | 76528.52 | 83848.06 | 34791.50 | 76640.90 | MGF 110-5L |
| X69R | 1619.32 | -1.37 | 0.25 | 1.21E-07 | 31601.16 | 38287.03 | 15855.77 | 11074.39 | PSP, TR |
| XP124L | 13515.91 | -0.95 | 0.49 | 7.50E-02 | 317126.22 | 336671.16 | 87155.14 | 252147.72 | MGF 110-3L |
| Y118L | 12028.94 | -0.92 | 0.42 | 4.42E-02 | 84305.10 | 107054.38 | 32494.35 | 68996.31 | MGF 110-6L |

**Supplementary Table 5. DESEq2 analysis of reads mapping to ASFV Transcription Units.** DESeq2 results output as described for Supplementary Table 4.

| Gene Name | Gene Product | Functional Group | Gene Type | In Viral Particle |
|---|---|---|---|---|
| A104R | Histone-like structural protein, IHF-like DNA-binding protein | Genome Organisation | Late | YES |
| A118R | Uncharacterised | Uncharacterised | Late | No |
| A151R | CXXC-motif containing protein | Structural / Viral Morphology | Early | No |
| A179L | bcl-2-bax homolog | Immune Evasion | Early | No |
| A238L | IkB-like protein | Immune Evasion | Early | No |
| A489R | MGF 505-2R | MGF 505 | Late | No |
| A505R | MGF 505-4R | MGF 505 | Early | No |
| A506R | MGF 505-9R | MGF 505 | Early | No |
| A528R | MGF 505-6R/7R | MGF 505 | Early | No |
| A542R | MGF 505-10R | MGF 505 | Early | No |
| B117L | Transmembrane region containing protein | TR / PSP | Late | YES |
| B119L | FAD-dependent thiol oxidase; ALR/ERV-like region | Other / Enzyme | Late | YES |
| B125R | E2 early regulatory protein | Transcription / RNA modification | Late | No |
| B169L | Putative signal peptide, transmembrane region and Bacteriocin AS-48 | TR / PSP | Late | YES |
| B175L | Late TF VLTF-2 | Transcription / RNA modification | Late | No |
| B263R | TBP | Transcription / RNA modification | Early | No |
| B318L | Prenyltransferase | Other / Enzyme | Late | No |
| B354L | P-loop-containing nucleoside triphosphate hydrolases | Other / Enzyme | Late | No |
| B385R | A2L-like TF | Transcription / RNA modification | Late | No |
| B407L | Uncharacterised | Uncharacterised | Late | No |
| B438L | p49 | Structural / Viral Morphology | Late | YES |
| B475L | Uncharacterised | Uncharacterised | Late | No |
| B602L | Chaperone | Structural / Viral Morphology | Late | No |
| B646L | p72 | Structural / Viral Morphology | Late | YES |
| B962L | RNA helicase | NA Metabolism / DNA Replication / Repair | Late | YES |
| C257L | Putative transmembrane domain | TR / PSP | Late | YES |
| C44L | Uncharacterised | Uncharacterised | Early | No |
| C475L | E1L-like polyA polymerase large subunit | Transcription / RNA modification | Late | YES |

| Gene Name | Gene Product | Functional Group | Gene Type | In Viral Particle |
|---|---|---|---|---|
| C717R | Uncharacterised | Uncharacterised | Late | YES |
| C962R | Putative DNA primase | NA Metabolism / DNA Replication / Repair | Late | No |
| CP123L | Putative signal peptide | TR / PSP | Late | YES |
| CP204L | Phosphoprotein | Other / Enzyme | Early | YES |
| CP2475L | 220kDa Polyprotein | Structural / Viral Morphology | Late | YES |
| CP312R | Uncharacterised | Uncharacterised | Early | YES |
| CP530R | 60 kDa polyprotein | Structural / Viral Morphology | Late | YES |
| D1133L | Helicase Superfamily II | Transcription / RNA modification | Late | YES |
| D117L | p17 | Structural / Viral Morphology | Late | YES |
| D250R | 8-Hydroxy-dGTpase Nudix hydrolase | Transcription / RNA modification | Early | No |
| D339L | RPB7-fusion | Transcription / RNA modification | Early | YES |
| DP141L | MGF 100-2L | MGF 100 | Early | No |
| DP146L | MGF 100-3L | MGF 100 | Early | No |
| DP238L | Uncharacterised | Uncharacterised | Early | No |
| DP311R | MGF 360-16R | MGF 360 | Early | No |
| DP42R | MGF 360-21R | MGF 360 | Early | No |
| DP60R | Uncharacterised | Uncharacterised | Early | No |
| E146L | Putative signal peptide | TR / PSP | Late | YES |
| E165R | dUTPase | NA Metabolism / DNA Replication / Repair | Early | YES |
| E183L | p54 | Structural / Viral Morphology | Late | YES |
| E184L | Transmembrane region containing protein | TR / PSP | Late | YES |
| E199L | Vaccinia J5-like virion membrane protein | Structural / Viral Morphology | Late | YES |
| E301R | Proliferating cell nuclear antigen-like protein | NA Metabolism / DNA Replication / Repair | Late | No |
| E423R | Uncharacterised | Uncharacterised | Late | YES |
| EP364R | ERCC4 domain; predicted nuclease domain; potential DEATH domain | NA Metabolism / DNA Replication / Repair | Late | No |
| EP402R | CD2 homolog | Immune Evasion | Late | YES |
| EP84R | Putative signal peptide | TR / PSP | Late | YES |
| F165R | Putative signal peptide, transmembrane region | TR / PSP | Late | No |

| Gene Name | Gene Product | Functional Group | Gene Type | In Viral Particle |
|---|---|---|---|---|
| F317L | Uncharacterised | Uncharacterised | Late | YES |
| F334L | Ribonucleotide reductase small subunit | NA Metabolism / DNA Replication / Repair | Early | No |
| H108R | Putative signal peptide | TR / PSP | Late | YES |
| H124R | Uncharacterised | Uncharacterised | Late | YES |
| H171R | Uncharacterised | Uncharacterised | Late | YES |
| H233R | Putative transmembrane domain | TR / PSP | Late | No |
| H240R | Uncharacterised | Uncharacterised | Late | YES |
| H339R | Alpha-NAC-binding protein | Other / Enzyme | Late | YES |
| H359L | RPB3 | Transcription / RNA modification | Early | YES |
| I177L | Putative signal peptide, transmembrane region and periplasmic chaperone C-domain-containing protein | TR / PSP | Late | YES |
| I196L | Putative signal peptide | TR / PSP | Late | No |
| I267L | RING finger containing protein | Structural / Viral Morphology | Early | No |
| I73R | Uncharacterised | Uncharacterised | Early | YES |
| J268L | MGF 300-1L | MGF 300 | Early | No |
| J319L | MGF 360-8L | MGF 360 | Early | No |
| K205R | Uncharacterised | Uncharacterised | Early | No |
| K78R | p10; DNA-binding activity | Structural / Viral Morphology | Late | YES |
| L270L | MGF 110-1L | MGF 110 | Early | No |
| L57L | Uncharacterised | Uncharacterised | Late | No |
| M1249L | Ubiquitin-like domain containing protein | Other / Enzyme | Late | YES |
| M448R | Microbody targeting signal-containing protein | Other / Enzyme | Early | YES |
| pNG1 | Uncharacterised | Uncharacterised | Early | NA |
| pNG2 | Uncharacterised | Uncharacterised | Early | NA |
| pNG3 | Uncharacterised | Uncharacterised | Early | NA |
| pNG4 | Uncharacterised | Uncharacterised | Early | NA |
| O61R | p12 | Structural / Viral Morphology | Late | YES |
| Q706L | Helicase superfamily II | Transcription / RNA modification | Late | YES |
| QP383R | NifS-like PLP-dependent transferase | Other / Enzyme | Late | YES |

| Gene Name | Gene Product | Functional Group | Gene Type | In Viral Particle |
|---|---|---|---|---|
| QP509L | Helicase superfamily II | NA Metabolism / DNA Replication / Repair | Late | No |
| R298L | Serine protein kinase | NA Metabolism / DNA Replication / Repair | Late | YES |
| S183L | Uncharacterised | Uncharacterised | Late | No |
| S273R | Ulp1 protease Family | Structural / Viral Morphology | Late | YES |
| U104L | MGF 110-2L | MGF 110 | Early | No |
| UP60L | MGF 110-7L/MGF 360-6L | MGF 110-360 | Early | No |
| Y118L | MGF 110-6L | MGF 110 | Early | No |

**Supplementary Table 6. Details of 91 genes with matching differential expression patterns between CAGE-seq and RNA-seq.** Functional groups were broad groups encompassing the VOCS database functions in column 2 and according to these placed into broad functional groups (column 3). Presence in viral particle (column 5) from Alejo *et al.* [10].

| Gene | TSS sequence | Stage | Extensions in the early stage (more than 5% of total reads and at least 5 reads) | Extensions in the late stage (more than 5% of total reads and at least 5 reads) | Total reads comprising TSS in the early stage | Total reads comprising TSS in the late stage |
|---|---|---|---|---|---|---|
| pNG10 | TAATATTTTT A AATGCAACCA | Late | A (31.3%), AA (21.8%), AAA (6.2%) | A (28.6%), AA (17.7%), AAA (5.4%) | 1871 | 2498 |
| F317L | TTTGTTAAGA A TAAATGGTTG | Late | | ATA (25.2%), AT (13.7%), ATATA (9.1%) | 18 | 20816 |
| pNG11 | GAGAAATGTT A AATGAGTGAA | Late | A (30.2%), GAA (18.9%) | A (25.7%), GAA (18.9%) | 1801 | 1628 |
| E111R | TTTGGAAATC A TATAGCCATA | Late | | AT (37.8%), ATAT (7.2%) | 23 | 23036 |
| DP146L | ATCTACACCT A AAACCATGGG | Early | A (32.4%), AA (16.4%), AAA (8.6%) | A (28.2%), AA (11.0%), AAA (5.8%) | 1400 | 1713 |
| A118R | TTTGTTTCAA A TAAACGTTGT | Late | | AT (21.3%), ATA (13.8%) | 0 | 478 |
| QP509L | TGTTGAATAT A AAAGCTTAGA | Late | | AAA (14.2%), A (6.6%), AAAA (5.5%) | 0 | 1137 |
| E120R_L | AGGTATTGTT A TAGTGATGGC | Late | | GAT (29.0%), AT (5.8%) | 60 | 112644 |
| D117L | TTTGAGATTA A TATAACTGTT | Late | | AT (33.9%) | 37 | 83180 |

| Gene | TSS sequence | Stage | Extensions in the early stage (more than 5% of total reads and at least 5 reads) | Extensions in the late stage (more than 5% of total reads and at least 5 reads) | Total reads comprising TSS in the early stage | Total reads comprising TSS in the late stage |
|---|---|---|---|---|---|---|
| I226R | TTTTGTTTTA A TATTTGCATG | Late | | AT (35.9%) | 5 | 12594 |
| B125R | TTTTTATTAA A TATGGCGGTT | Late | | AT (30.4%) | 25 | 30312 |
| Q706L | TGGGATCCTT A TAACGAGTCA | Late | | AT (29.8%) | 7 | 5562 |
| H124R | AAAAATCATT A TAAAATGAAT | Late | | AT (27.9%) | 1 | 4127 |
| J154R_L | TAAATAAACT A TAAATGAAAA | Late | | AT (26.5%) | 11 | 9144 |
| C105R | TTCCTAGAGG A GAATTAGTTT | Late | AG (33.1%) | AG (28.7%) | 2589 | 2118 |
| P1192R | TTATAGGAAT A AAAATGGAAG | Late | A (11.8%) | GATAT (15.9%), A (5.6%) | 1257 | 1708 |
| H233R | TCTAATACAC A TATTCCCTAC | Late | | AT (18.7%), ATAT (5.7%), T (5.6%) | 7 | 12452 |
| H339R | ACTAACTAAT A AATGGCCGGT | Late | | A (11.1%), AAA (7.8%) | 12 | 29191 |
| QP383R | GTAGCTTCTT A TAATTTATTC | Late | | AT (17.6%), ATAT (7.7%) | 0 | 392 |
| DP71L | TTATGTTATT A TAGGTATTAA | Late | | AT (14.8%), ATAT (5.8%) | 28 | 7961 |
| K78R | CATAATATAC A TAGAATGCCT | Late | | AT (20.7%) | 738 | 407653 |
| A104R | TTAGTTTTTT A TACAAGAATG | Late | | AT (22.0%) | 82 | 172164 |
| DP42R | TTGTAAAAAA A TATGCCTACT | Early | | AT (21.4%) | 766 | 1854 |
| K421R | TTGCCATTTT A TAGAATGTAC | Late | | AT (19.7%) | 2 | 3831 |
| C257L | ATTGTTTAAC A TAGGAGGAAA | Late | | AT (12.6%), ATAT (7.1%) | 16 | 34281 |
| L356L | AACAAAATAA A CATGCAGCCA | Late | AC (14.2%), AAA (6.6%) | AC (17.6%) | 106 | 68 |
| DP60R | AGCGGTAATA A TAATTGATAC | Early | AT (17.2%), A (5.2%) | AT (18.3%) | 58 | 71 |
| C84L | GCTGCTATTT A TATGGATCAG | Late | | AT (17.5%) | 35 | 48685 |
| B385R | TAAGGGGGGT A TAACAATGGA | Late | | GAT (18.2%) | 7 | 9460 |
| A505R | TTTGGTAAAC A AATGTTTTCT | Early | A (55.0%) | A (17.0%) | 218 | 507 |
| I73R | AAAAAGAAGT A TACTCTCCTT | Early | AT (16.1%) | AT (15.6%) | 60611 | 48749 |
| I243L_L | ATGAAAATGC A TATAGCCCGC | Late | | AT (14.7%) | 605 | 7737 |
| G1340L | TATACCGGGT A TAATGGATTT | Late | | GAT (13.0%) | 31 | 9740 |

| Gene | TSS sequence | Stage | Extensions in the early stage (more than 5% of total reads and at least 5 reads) | Extensions in the late stage (more than 5% of total reads and at least 5 reads) | Total reads comprising TSS in the early stage | Total reads comprising TSS in the late stage |
|---|---|---|---|---|---|---|
| F1055L | AATTATCAAA A TGCAAGAAAC | Late | AAAA (13.7%), AAAAA (6.2%), AAAAAA (6.2%) | AAAA (8.8%) | 306 | 421 |
| CP530R | AAATTATAAA A TAATAAGAAG | Late | | AT (13.6%) | 10 | 15575 |
| KP177R | GTTTAATATT A AAATGACAAT | Late | | A (7.2%) | 3 | 10358 |
| E146L | TTTTTATTTA A TAATGGGCGG | Late | | ATA (8.4%) | 3 | 15395 |
| O174L | TTAATATTAA A TATAAAATGT | Late | | AT (9.6%) | 83 | 3251 |
| C717R | GCTTATAAAT A TACCATGACA | Late | | ATAT (9.7%) | 2 | 2797 |
| K196R | GAAAAAAATT A AACGGTCAAA | Late | A (11.3%), AA (5.9%) | A (8.7%) | 3832 | 6061 |
| D339L | AGAATATATT A TAGATATGAT | Early | AT (8.8%) | AT (9.1%) | 3557 | 6878 |
| D129L | GCTTAATAAT A ATGGATATAA | Late | | ATAT (9.1%) | 10 | 15512 |
| V82L | TATAAAGAAT A GAGAGGGCGT | Late | | ATAT (8.0%) | 2040 | 1548 |
| EP424R | ATAAAGTAAG A AAATGTCCAA | Late | | A (6.8%) | 514 | 2226 |
| EP364R | CACGTATTTA A TATATACTAC | Late | | AT (8.7%) | 5 | 10976 |
| E184L | ATACTGTTGT A TATTATAAGA | Late | | AT (8.1%) | 183 | 433042 |
| J268L | TTTTTTTAGT A AAGACTTTTA | Early | GA (10.9%) | GA (5.9%) | 2727 | 2054 |
| H359L | CTATATATTT A TAATACAAAT | Early | AT (21.7%) | AT (7.3%) | 1767 | 12301 |
| D79L | AATTATAGTT A TATACAAATA | Late | | GAT (7.9%) | 8 | 13006 |
| I329L | TTTTAATAGT A TATACAGGAT | Late | | AT (8.2%) | 5 | 13108 |
| EP402R | ACATTATTTT A TATCATAATT | Late | | AT (7.2%) | 4 | 3191 |
| E301R | AGAATAGTAT A ATGTCTGAAG | Late | | ATAT (5.5%) | 1 | 1251 |
| A137R | ATTTATATGT A TACAACTGTA | Late | | AT (7.4%) | 438 | 1043386 |
| CP80R_E | AGTTACTTCA A GAATAACTAT | Late | | | 560 | 1142 |
| CP123L | AACGTTACTT A TATAACAAAA | Late | | AT (6.1%) | 3 | 6645 |
| CP80R_L | AAAAAATGAA A TACTAAAGTT | Late | | GAT (6.1%) | 7 | 16561 |
| I215L | AAAAAACAAA A GAGGTTCATC | Late | | | 2185 | 1018 |

| Gene | TSS sequence | Stage | Extensions in the early stage (more than 5% of total reads and at least 5 reads) | Extensions in the late stage (more than 5% of total reads and at least 5 reads) | Total reads comprising TSS in the early stage | Total reads comprising TSS in the late stage |
|---|---|---|---|---|---|---|
| B175L | GTATAAAAAT A ACTATCAAAA | Late | | | 4 | 6895 |
| B66L | GTTTTAAAGT A TATGGATATA | Late | A (14.3%) | AT (6.2%) | 7 | 8494 |
| CP312R | TTTTAATGTT A CTAGTAAAAA | Early | ACT (6.1%) | ACT (5.9%) | 46733 | 101877 |
| A224L | TATATTATAT A TAAGAATTTA | Late | | | 124 | 45570 |
| CP2475L | TTTTTTTATT A TAATGGGTAA | Late | | | 56 | 43025 |
| KP360L | TTCGGAAATA A TTATTTTGCA | Late | | | 322 | 178 |
| DP63R | AGCATTTACT A TAGTTATATT | Late | | | 2 | 2401 |
| DP148R | AAATAGTTAA A TATAGAAGTT | Late | | | 0 | 567 |
| D1133L | TTTTTTTATG A TAATATGGCG | Late | | | 1 | 1474 |
| C44L | AGATAATTAT G ACTAATAATA | Early | | | 4780 | 7359 |
| C962R | GTTTCTACAA T AATAAAATGC | Late | | | 1 | 9545 |
| D345L | GAGCTTAAAC A TATTCGCCAA | Late | AT (8.4%) | | 202 | 317 |
| F165R | TTTTTCTTCT A TATAATGGAA | Late | | | 3 | 9618 |
| EP153R | TTGTGATGGA A GACATACATA | Late | | | 4683 | 5427 |
| DP93R | TGGGATTTTT A GGTGCAACAT | Late | | | 5 | 38 |
| A240L | TTACCCATTT A TTAATCATGC | Late | | | 2379 | 248 |
| DP86L | TTCGTAAAAA A TCGCCAGTCA | Late | | | 47 | 233 |

**Supplementary Table 7. Summary of 5' extensions detected from CAGE-seq.**

| TTS Location | Strand | Cluster CAGEfightR Score | Cluster width | pTTS or npTTS | 1st ORF | 2nd ORF | 3rd ORF | TTS to 1st ORF (nt) | TTS to 2nd ORF (nt) | TTS to 3rd ORF (nt) | polyT | 1st ORF gene type | Last polyT T to TTS (nt) | polyT length |
|---|---|---|---|---|---|---|---|---|---|---|---|---|---|---|
| 47428 | + | 936 | 3 | P | K78R | K205R | - | 393 | 718 | - | Yes | Late | 1 | 5 |
| 56502 | + | 1179 | 7 | P | EP153R | EP152R | - | 334 | 798 | - | Yes | Early | 1 | 5 |
| 57850 | + | 7121 | 50 | P | EP402R | - | - | 398 | - | - | Yes | Late | 0 | 7 |
| 64452 | + | 544 | 2 | P | M448R | - | - | 598 | - | - | Yes | Early | 0 | 8 |
| 64452 | + | 544 | 2 | P | C129R | - | - | 122 | - | - | Yes | Late | 0 | 8 |
| 71325 | + | 6425 | 6 | P | C315R | - | - | 235 | - | - | Yes | Late | 3 | 9 |
| 10835 | + | 198549 | 33 | P | pNG4 * | - | - | 46 | - | - | Yes | Early | 3 | 9 |
| 92481 | + | 1102 | 3 | P | B263R | - | - | 381 | - | - | Yes | Early | 0 | 9 |
| 100168 | + | 827 | 4 | P | G1211R | - | - | 167 | - | - | Yes | Early | 0 | 9 |
| 2300 | + | 158 | 1 | P | KP86R | - | - | 295 | - | - | Yes | NC | 1 | 6 |
| 111578 | + | 12713 | 41 | P | CP312R | - | - | 153 | - | - | Yes | Early | 2 | 9 |
| 121608 | + | 47678 | 45 | P | D250R | NP868R | - | 179 | 988 | - | Yes | Early | 2 | 5 |
| 11821 | + | 257 | 1 | P | X69R | - | - | 285 | - | - | Yes | Early | 0 | 6 |
| 128801 | + | 376 | 2 | P | D205R | - | - | 610 | - | - | Yes | Late | 1 | 5 |
| 134586 | + | 157 | 1 | P | P1192R | - | - | 285 | - | - | Yes | Early | 1 | 6 |
| 12896 | + | 630293 | 13 | P | pNG3 * | - | - | 36 | - | - | Yes | Early | 3 | 9 |
| 150117 | + | 17583 | 7 | P | E165R | - | - | 10 | - | - | Yes | Early | 2 | 8 |
| 152548 | + | 19756 | 46 | P | E111R | E296R | - | 10 | 348 | - | Yes | Late | 1 | 6 |
| 155700 | + | 104 | 1 | P | I73R | - | - | 268 | - | - | Yes | Early | -2 | 7 |
| 13435 | + | 640354 | 10 | P | pNG1 | - | - | 82 | - | - | Yes | Early | 2 | 8 |
| 165126 | + | 16958 | 47 | P | DP96R | DP148R | - | 13 | 597 | - | Yes | Early | 2 | 8 |
| 166842 | + | 1074 | 3 | P | DP363R | - | - | 25 | - | - | Yes | Early | 2 | 9 |
| 167336 | + | 3081 | 3 | P | pNG6 | - | - | 151 | - | - | Yes | Early | 2 | 7 |
| 167959 | + | 1024 | 4 | P | DP60R | DP42R | - | 83 | 349 | - | Yes | Early | 2 | 9 |

| TTS Location | Strand | Cluster CAGEfightR Score | Cluster width | pTTS or npTTS | 1st ORF | 2nd ORF | 3rd ORF | TTS to 1st ORF (nt) | TTS to 2nd ORF (nt) | TTS to 3rd ORF (nt) | polyT | 1st ORF gene type | Last polyT T to TTS (nt) | polyT length |
|---|---|---|---|---|---|---|---|---|---|---|---|---|---|---|
| 168585 | + | 106 | 1 | P | DP93R | DP60R | DP42R | 353 | 709 | 975 | Yes | NC | 2 | 8 |
| 13721 | + | 914 | 3 | P | J64R | - | - | 41 | - | - | Yes | Early | 0 | 6 |
| 1722 | - | 105 | 1 | P | KP93L | KP360L | - | 148 | 481 | - | Yes | NC | 0 | 5 |
| 7150 | - | 219 | 1 | P | L270L | - | - | 129 | - | - | Yes | Early | 0 | 7 |
| 8190 | - | 23578 | 8 | P | U104L | XP124L | V82L | 29 | 440 | 992 | Yes | Early | 1 | 8 |
| 8601 | - | 28263 | 53 | P | XP124L | V82L | - | 29 | 581 | - | Yes | Early | 1 | 7 |
| 9172 | - | 1033 | 4 | P | V82L | Y118L | - | 10 | 482 | - | Yes | Early | 0 | 7 |
| 9600 | - | 31576 | 7 | P | Y118L | UP60L | - | 54 | 633 | - | Yes | Early | 2 | 7 |
| 10173 | - | 3528 | 5 | P | UP60L | - | - | 60 | - | - | Yes | Early | 1 | 6 |
| 14476 | - | 302 | 3 | P | J328L | - | - | 356 | - | - | Yes | Early | 1 | 9 |
| 15916 | - | 1031 | 4 | P | J319L | - | - | 20 | - | - | Yes | Early | 2 | 9 |
| 16911 | - | 550 | 2 | P | A125L | - | - | 146 | - | - | Yes | Early | 2 | 9 |
| 14821 | + | 8848 | 7 | P | J154R | - | - | 96 | - | - | Yes | NC | 2 | 9 |
| 29074 | - | 6214 | 45 | P | A224L | - | - | 46 | - | - | Yes | Late | -1 | 8 |
| 29827 | - | 33009 | 9 | P | A240L | - | - | 650 | - | - | Yes | Early | 1 | 8 |
| 29827 | - | 33009 | 9 | P | pNG2 | - | - | 7 | - | - | Yes | Early | 1 | 8 |
| 36205 | - | 3969 | 4 | P | A179L | - | - | 465 | - | - | Yes | Early | 1 | 7 |
| 37083 | - | 3272 | 20 | P | F317L | - | - | 816 | - | - | Yes | Late | 0 | 4 |
| 38872 | - | 18584 | 6 | P | F334L | - | - | 5 | - | - | Yes | Early | 2 | 8 |
| 42597 | - | 1631 | 3 | P | F1055L | - | - | 158 | - | - | Yes | Early | 1 | 7 |
| 49169 | - | 1014 | 3 | P | EP1242L | - | - | 221 | - | - | Yes | NC | 0 | 7 |
| 20387 | + | 9066 | 37 | P | A280R | - | - | 152 | - | - | Yes | Early | 2 | 8 |
| 64640 | - | 1566 | 9 | P | C44L | - | - | 50 | - | - | Yes | Early | 4 | 9 |
| 71074 | - | 2097 | 12 | P | C147L | C62L | - | 95 | 698 | - | Yes | Late | 0 | 7 |
| 21929 | + | 812 | 3 | P | A505R | - | - | 12 | - | - | Yes | Early | 0 | 7 |
| 79127 | - | 128 | 1 | P | B438L | - | - | 52 | - | - | Yes | Late | -6 | 4 |

| TTS Location | Strand | Cluster CAGEfightR Score | Cluster width | pTTS or npTTS | 1st ORF | 2nd ORF | 3rd ORF | TTS to 1st ORF (nt) | TTS to 2nd ORF (nt) | TTS to 3rd ORF (nt) | polyT | 1st ORF gene type | Last polyT T to TTS (nt) | polyT length |
|---|---|---|---|---|---|---|---|---|---|---|---|---|---|---|
| 79967 | - | 135 | 1 | P | B169L | - | - | 541 | - | - | Yes | Late | -1 | 7 |
| 80798 | - | 460 | 1 | P | B475L | - | - | 236 | - | - | Yes | Late | 0 | 5 |
| 23619 | + | 2851 | 5 | P | A498R | - | - | 177 | - | - | Yes | Early | 3 | 9 |
| 86645 | - | 1250 | 5 | P | B646L | - | - | 147 | - | - | Yes | Late | -4 | 4 |
| 88542 | - | 248 | 1 | P | B117L | B407L | - | 580 | 940 | - | Yes | Late | 0 | 4 |
| 27357 | + | 34000 | 12 | P | A506R | - | - | 303 | - | - | Yes | Early | 3 | 9 |
| 107914 | - | 92990 | 11 | P | CP204L | - | - | 62 | - | - | Yes | Early | 3 | 9 |
| 116645 | - | 1618 | 4 | P | NP419L | - | - | 43 | - | - | Yes | Late | 3 | 10 |
| 122047 | - | 389 | 5 | P | D79L | D339L | - | 23 | 331 | - | Yes | Late | 0 | 4 |
| 123261 | - | 209 | 1 | P | D1133L | - | - | 269 | - | - | Yes | Late | -6 | 4 |
| 127867 | - | 866 | 4 | P | D345L | - | - | 349 | - | - | Yes | Late | 0 | 8 |
| 134156 | - | 3483 | 10 | P | H359L | - | - | 174 | - | - | Yes | Early | 5 | 8 |
| 151694 | - | 577 | 3 | P | E66L | - | - | 919 | - | - | Yes | Late | 2 | 8 |
| 31635 | + | 47593 | 71 | P | A118R | - | - | 166 | - | - | Yes | Late | 0 | 4 |
| 155276 | - | 900 | 4 | P | I329L | - | - | 390 | - | - | Yes | Late | -3 | 5 |
| 156714 | - | 10059 | 6 | P | I215L | I177L | - | 249 | 934 | - | Yes | Early | 3 | 8 |
| 158788 | - | 1117 | 3 | P | DP238L | - | - | 70 | - | - | Yes | Early | 3 | 10 |
| 160652 | - | 2167 | 5 | P | DP542L | - | - | 233 | - | - | Yes | Early | 3 | 11 |
| 162534 | - | 5943 | 6 | P | DP141L | DP146L | - | 98 | 742 | - | Yes | Early | 3 | 9 |
| 163270 | - | 836 | 26 | P | DP146L | - | - | 6 | - | - | Yes | Early | 0 | 4 |
| 164478 | - | 345 | 1 | P | DP71L | - | - | 32 | - | - | Yes | Late | 0 | 5 |
| 32167 | + | 21021 | 36 | P | A151R | A118R | - | 213 | 698 | - | Yes | Early | 1 | 7 |
| 34114 | + | 2439 | 18 | P | A276R | - | - | 985 | - | - | Yes | Early | -1 | 8 |
| 38077 | + | 216 | 1 | P | A137R | - | - | 209 | - | - | Yes | Late | 0 | 8 |
| 42345 | + | 14075 | 7 | P | F778R | - | - | 105 | - | - | Yes | Early | 3 | 9 |
| 91745 | - | manual | 1 | P | B66L | G1340L | - | 350 | 556 | - | Yes | Late | 1 | 9 |

| TTS Location | Strand | Cluster CAGEfightR Score | Cluster width | pTTS or npTTS | 1st ORF | 2nd ORF | 3rd ORF | TTS to 1st ORF (nt) | TTS to 2nd ORF (nt) | TTS to 3rd ORF (nt) | polyT | 1st ORF gene type | Last polyT T to TTS (nt) | polyT length |
|---|---|---|---|---|---|---|---|---|---|---|---|---|---|---|
| 67757 | + | manual | 1 | P | C105R | C717R | - | 245 | 549 | - | Yes | Early | 0 | 5 |
| 137859 | + | manual | 1 | P | H108R | H339R | - | 105 | 443 | - | Yes | Late | -1 | 6 |
| 158130 | - | manual | 1 | P | I196L | DP238L | - | 44 | 728 | - | Yes | Late | 2 | 6 |
| 48093 | + | manual | 1 | P | K145R | K196R | - | 18 | 471 | - | Yes | Late | 0 | 9 |
| 112138 | - | manual | 1 | P | NP1450L | - | - | 115 | - | - | Yes | Early | 1 | 8 |
| 113189 | + | manual | 1 | P | O61R | - | - | 984 | - | - | Yes | Late | 1 | 9 |
| 139180 | - | manual | 1 | P | R298L | Q706L | - | 67 | 938 | - | Yes | Late | 2 | 4 |
| 46950 | + | 1531 | 9 | P | K205R | - | - | 240 | - | - | No | Early | - | - |
| 47738 | + | 425 | 4 | P | K196R | K78R | - | 116 | 703 | - | No | Early | - | - |
| 54261 | + | 115 | 1 | P | EP84R | - | - | 808 | - | - | No | Late | - | - |
| 54886 | + | 531 | 4 | P | EP424R | - | - | 123 | - | - | No | Early | - | - |
| 111205 | + | 1102 | 42 | P | CP80R | CP530R | - | 745 | 990 | - | No | Late | - | - |
| 138527 | + | 147 | 1 | P | H233R | H108R | - | 112 | 773 | - | No | Late | - | - |
| 139295 | + | 165 | 1 | P | H240R | H233R | - | 43 | 880 | - | No | Late | - | - |
| 147673 | + | 135 | 1 | P | E423R | - | - | 406 | - | - | No | Late | - | - |
| 149305 | + | 236 | 3 | P | E301R | - | - | 980 | - | - | No | Late | - | - |
| 151049 | + | 429 | 1 | P | E248R | E165R | - | 173 | 942 | - | No | Late | - | - |
| 151979 | + | 276 | 3 | P | E120R | - | - | 699 | - | - | No | Late | - | - |
| 152443 | + | 4881 | 29 | P | E296R | - | - | 243 | - | - | No | Early | - | - |
| 155328 | + | 132 | 1 | P | I226R | - | - | 930 | - | - | No | Late | - | - |
| 2174 | - | 319 | 8 | P | KP360L | - | - | 29 | - | - | No | Early | - | - |
| 5631 | - | 1061 | 12 | P | L356L | - | - | 351 | - | - | No | Early | - | - |
| 33475 | - | 135 | 1 | P | A859L | - | - | 614 | - | - | No | Late | - | - |
| 58039 | - | 1190 | 29 | P | M1249L | - | - | 678 | - | - | No | Late | - | - |
| 64055 | - | 293 | 4 | P | C84L | C44L | - | 320 | 635 | - | No | Late | - | - |
| 68231 | - | 2495 | 12 | P | C475L | - | - | 430 | - | - | No | Late | - | - |

| TTS Location | Strand | Cluster CAGEfightR Score | Cluster width | pTTS or npTTS | 1st ORF | 2nd ORF | 3rd ORF | TTS to 1st ORF (nt) | TTS to 2nd ORF (nt) | TTS to 3rd ORF (nt) | polyT | 1st ORF gene type | Last polyT T to TTS (nt) | polyT length |
|---|---|---|---|---|---|---|---|---|---|---|---|---|---|---|
| 74879 | - | 787 | 6 | P | B962L | - | - | 189 | - | - | No | Late | - | - |
| 78229 | - | 123 | 1 | P | B318L | B438L | - | 27 | 950 | - | No | Late | - | - |
| 82805 | - | 1199 | 12 | P | B602L | - | - | 868 | - | - | No | Late | - | - |
| 89177 | - | 1706 | 24 | P | B407L | - | - | 305 | - | - | No | Late | - | - |
| 99739 | - | 1691 | 7 | P | CP123L | CP2475L | - | 257 | 737 | - | No | Late | - | - |
| 121363 | - | 152 | 1 | P | D129L | D79L | - | 245 | 707 | - | No | Late | - | - |
| 144750 | - | 1914 | 13 | P | E184L | E183L | - | 64 | 662 | - | No | Late | - | - |
| 145178 | - | 2294 | 19 | P | E183L | - | - | 234 | - | - | No | Late | - | - |
| 148667 | - | 637 | 1 | P | E199L | - | - | 314 | - | - | No | Late | - | - |
| 154331 | - | 129 | 1 | P | I243L | - | - | 62 | - | - | No | NC | - | - |
| 77879 | - | manual | 1 | P | B119L | B318L | - | 37 | 377 | - | No | Late | - | - |
| 49442 | + | manual | 1 | P | K421R | - | - | 69 | - | - | No | Late | - | - |
| 111504 | + | 4130 | 19 | NP | CP312R | - | - | 79 | - | - | Yes | Early | -1 | 5 |
| 111693 | + | 650 | 2 | NP | CP312R | - | - | 268 | - | - | Yes | Early | -1 | 7 |
| 111797 | + | 615 | 4 | NP | CP312R | - | - | 372 | - | - | Yes | Early | 0 | 8 |
| 111889 | + | 466 | 13 | NP | CP312R | - | - | 464 | - | - | Yes | Early | 0 | 5 |
| 111948 | + | 13001 | 6 | NP | CP312R | - | - | 523 | - | - | Yes | Early | 1 | 8 |
| 112502 | + | 251 | 3 | NP | O61R | - | - | 297 | - | - | Yes | Late | -4 | 9 |
| 122032 | + | 737 | 2 | NP | D250R | - | - | 603 | - | - | Yes | Early | -1 | 9 |
| 122239 | + | 794 | 4 | NP | D250R | - | - | 810 | - | - | Yes | Early | 3 | 5 |
| 12323 | + | 783 | 3 | NP | X69R | - | - | 787 | - | - | Yes | Early | 2 | 8 |
| 138804 | + | 131 | 1 | NP | H233R | - | - | 389 | - | - | Yes | Late | 0 | 6 |
| 150268 | + | 110 | 1 | NP | E165R | - | - | 161 | - | - | Yes | Early | 0 | 4 |
| 152859 | + | 323 | 2 | NP | E296R | - | - | 659 | - | - | Yes | Early | 3 | 10 |
| 152859 | + | 323 | 2 | NP | E111R | - | - | 321 | - | - | Yes | Late | 3 | 10 |
| 156203 | + | 1267 | 3 | NP | I73R | - | - | 771 | - | - | Yes | Early | 1 | 5 |

| TTS Location | Strand | Cluster CAGEfightR Score | Cluster width | pTTS or npTTS | 1st ORF | 2nd ORF | 3rd ORF | TTS to 1st ORF (nt) | TTS to 2nd ORF (nt) | TTS to 3rd ORF (nt) | polyT | 1st ORF gene type | Last polyT T to TTS (nt) | polyT length |
|---|---|---|---|---|---|---|---|---|---|---|---|---|---|---|
| 160849 | + | 6111 | 6 | NP | DP63R | DP311R | - | 40 | 161 | - | Yes | Late | 3 | 7 |
| 161076 | + | 369 | 3 | NP | DP63R | DP311R | - | 267 | 388 | - | Yes | Late | 0 | 7 |
| 161333 | + | 688 | 2 | NP | DP63R | DP311R | - | 524 | 645 | - | Yes | Late | 1 | 4 |
| 165211 | + | 872 | 3 | NP | DP96R | DP148R | - | 98 | 682 | - | Yes | Early | 2 | 7 |
| 165458 | + | 495 | 2 | NP | DP96R | DP148R | - | 345 | 929 | - | Yes | Early | 0 | 7 |
| 167336 | + | 3081 | 3 | NP | DP363R | - | - | 519 | - | - | Yes | Early | 2 | 7 |
| 168380 | + | 103 | 1 | NP | DP60R | DP42R | - | 504 | 770 | - | Yes | Early | 0 | 5 |
| 168380 | + | 103 | 1 | NP | DP93R | - | - | 148 | - | - | Yes | NC | 0 | 5 |
| 2142 | - | 105 | 1 | NP | KP360L | - | - | 61 | - | - | Yes | Early | 1 | 9 |
| 9870 | - | 890 | 4 | NP | UP60L | - | - | 363 | - | - | Yes | Early | 0 | 4 |
| 28232 | - | 140 | 1 | NP | A224L | - | - | 888 | - | - | Yes | Late | 0 | 6 |
| 28271 | - | 165 | 1 | NP | A224L | - | - | 849 | - | - | Yes | Late | 0 | 7 |
| 28709 | - | 3710 | 24 | NP | A224L | - | - | 411 | - | - | Yes | Late | 2 | 7 |
| 15120 | + | 420 | 3 | NP | J154R | - | - | 395 | - | - | Yes | NC | 0 | 7 |
| 30177 | - | 5071 | 4 | NP | A240L | - | - | 300 | - | - | Yes | Early | 2 | 9 |
| 30276 | - | 4659 | 4 | NP | A240L | - | - | 201 | - | - | Yes | Early | 0 | 6 |
| 49321 | - | 102 | 1 | NP | EP1242L | - | - | 69 | - | - | Yes | NC | 0 | 5 |
| 116083 | - | 139 | 1 | NP | NP419L | - | - | 605 | - | - | Yes | Late | -1 | 6 |
| 122012 | - | 501 | 3 | NP | D79L | D339L | - | 58 | 366 | - | Yes | Late | 2 | 8 |
| 127813 | - | 171 | 1 | NP | D345L | - | - | 403 | - | - | Yes | Late | -1 | 8 |
| 156951 | - | 1262 | 17 | NP | I215L | I177L | - | 12 | 697 | - | Yes | Early | 0 | 4 |
| 162359 | - | 4407 | 5 | NP | DP141L | DP146L | - | 273 | 917 | - | Yes | Early | 1 | 6 |
| 162987 | - | 608 | 2 | NP | DP146L | - | - | 289 | - | - | Yes | Early | -1 | 4 |
| 163802 | - | 106 | 1 | NP | DP71L | - | - | 708 | - | - | Yes | Late | 2 | 6 |
| 32320 | + | 1055 | 15 | NP | A151R | A118R | - | 366 | 851 | - | Yes | Early | -3 | 5 |
| 32863 | + | 139 | 1 | NP | A151R | - | - | 909 | - | - | Yes | Early | -3 | 4 |

| TTS Location | Strand | Cluster CAGEfightR Score | Cluster width | pTTS or npTTS | 1st ORF | 2nd ORF | 3rd ORF | TTS to 1st ORF (nt) | TTS to 2nd ORF (nt) | TTS to 3rd ORF (nt) | polyT | 1st ORF gene type | Last polyT T to TTS (nt) | polyT length |
|---|---|---|---|---|---|---|---|---|---|---|---|---|---|---|
| 33414 | + | 472 | 3 | NP | A276R | - | - | 285 | - | - | Yes | Early | 0 | 5 |
| 33761 | + | 1308 | 4 | NP | A276R | - | - | 632 | - | - | Yes | Early | 1 | 8 |
| 89070 | - | manual | 1 | NP | B117L | B407L | - | 52 | 412 | - | Yes | Late | 0 | 6 |
| 47504 | + | 527 | 7 | NP | K78R | K205R | - | 469 | 794 | - | No | Late | - | - |
| 47594 | + | 5235 | 73 | NP | K78R | K205R | - | 559 | 884 | - | No | Late | - | - |
| 47942 | + | 635 | 15 | NP | K196R | K78R | - | 320 | 907 | - | No | Early | - | - |
| 54225 | + | 103 | 1 | NP | EP84R | - | - | 772 | - | - | No | Late | - | - |
| 54937 | + | 139 | 1 | NP | EP424R | - | - | 174 | - | - | No | Early | - | - |
| 55003 | + | 105 | 1 | NP | EP424R | - | - | 240 | - | - | No | Early | - | - |
| 55041 | + | 218 | 1 | NP | EP424R | - | - | 278 | - | - | No | Early | - | - |
| 55109 | + | 121 | 1 | NP | EP424R | - | - | 346 | - | - | No | Early | - | - |
| 57107 | + | 614 | 8 | NP | EP153R | - | - | 939 | - | - | No | Early | - | - |
| 57524 | + | 1916 | 36 | NP | EP402R | - | - | 72 | - | - | No | Late | - | - |
| 58425 | + | 160 | 1 | NP | EP402R | - | - | 973 | - | - | No | Late | - | - |
| 64194 | + | 121 | 1 | NP | M448R | - | - | 340 | - | - | No | Early | - | - |
| 110639 | + | 659 | 8 | NP | CP80R | CP530R | - | 179 | 424 | - | No | Late | - | - |
| 112369 | + | 906 | 27 | NP | O61R | CP312R | - | 164 | 944 | - | No | Late | - | - |
| 112536 | + | 137 | 1 | NP | O61R | - | - | 331 | - | - | No | Late | - | - |
| 112620 | + | 409 | 12 | NP | O61R | - | - | 415 | - | - | No | Late | - | - |
| 112749 | + | 124 | 1 | NP | O61R | - | - | 544 | - | - | No | Late | - | - |
| 112802 | + | 276 | 2 | NP | O61R | - | - | 597 | - | - | No | Late | - | - |
| 112854 | + | 1675 | 13 | NP | O61R | - | - | 649 | - | - | No | Late | - | - |
| 137727 | + | 266 | 1 | NP | H339R | - | - | 311 | - | - | No | Late | - | - |
| 139408 | + | 120 | 1 | NP | H240R | H233R | - | 156 | 993 | - | No | Late | - | - |
| 139595 | + | 104 | 1 | NP | H240R | - | - | 343 | - | - | No | Late | - | - |
| 139641 | + | 146 | 1 | NP | H240R | - | - | 389 | - | - | No | Late | - | - |

| TTS Location | Strand | Cluster CAGEfightR Score | Cluster width | pTTS or npTTS | 1st ORF | 2nd ORF | 3rd ORF | TTS to 1st ORF (nt) | TTS to 2nd ORF (nt) | TTS to 3rd ORF (nt) | polyT | 1st ORF gene type | Last polyT T to TTS (nt) | polyT length |
|---|---|---|---|---|---|---|---|---|---|---|---|---|---|---|
| 139697 | + | 167 | 1 | NP | H240R | - | - | 445 | - | - | No | Late | - | - |
| 139733 | + | 6217 | 25 | NP | H240R | - | - | 481 | - | - | No | Late | - | - |
| 140009 | + | 349 | 5 | NP | H240R | - | - | 757 | - | - | No | Late | - | - |
| 8935 | - | 120 | 1 | NP | V82L | Y118L | - | 247 | 719 | - | No | Early | - | - |
| 9763 | - | 146 | 1 | NP | UP60L | - | - | 470 | - | - | No | Early | - | - |
| 30407 | - | 226 | 1 | NP | A240L | - | - | 70 | - | - | No | Early | - | - |
| 38384 | - | 100 | 1 | NP | F334L | - | - | 493 | - | - | No | Early | - | - |
| 57914 | - | 1045 | 11 | NP | M1249L | - | - | 803 | - | - | No | Late | - | - |
| 58089 | - | 1013 | 3 | NP | M1249L | - | - | 628 | - | - | No | Late | - | - |
| 63642 | - | 443 | 5 | NP | C84L | - | - | 733 | - | - | No | Late | - | - |
| 63745 | - | 335 | 1 | NP | C84L | C44L | - | 630 | 945 | - | No | Late | - | - |
| 64412 | - | 351 | 1 | NP | C44L | - | - | 278 | - | - | No | Early | - | - |
| 68277 | - | 310 | 1 | NP | C475L | - | - | 384 | - | - | No | Late | - | - |
| 74993 | - | 595 | 5 | NP | B962L | - | - | 75 | - | - | No | Late | - | - |
| 78306 | - | 134 | 1 | NP | B438L | - | - | 873 | - | - | No | Late | - | - |
| 82899 | - | 297 | 4 | NP | B602L | - | - | 774 | - | - | No | Late | - | - |
| 86260 | - | 227 | 4 | NP | B646L | - | - | 532 | - | - | No | Late | - | - |
| 86532 | - | 574 | 1 | NP | B646L | - | - | 260 | - | - | No | Late | - | - |
| 120653 | - | 106 | 1 | NP | D129L | - | - | 955 | - | - | No | Late | - | - |
| 127337 | - | 602 | 27 | NP | D345L | - | - | 879 | - | - | No | Late | - | - |
| 127977 | - | 135 | 1 | NP | D345L | - | - | 239 | - | - | No | Late | - | - |
| 144645 | - | 1111 | 2 | NP | E184L | E183L | - | 169 | 767 | - | No | Late | - | - |
| 31558 | + | 1838 | 27 | NP | A118R | - | - | 89 | - | - | No | Late | - | - |
| 162601 | - | 1461 | 23 | NP | DP141L | DP146L | - | 31 | 675 | - | No | Early | - | - |
| 163872 | - | 176 | 1 | NP | DP71L | - | - | 638 | - | - | No | Late | - | - |
| 32063 | + | 4600 | 36 | NP | A151R | A118R | - | 109 | 594 | - | No | Early | - | - |

| TTS Location | Strand | Cluster CAGEfightR Score | Cluster width | pTTS or npTTS | 1st ORF | 2nd ORF | 3rd ORF | TTS to 1st ORF (nt) | TTS to 2nd ORF (nt) | TTS to 3rd ORF (nt) | polyT | 1st ORF gene type | Last polyT T to TTS (nt) | polyT length |
|---|---|---|---|---|---|---|---|---|---|---|---|---|---|---|
| 32722 | + | 134 | 1 | NP | A151R | - | - | 768 | - | - | No | Early | - | - |
| 32826 | + | 392 | 4 | NP | A151R | - | - | 872 | - | - | No | Early | - | - |
| 32919 | + | 5253 | 38 | NP | A151R | - | - | 965 | - | - | No | Early | - | - |
| 33132 | + | 146 | 1 | NP | A276R | - | - | 3 | - | - | No | Early | - | - |
| 38394 | + | 254 | 1 | NP | A137R | - | - | 526 | - | - | No | Late | - | - |
| 154580 | + | manual | 1 | NP | I226R | - | - | 182 | - | - | No | Late | - | - |

**Supplementary Table 8. Summary of 212 TTS clusters detected with CAGEfightR analysis and polyA filtering (see Methods).** Manual refers to manually annotated clusters. TTS location is the highest point of each cluster. * clusters from pNG3 and pNG4 were filtered out from polyA filtering but added back due to high RNA-seq agreement with termination signals.
